# Ssl2/TFIIH function in Transcription Start Site Scanning by RNA Polymerase II in *Saccharomyces cerevisiae*

**DOI:** 10.1101/2021.05.05.442816

**Authors:** Tingting Zhao, Irina O. Vvedenskaya, William K.M. Lai, Shrabani Basu, B. Franklin Pugh, Bryce E. Nickels, Craig D. Kaplan

## Abstract

In *Saccharomyces cerevisiae*, RNA Polymerase II (Pol II) selects transcription start sites (TSS) by a unidirectional scanning process. During scanning, a preinitiation complex (PIC) assembled at an upstream core promoter initiates at select positions within a window ∼40-120 basepairs downstream. Several lines of evidence indicate that Ssl2, the yeast homolog of XPB and an essential and conserved subunit of the general transcription factor (GTF) TFIIH, drives scanning through its DNA-dependent ATPase activity, therefore potentially controlling both scanning rate and scanning extent (processivity). To address questions of how Ssl2 functions in promoter scanning and interacts with other initiation activities, we leveraged distinct initiation-sensitive reporters to identify novel *ssl2* alleles. These *ssl2* alleles, many of which alter residues conserved from yeast to human, confer either upstream or downstream TSS shifts at the model promoter *ADH1* and genome-wide. Specifically, tested *ssl2* alleles alter TSS selection by increasing or narrowing the distribution of TSSs used at individual promoters. Genetic interactions of *ssl2* alleles with other initiation factors are consistent with *ssl2* allele classes functioning through increasing or decreasing scanning processivity but not necessarily scanning rate. These alleles underpin a residue interaction network that likely modulates Ssl2 activity and TFIIH function in promoter scanning. We propose that the outcome of promoter scanning is determined by two functional networks, the first being Pol II activity and factors that modulate it to determine initiation efficiency within a scanning window, and the second being Ssl2/TFIIH and factors that modulate scanning processivity to determine the width of the scanning widow.

## INTRODUCTION

Transcription of eukaryotic protein-coding genes is carried out by RNA polymerase II (Pol II) in three sequential steps: initiation, elongation and termination^1^. Accurate initiation requires minimally the assistance of five general transcription factors (GTFs) TFIIB, TFIID, TFIIE, TFIIF and TFIIH, which together with Pol II, comprise the basal transcription machinery. At the beginning of transcription, this machinery assembles at a defined DNA region for each transcript called a promoter, melts the double-stranded DNA yielding a region of unwound DNA forming a “transcription bubble”. Within this bubble a position or positions will be identified to serve as transcription start sites (TSS). Initial promoter melting appears to occur stereotypically ∼20-25 nucleotides downstream of promoter elements such as the TATA box across eukaryotes, though most promoters lack a TATA box or other strong sequence signature of promoter elements. During initiation, the process of TSS selection determines the identity and distribution of transcript isoforms that differ by their 5’ ends. Differences in 5’ UTR can alter transcript properties such as translation efficiency or transcript stability through differences in sequence or RNA secondary structure^2-8^. Furthermore, in conjunction with activators and coactivators, the efficiency of the initiation process will also establish mRNA synthesis rates. How TSS selection is governed by these factors is not well understood for the majority of eukaryotic promoters that utilize multiple TSSs.

Transcription initiation by *S. cerevisiae* Pol II has been the subject of extensive analysis both in vivo and in vitro, and thus provides a powerful model for system for mechanistic studies of TSS selection. TSS selection by *S. cerevisiae* Pol II occurs over a range of positions located ∼40-120 bp downstream of the core promoter region. Numerous lines of evidence suggest TSS selection by *S. cerevisiae* Pol II involves a unidirectional scanning mechanism in which the preinitiation complex (PIC) assembles at an upstream core promoter and interrogates consecutive downstream positions for usable TSSs^9-13^. TFIIH is proposed to drive Pol II scanning through ATP-dependent DNA translocase activity^12, 14-17^. An optical-tweezer-based single molecule analysis of reconstituted *S. cerevisiae* PICs indicated that an ATP/dATP-induced activity within the PIC causes shortening of the distance between upstream and downstream DNA^17^. This shortened distance approximates the distance downstream from TATA-elements where TSSs are positioned in yeast (40-120nt)^18^ and suggests downstream DNA movement and compaction during promoter scanning by the PIC. Separately, a magnetic-tweezer-based single molecule assay suggested that an initial melted region of 6nt (a “bubble”) is the direct consequence of TFIIH’s ATPase activity^15^. Due to inability of magnetic-tweezers to detect DNA compaction in the particular setup used, how the Pol II machinery reaches downstream TSSs, whether through generation of a large bubble or translocation of a small bubble was not clear. Nevertheless, both studies agree that an ATP-dependent PIC activity for promoter opening is likely Ssl2 within TFIIH, which has been demonstrated as a DNA translocase within purified TFIIH in vitro^16^.

In support of Ssl2/TFIIH’s role in movement of downstream DNA toward the PIC, *ssl2* mutants have been identified as altering TSS selection at *ADH1* and showed genetic interactions with *sua7* (TFIIB) mutants^19^. Specifically, the identified *ssl2* mutants shifted TSSs upstream at *ADH1*. Polar shifts in TSSs distributions have been observed in mutants within Pol II, the GTFs TFIIB and TFIIF, and the PIC cofactor Sub1^10, 13, 19-38^. In a promoter scanning initiation mechanism, altering initiation efficiency is predicted to alter TSS distributions in a polar fashion when initiation efficiency increases or decreases. We have recently observed polar (directional) shifts at essentially all promoters examined across the genome in yeast for tested Pol II and GTF mutants, as predicted for scanning operating universally across promoter classes^13^. We found that hyperactive Pol II catalytic mutants shifted TSSs distributions upstream at promoters genome-wide, consistent with a higher probability of initiation at every TSS, and thus, initiation happening on average *earlier* in the scanning process. Conversely, hypoactive Pol II catalytic mutants shift TSS distributions downstream at promoters genome-wide, consistent with initiation happening *later* in the scanning process. Our previous data on genetic interactions between Pol II and TFIIF or TFIIB support the idea that these mutations altered initiation additively^39^, consistent with their acting in the same pathway during scanning. Tested TFIIB mutants appeared to reduce initiation efficiency while tested TFIIF mutants appeared to increase initiation efficiency. Consistent with this idea, a double mutant between TFIIF and a hyperactive Pol II mutant had stronger effects on TSS shifts than either mutant alone across all promoters. Pol II mutants are proposed to control initiation efficiency because active site residues important for catalytic activity alter TSS distributions correlating with the strengths of their catalytic defects in RNA chain elongation. Initiation by promoter scanning should be dependent both on Pol II catalytic rate together and by whichever factors control the actual scanning step, *i*.*e*. presumptively the rate and processivity of TFIIH as the putative scanning translocase. Therefore, to understand how promoter scanning works, it is critical to understand how TFIIH contributes and how its activity is regulated within the PIC.

We have described the scanning process previously using a “Shooting Gallery” analogy^13, 40^. In this model, initiation is controlled by the rate (TFIIH’s translocase activity) in which a “target” (TSS) passes the “line of fire” (the Pol II active site) along with the rate of firing (Pol II catalytic activity) and the size of the target (innate sequence strength). Together, the cooperation and competition between these rates determines the probability a target is hit (initiation happening). Alteration of enzymatic activities supporting initiation, either the Pol II active site or TFIIH translocation should have predictable effects on overall TSS distributions when initiation proceeds by scanning. In addition to the TFIIH translocation rate, it is predicted that processivity of TFIIH DNA translocation should strongly modulate scanning, but in ways distinct from controlling innate initiating efficiency. Here, TFIIH processivity would control the probability that a TSS could be reached during a scanning foray from a core promoter, which appears to be facilitated by Ssl2’s translocase activity^17^. Optical tweezer experiments are consistent with TFIIH within the PIC having median processivity on the order of ∼90 basepairs, consistent with the average distance of TSSs downstream of yeast TATA boxes. However, purified holo-TFIIH from yeast has much reduced measured processivity and human TFIIH has essentially none^14^. Given that TFIIH activity is predicted to be altered extensively by cofactor interactions in both transcription and nucleotide excision repair (NER), it will be important to understand TFIIH functions within the PIC in vivo. How alterations to Ssl2/TFIIH translocase activity control TSS distributions has not been extensively investigated.

To test if distinct alterations to Ssl2 function have broad effects on promoter scanning and TSS selection, we used existing and newly identified *ssl2* alleles to examine their effect on TSS distributions genome-wide. Our novel alleles were identified through use of genetic reporters we have developed and found to be sensitive to different kinds of initiation defects^41^. We have found that *ssl2* alleles affect TSS distributions for the majority of promoters in yeast and for all promoter classes. Furthermore, we find that *ssl2* alleles alter TSS selection distinctly from how changes to the Pol II active site alter TSS selection, consistent with a distinct role for Ssl2 in promoter scanning. *ssl2* alleles appear to extend or truncate scanning windows at promoters genome-wide, consistent with increase or decrease in the processivity of scanning. Scanning-window truncating alleles map throughout the Ssl2 structure, consistent with a hypothetical loss of functions in Ssl2 DNA translocase enzymatic activity that lead to decreased TFIIH processivity. Conversely, scanning-window extending *ssl2* alleles are much more localized within the Ssl2 N-terminus, including conserved residues within regions that connect Ssl2 helicase domains to the TFIIH component and regulator of Ssl2 activity Tfb2. Our alleles are consistent with alteration to an Ssl2 regulatory domain resulting in modulated translocase activity or TFIIH processivity. We further test our model for initiation by promoter scanning through examination of genetic interactions of initiation-altering Pol II/GTF alleles and *ssl2* alleles. The genetic interactions between Pol II/GTF and Ssl2 alleles support the idea of two major networks controlling TSS selection in *S. cerevisiae*. One network shapes TSS distributions through affecting initiation efficiency, represented by Pol II, TFIIB and TFIIF functions. A second network appears to alter TSS distributions through regulating TFIIH’s processivity, and includes Ssl2, Sub1, and potentially TFIIF.

## RESULTS

### Existing *SSL2* alleles show transcription-dependent growth phenotypes and distinct TSS usage patterns

To understand how TSSs are identified by promoter scanning and the potential roles for TFIIH, we first examined previously identified *ssl2* mutants^19, 42-44^ for transcription-related phenotypes that we have demonstrated are predictive of specific initiation defects **(Figure 1A)**. Two such phenotypes relate to altered initiation at the *IMD2* gene, whose promoter is regulated by a transcription start site switch **(Figure 1A, *IMD2, imd2***Δ***::HIS3*)**^45, 46^. We have previously shown that tested mutants that shift TSSs upstream due to altered promoter scanning result in an inability to express a functional *IMD2* transcript, causing sensitivity to the IMPDH inhibitor mycophenolic acid (MPA)^13, 39, 41, 47^ **(Figure 1A, *IMD2*)**. In the presence of GTP starvation induced by MPA, WT strains shift start site usage at *IMD2* from TATA-proximal GTP-initiating TSSs to a downstream ATP-initiating TSS. This shift results in a functional *IMD2* transcript that is required for yeast to survive MPA. Catalytically hyperactive Pol II mutants (termed GOF for “Gain-of-Function”) do not shift TSS usage to the downstream functional *IMD2* TSS but instead shift to intermediately positioned non-functional upstream sites, rendering yeast sensitive to MPA (MPA^S^) ^41^. Pol II GOF mutants with MPA^S^ phenotypes and a *tfg2* MPA^S^ mutant were additionally found to shift TSS distributions upstream at *ADH1* and subsequently genome-wide^13, 47, 48^. Correlation between strength of MPA-sensitive phenotypes and quantitative upstream TSS shifts at *ADH1* and genome-wide suggest that MPA-sensitivity is a strong predictor for upstream-shifting TSS mutants. Conversely, we found previously that mutants shifting TSSs downstream (reduced catalytic activity loss of function “LOF” Pol II mutants) constitutively express *IMD2* in the absence of using MPA as inducer^47^. Pol II TSS downstream shifting phenotypes at *IMD2* can be detected using a reporter allele where *IMD2* is replaced by *HIS3*, placing *HIS3* under control of *IMD2* promoter and TSS selection **(Figure 1A, *imd2***Δ***::HIS3*)** ^41^. Indeed, these same LOF Pol II mutants shift TSS distributions downstream at *ADH1* and genome-wide^13, 39, 47^. These previous results suggest we have phenotypes predictive of alterations to promoter scanning in both directions and can form the basis of a system to characterize mutants across the genome for effects on TSS selection by promoter scanning.

**Figure 1:**
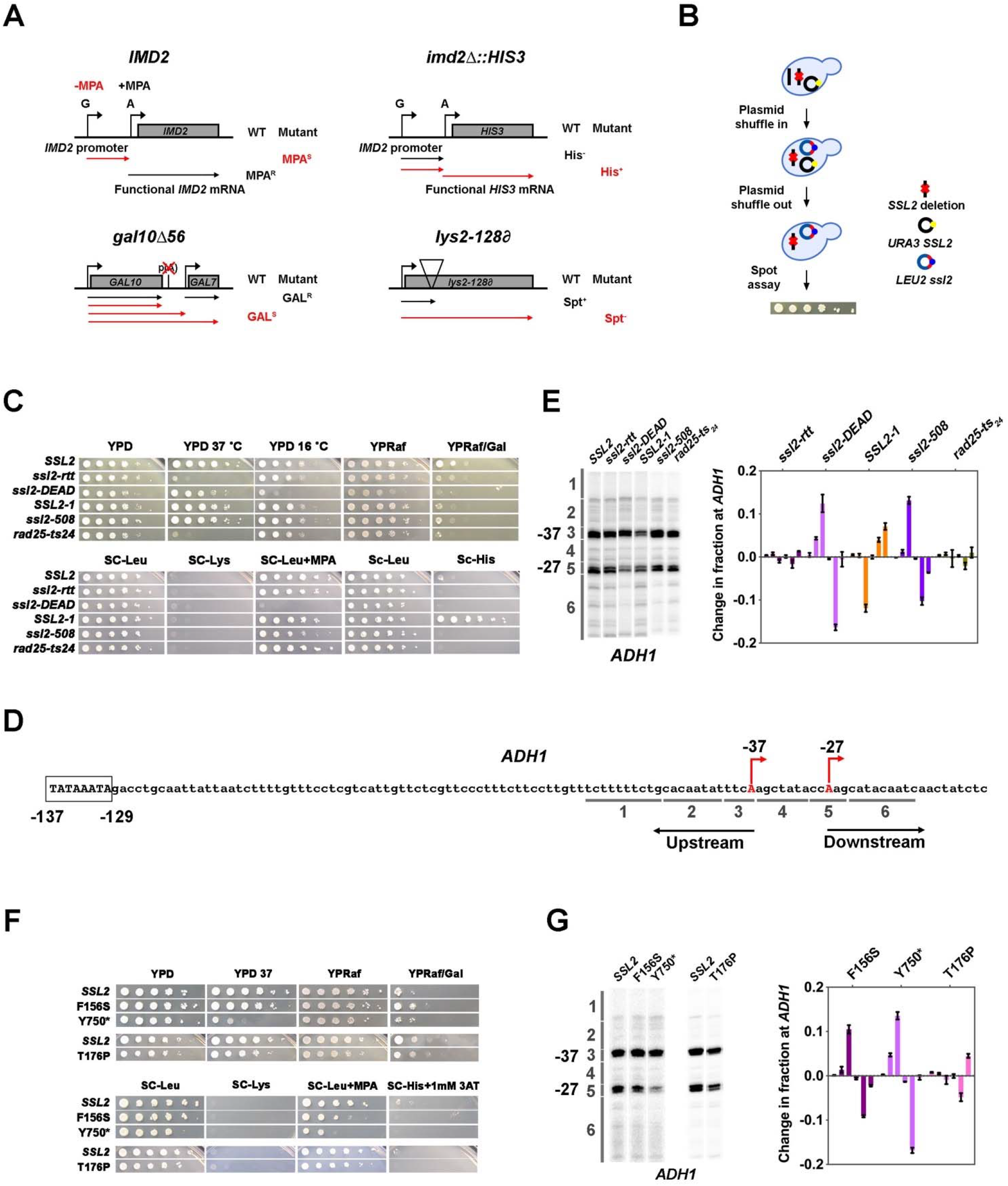
Genetic screening identifies novel *ssl2* alleles with transcriptional defects. **(A)** Schematics illustrating transcriptional phenotypes of four transcriptional phenotypes utilized in this study (*IMD2, imd2*Δ*::HIS3, gal10*Δ*56, lys2-128*∂). ***IMD2***, In GTP replete conditions, *IMD2* transcription initiates at upstream TSSs utilizing GTP as the first nucleotide. These are non-functional due to the presence of a terminator prior to the *IMD2* coding sequence. Upon GTP starvation induced by the drug MPA, initiation shifts to a downstream ATP-initiated TSS, enabling functional *IMD2* expression, conferring resistance to MPA. Inability to shift to the downstream *IMD2* TSS in the presence of MPA leads to MPA sensitivity (MPA^S^), commonly found in mutants that shift TSSs upstream. ***imd2***Δ***::HIS3***, The *IMD2* ORF is replaced by the *HIS3* ORF creating a transcriptional fusion between the *IMD2* promoter and *HIS3*. If the downstream *IMD2* TSS is used constitutively, *HIS3* mRNA production to supports growth on medium lacking histidine (SC-His). Inappropriate use of the downstream *IMD2* start site by downstream TSS shifting mutants in the absence of MPA can be detected by a His+ phenotype in cells with the *imd2*Δ*::HIS3* reporter. ***gal10***Δ***56***, Deletion of *GAL10* polyadenylation signal at p(A) site interferes with *GAL10* 3’-end formation and *GAL7* initiation, resulting in galactose toxicity^77^. Transcriptional mutants that increase *GAL10* 3’-end formation or *GAL7* initiation allow suppression of galactose toxicity and display galactose resistance (GAL^R^). ***lys2-128***∂, Insertion of a Ty transposon ∂ element into *LYS2* causes premature termination of Pol II initiating at *LYS2*, resulting in lysine auxotrophy. Certain mutants allow expression of *LYS2* from a cryptic site within the Ty d element and allow yeast growth on medium lacking lysine (SC-Lys), conferring the “Suppressor of Ty” (Spt^-^) phenotype. **(B)** Schematic illustrating the plasmid shuffle assay to examine *ssl2* mutant phenotypes (see Materials and Methods). **(C)** Transcription-related growth phenotypes of five classical *ssl2* alleles. All spot dilutions shown throughout manuscript are representative of at least two independent transformants **(D)** Schematic illustrating the TSS region of *S. cerevisiae ADH1*. The *ADH1* promoter contains two commonly used TSSs (red letters), which are 37nt (−37) and 27nt (−27) upstream from the translational start codon. For quantification of changes to *ADH1* TSS distribution by primer extension, *ADH1* TSS usage is separated into six bins. **(E) Left panel**, primer extension-detected TSS usage of the WT and five existing *ssl2* mutants at *ADH1*. **Right panel**, quantitative analysis of five classical *ssl2* alleles at *ADH1*. Average of ≥ 3 biological replicates +/- standard deviation are shown. **(F)** Transcription-related growth phenotypes of *ssl2* alleles homologous to human disease alleles of XPB. **(G)** Primer extension detection of TSS usage at *ADH1* for alleles shown in (F). Average of ≥ 3 biological replicates +/- standard deviation are shown.

We used site-directed mutagenesis and plasmid shuffling to recreate and phenotype five previously described *ssl2* mutants, reasoning that this would allow us a first glimpse at the effects and potential diversity present in these classic alleles. These alleles are *ssl2-rtt* (*ssl2* E556K)^44^, *ssl2-DEAD* (*ssl2* V490A/H491D) and *SSL2-1* (*ssl2* W427L)^42^, *ssl2-508* (*ssl2* H508R)^19^, and *rad25-ts24* (*ssl2* V552I/E556K)^43^ **(Figure 1B,1C)**. Analysis of these five *ssl2* mutants showed phenotypes consistent with altered TSS selection **(Figure 1C)**. First, *ssl2-DEAD* and *ssl2-508* exhibited strong and weak MPA^S^ phenotypes, respectively. These results were consistent with prior analysis of initiation at *ADH1* in these mutants^19^ suggesting that our genetic phenotypes using *IMD2* are predictive of potentially global alterations to TSS selection. *SSL2-1* exhibited a His^+^ phenotype, which is predictive of downstream shifts in TSS usage and consistent with its identification as a dominant mutant bypassing an inhibitory stem loop in the *his4-316* mRNA^42^. We now can rationalize the original “Suppressor of Stem-loop (SSL) phenotype of *SSL2-1* as usage of TSS downstream or within the inhibitory 5’ stem loop *his4-316* sequence (though not apparent in Gulyas *et al*^42^). *ssl2-rtt* and *rad25-ts24* show temperature-sensitive phenotypes as expected; however, assayed transcription-related plate phenotypes were not observed. Due to the absence of His^+^ or MPA^S^ phenotypes, we predicted that *ssl2-rtt* or *rad25-ts24* alleles would not shift TSS usage. Notably, there was no *lys2-128*∂ Spt^-^ phenotype observed among these five existing *ssl2* alleles, in contrast to our previous observation of an Spt^-^ phenotype in a subset of MPA^S^ Pol II TSS alleles^49, 50^, our first indication that *ssl2* alleles may alter TSS selection in a distinct fashion from Pol II mutants.

To quantitatively examine TSS usage of these *ssl2* mutants, we chose *ADH1* initiation. *ADH1* has been widely used as a model gene for TSS studies in *S. cerevisiae*. It contains two major TSSs that are 27 and 37 nucleotides upstream of the start codon **(Figure 1D)**. Using primer extension, transcription products of these two TSSs appear as bands of differing mobility on denaturing polyacrylamide gels **(Figure 1 – Figure supplement 1A, left panel WT)**. Other TSSs positions show minor usage. In most studies, the two major starts are compared qualitatively, but such comparisons miss meaningful alterations that may tell us about initiation mechanisms. To establish that genetic phenotypes using *IMD2* correlate with altered TSS selection elsewhere in the genome our quantitative analysis of the *ADH1* promoter divides TSSs observed by primer extension into six bins from upstream to downstream, in which bin three and five each contain the TSS for one of the major *ADH1* mRNA isoforms **(Figure 1D, and Figure 1 – Figure supplement 1A left panel WT)**. In order to compare a mutant’s TSS distribution to that of WT, distributions are normalized, and the WT distribution is subtracted from tested mutant distributions bin by bin **(Figure 1 – Figure supplement 1A, middle and right panel)**. Negative or positive values indicate that the mutant has relatively lower or higher usage for TSSs in that particular bin, respectively **(Figure 1 – Figure supplement 1A, right panel)**. For example, the Pol II GOF allele E1103G increases relative TSS usage at upstream minor sites (TSS bin two) and decreases relative TSS usage at the downstream major site (TSS bin four) **(Figure 1 – Figure supplement 1A, E1103G)**. Because of the dramatic effect of E1103G on TSS usage, the change of TSS usage can be easily visually detected on a primer extension gel **(Figure 1 – Figure supplement 1A, left panel E1103G)**. However, less visually obvious but highly reproducible phenotypes are detected upon quantification **(Figure 1 – Figure supplement 1A, right panel E1103G-WT)**. As predicted from plate phenotypes observed and a previous study^19^, *ssl2-DEAD* and *ssl2-508* showed upstream shifts in their *ADH1* TSS distributions **(Figure 1C and 1E)**. However, we observed that these *ssl2* alleles were quantitatively distinct in the amount of upstream shifting from Pol II and GTF alleles with comparable MPA sensitivities. *ssl2-DEAD* and *ssl2-508* appeared primarily to reduce downstream TSS usage (loss in bin 5 and gain in bin 3) whereas E1103G has its largest gain in bin 2 **(compare Figure 1 – Figure supplement 1A E1103G, and Figure 1E *ssl2-DEAD* and *ssl2-508*)**. Consistent with the prediction based on its *imd2*Δ*::HIS3* phenotype, the His^+^ *SSL2-1* mutant shifted the overall TSS distribution downstream through increased relative downstream TSS utilization **(Figure 1C and 1E)**. Previously, it had been concluded that *SSL2-1* had no obvious effects on TSSs distribution when comparing usage of just the two major starts^19^. Two other mutants, *ssl2-rtt* and *rad25-ts24*, had no obvious effects on TSS utilization at *ADH1*, consistent with their lack of plate phenotypes **(Figure 1C and 1E)**.

We additionally constructed and tested human disease-related XPB mutations^51-54^ in the yeast *SSL2* system, together with mutations in the ultra-conserved Arginine-Glutamic Acid-Aspartic Acid (RED) motif. As human disease-related alleles are many times found in conserved residues, we reasoned that some may have effects on Ssl2 biochemistry detectable in our sensitive system. We examined four human disease-related mutants that confer distinct inherited autosomal recessive disorders xeroderma pigmentosum (XP), trichothiodystrophy (TTD), and Cockayne syndrome (CS). Of these Q592*, which creates a C-terminally truncated Ssl2 protein^51^, confers lethality **(Figure 1 – Figure supplement 1B)**, and T176P^52-54^ confers little if any growth defects and no MPA^S^ or His^+^ phenotypes **(Figure 1F**), consistent with unaltered TSS usage **(Figure 1G)**. In contrast, F156S^52, 54, 55^ conferred a mild MPA^S^ phenotype and shifted TSS distribution upstream at *ADH1* **(Figure 1G)**. Mutant Y750*, which mimics a disease related C-terminally truncated protein^19, 56, 57^, shows a mild to moderate level MPA^S^ phenotype **(Figure 1F)** and shifts TSS distribution upstream at *ADH1* **(Figure 1G)**. The lethal phenotypes of RED motif substitutions in *ssl2* revealed their essential roles in *S. cerevisiae*. These results suggest that a subset of human disease alleles can alter TFIIH functions when placed in the yeast system.

### Novel *ssl2* mutants with transcriptional defects

Our establishment of a genetic system sensitive to *ssl2* mutant mediated initiation defects allowed us to obtain a broad set of alleles for the study of Ssl2 function in promoter scanning by large-scale genetic screens (see Methods). Our genetic screening has identified at least two phenotypically distinguishable classes of *ssl2* alleles: the first class is putatively defective for the induction of the *IMD2* gene, resulting in sensitivity to MPA, the second class confers constitutive expression of *imd2*Δ*::HIS3*, resulting in a His^+^ phenotype **(Figure 1A, Figure 2 – Figure supplement 1)**. Other transcription-related or conditional phenotypes, Spt^-58^, suppression of *gal10*Δ*56* (Gal^R^)^59, 60^ or temperature sensitivity (Tsm^-^), were observed in distinct subsets of alleles of the two major classes **(Figure 1A, Figure 2 – Figure supplement 1)**. The Spt^-^ phenotype reporter used in our strains, *lys2-128*∂, detects activation of a TSS within a Ty1 ∂ element at the 5’ end of the *LYS2* gene^58^. Importantly, a subset of TSS-shifting alleles show the Spt^-^ phenotype and it is useful to further classify identified *ssl2* alleles. We observed that Spt^-^ and His^+^ phenotypes were dominant for tested alleles (not shown), suggestive of possible gain of function; in contrast, there were no dominant alleles found among MPA^S^ mutants, consistent with either recessive loss-of-function mutations or the nature of the phenotype (sensitivity) or both. We then asked if TSS usage at *ADH1* was altered predictably in the mutant classes as we observed for existing *ssl2* mutants and previously studied Pol II and other GTF mutants. We find that the two major classes of *ssl2* mutants exhibited predicted TSS shifts, with all tested MPA^S^ alleles shifting TSS usage upstream and all His^+^ alleles shifting TSS usage downstream **(Figure 2 – Figure supplement 2)** validating our genetic method for identifying *ssl2* alleles conferring altered initiation properties.

We examined our substitution mutants in the context of the structure of Ssl2 within the yeast preinitiation complex as determined by the Cramer lab^61^ to understand how these alleles relate to PIC architecture and interactions. Substitutions causing MPA^S^ phenotypes and upstream TSS shifts alter amino acids distributed across the protein, with a large number in conserved domains and highly conserved residues **(Figure 2A, Figure 2 – Figure supplement 3)**. In contrast, mutations related to His^+^ phenotypes, while also generally conserved, alter amino acids clustered N-terminally, within a domain predicted^62, 63^ and now observed to be homologous to Ssl2’s interaction partner Tfb2^61, 64^ **(Figure 2A)**. This domain was first fully observed in a human core TFIIH structure (PDB 6NMI)^64^ but has now been observed in just published higher resolution PIC structures^61^ **(Figure 2B)**. This key visualization of the conserved Ssl2 N-terminus allows placement of most *ssl2* mutant residues identified as His^+^ or both His^+^ and Spt^-^ **(Figure 2C, D)**. In addition, the Spt^-^ phenotype, not previously observed for classical *ssl2* alleles, was found exclusively within a subset of stronger downstream TSS shifting His^+^ mutants **(Figure 2A)**. The coincidence between His^+^ and Spt^-^ phenotypes for *ssl2* mutants is in contrast to what is observed for Pol II mutants. In our previous studies of Pol II mutants, the Spt^-^ phenotype was observed in Pol II catalytic center substitutions, overlapping with MPA^S^ to a large extent, and tightly linked with increased Pol II activity^47, 49, 50^. Pol II alleles with both Spt^-^ and MPA^S^ phenotypes increased the efficiency of TSS usage resulting in upstream shifts to TSS distributions across promoters in vivo. However, in our identified *ssl2* alleles, none of the Spt^-^ mutants also conferred MPA^S^ **(Figure 2A)**. These observations together are consistent with distinct effects on structure and function in *ssl2* mutant classes.

**Figure 2:**
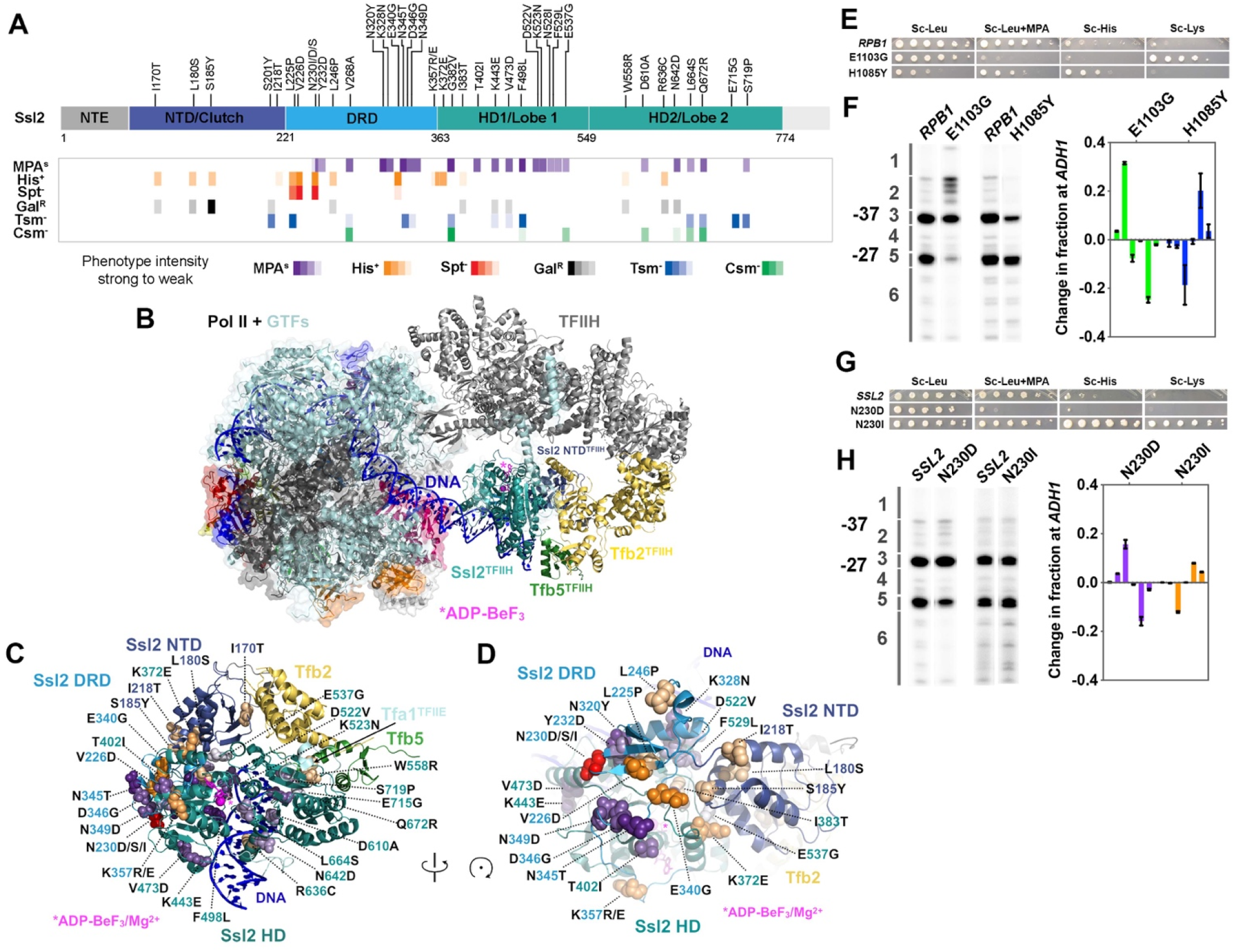
*ssl2* alleles have distinct behavior from Pol II and other GTFs alleles for TSS usage. **(A)** Identified *ssl2* substitutions and their phenotypes relative to Ssl2 domains. Structure colored by identified Ssl2 domains as in^64^. Light grey areas in the schematic have not yet been observed in any Ssl2 or homolog structures. **(B)** Position of Ssl2 relative to TFIIH and Pol II in the yeast PIC (PDB 7O4I). **(C)** Identified substituted residues illustrated as spheres on cartoon of the Ssl2 structure from (B). Residue numbers are color coded by Ssl2 domain color from (A). **(D)** Rotation of (C) illustrating the mutants clustered at NTD/DRD/Helicase Domain (HD) 1 interface. **(E)** Spot assay showing example *rpb1* mutant transcription phenotypes. **(F)** Primer extension and quantification showing *rpb1* mutant effects on *ADH1* TSS distribution. Color coding of *rpb1* allele class in this graph is used throughout the figures. Green bars represent upstream shifting *rpb1* alleles when used to annotate figures. Blue bars represent downstream shifting *rpb1* alleles. Average of ≥ 3 biological replicates +/- standard deviation are shown. **(G)** Spot assay showing example *ssl2* mutant transcription phenotypes. **(H)** Primer extension and quantification showing example *ssl2* mutant effects on *ADH1* TSS distribution. Color coding of *ssl2* allele class in this graph is used throughout the figures. Orange bars represent downstream shifting *ssl2* alleles when used to annotate figures. Purple bars represent upstream shifting *ssl2* alleles. Average of ≥ 3 biological replicates +/- standard deviation are shown.

Some TSS shifting substitutions are located on the Ssl2 surface **(Figure 2B)** and a subset (*e*.*g*. D610) are located proximal to DNA **(Figure 2C)**. D610A confers an upstream TSS shift. In contrast, R636C, which is also close to DNA, confers a downstream TSS shift. F498, which is located in the groove of Ssl2 lobe 1 and facing DNA, caused an upstream TSS shift when substituted with leucine. Additionally, a small patch of residues with many TSS shifting substitutions is found on the Ssl2 lobe 1 surface **(Figure 2A)**. These substitutions are from the helicase domain 1 and shift TSSs upstream, including D522V, K523N, N528I, F529L and E537G. In addition, substitutions I383T and K372E are in residues proximal to this small patch but shift TSS downstream. Intriguingly a number of alleles of both classes are found in the Ssl2 N-terminal domain homologous to Tfb2 (TFB2C-like or “Clutch”) that forms interaction with Tfb2 and bridges Tfb2 with Ssl2 helicase domain 1 **(Figure 2A-D)**. As Tfb2/p52 recruits Ssl2/XPB into TFIIH^65^ and stimulates Ssl2/XPB catalytic activity^66^ this interface is of special interest for how other factors might communicate with Ssl2/XBP function in both transcription and NER. Highlighting the uniqueness and potential plasticity of this region, we identified multiple substitutions at the conserved N230 in this domain, with N230D/S conferring MPA^S^ and an upstream TSS shifts, while N230I conferred both His^+^ and Spt^-^ phenotypes and a downstream TSS shift **(Figure 2A-D)**. These results suggest that altered Ssl2 DNA or intradomain interactions alter Ssl2 function in TSS selection in distinct ways, likely through distinct effects on Ssl2 biochemical activity, discussed below and in Discussion.

### *ssl2* alleles behave differently from Pol II and other GTF alleles for TSS usage

We highlight two alleles as representative of distinct *ssl2* allele classes in comparison to examples of the two Pol II allele classes: *ssl2* N230D, which is MPA sensitive and appears to reduce ability to use downstream TSSs at *ADH1*, and *ssl2* N230I, which shows both His^+^ and Spt^-^ phenotypes and shifts TSSs downstream relative to WT at *ADH1* **(Figure 2E, F** *rpb1* alleles, **2G, H** *ssl2* alleles). As we have shown previously, *rpb1* mutants also fall into two major classes regarding transcription phenotypes and TSS shifts. As a comparison, *rpb1* E1103G is MPA^S^ and shifts TSS usage upstream while *rpb1* H1085Y is His^+^ and shifts TSS usage downstream^41, 47^. MPA-sensitivities for both *rpb1* E1103G and *ssl2* N230D correlate with an upstream TSS shift at *ADH1*, similarly to our observation for existing *ssl2* MPA^S^ alleles, *ssl2-DEAD* and *ssl2-508*. However, all *ssl2* MPA^S^ alleles examined appear to shift TSS distribution upstream by limiting or truncating TSS usage at downstream sites and not by activating lowly used upstream sites as *rpb1* E1103G does (and as known TFIIF alleles do)^16^ **(Figure 2F, H)**. This behavior suggests that *rpb1* and *ssl2* alleles may alter TSS usage by distinct mechanisms.

### Transcription start site sequencing (TSS-seq) identifies global effects on TSSs in *ssl2 alleles* in *S. cerevisiae*

To gain an insight into the impact of TFIIH’s activity on TSSs genome-wide, we have examined 5’ ends of RNA transcripts for these two classes of *ssl2* allele in *S. cerevisiae* by performing TSS-seq^13, 67^ **(Figure 3A)**. In total, six *ssl2* alleles were analyzed along with a WT control, including three His^+^ alleles (L225P, N230I and R636C) that shift TSSs downstream at *ADH1*, and three MPA^S^ alleles (N230D, D522V and Y750*) that shift TSSs upstream at *ADH1*. Furthermore, we performed TSS-seq on previously analyzed Pol II WT, E1103G and H1085Y for direct comparison purposes using our updated protocol (Methods). The positions and counts of the 5’ ends of uniquely mapped reads were extracted to evaluate correlation and assess the reproducibility between biological replicates **(Figure 3B, Figure 3 – Figure supplement 1A)**. We previously found that clustering of correlation coefficients among libraries could distinguish Pol II mutants into GOF and LOF groups^13^. Here, we also found that mutant phenotypic classes were distinguished by this clustering with *ssl2* and Pol II alleles separated suggesting effects observed at individual promoters are predictive of effects observed across the genome **(Figure 3C, Figure 3 – Figure supplement 1B)**.

**Figure 3:**
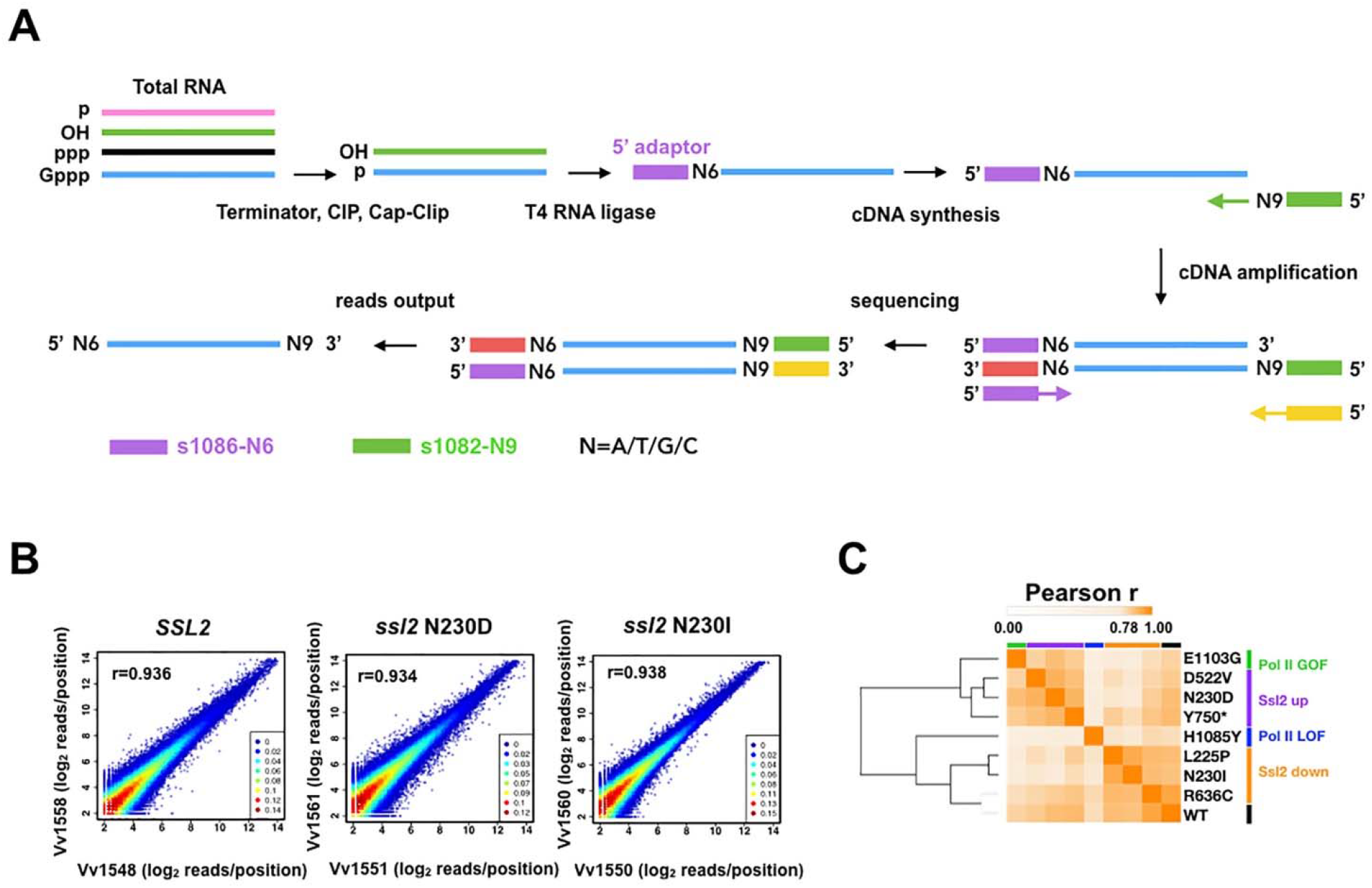
Transcription start site sequencing (TSS-seq) identifies global *ssl2* initiation effects. **(A)** TSS-seq library construction as in^67^. See Materials and Methods for details. **(B)** Scatter plot showing the correlation of log_2_ transformed reads at individual genome positions for all positions ≥3 reads for each position for example replicate pairs for *SSL2* WT, *ssl2* N230D, or *ssl2* N230I, see Figure supplements for other libraries and description of biological replicates performed for all genotypes. **(C)** Hierarchical cluster of Pearson r correlation coefficients for libraries of combined biological replicates.

### *ssl2* alleles shift transcription start site distribution genome-wide

As in Qiu, Jin *et al*^13^, we focused on initiation within a cohort of 5979 promoters for a large number of mRNA genes and non-coding RNAs. These promoters are separated into Taf1-enriched or -depleted subclasses as a proxy for the two primary promoter classes in yeast^68^. We took a simple approach to examine how mutants affect observed TSS distributions using a few key metrics, for example the position in a promoter window containing the median read in the distribution as a measure of where the TSS distribution is in a particular promoter window **(Figure 4A)**. We observed decreased TSS signal downstream of the WT median TSS signal in *ssl2* N230D and other *ssl2* MPA^S^ alleles **(Figure 4B, Figure 4 – Figure supplement 1)**. In contrast, substantially increased TSS usage was observed at downstream sites in *ssl2* N230I and other *ssl2* His^+^ alleles **(Figure 4B, Figure 4 – Figure supplement 1)**. To quantify the change of TSS usage, we measured the TSS shift between WT and mutant strains by subtracting the WT median TSS position within each promoter window from the mutant median TSS position **(Figure 4C)**. For each mutant, we show the median TSS shifts across promoters in both heatmap and boxplot format **(Figure 4D-E, Figure 4 – Figure supplement 2A)**. As predicted from our model, *ssl2* MPA^S^ alleles shift the median TSS positions upstream at most promoters, showing a similar profile to Pol II GOF mutant **(Figure 4D, Figure 4 – Figure supplement 2A)**. Also as predicted from our model, *ssl2* His^+^ alleles shift median TSS positions downstream at the majority of promoters and show a similar profile to a Pol II LOF mutant^13^ **(Figure 4D, Figure 4 – Figure supplement 2A)**. Additionally, mutants were clustered into upstream and downstream classes based on shifts in promoter median TSS position **(Figure 4D, Figure 4 – Figure supplement 2A)**. Principle Component Analysis (PCA) of TSS shifts distinguishes *ssl2* and Pol II mutants into four major classes, namely Pol II LOF (downstream shifting) and GOF (upstream shifting), and *ssl2* upstream and downstream shifting mutants **(Figure 4 – Figure supplement 2B)**. We observed that the magnitude of TSS shift was consistent with the strengths of putative TSS-shift dependent growth phenotypes in *ssl2* upstream shifting mutants **(Figure 4E)**. For example, alleles of N230D, D522V and Y750*, from the left to right, show weaker MPA^S^ phenotypes in genetic tests **(Figure 2A)**, while also showing a gradient in TSS shift magnitudes across promoters **(Figure 4E)**. Notably, the extents of TSS shifts in *ssl2* alleles are less than for Pol II activity mutants, indicating a more dramatic effect of Pol II’s catalytic activity on TSS distributions **(Figure 4E)**. These results are consistent with phenotypes of mutants being predictive of global TSS defects among mutants for a particular gene but not necessarily between genes, which we discuss later as indicative of different mechanisms for Pol II and Ssl2 effects on TSS selection.

**Figure 4:**
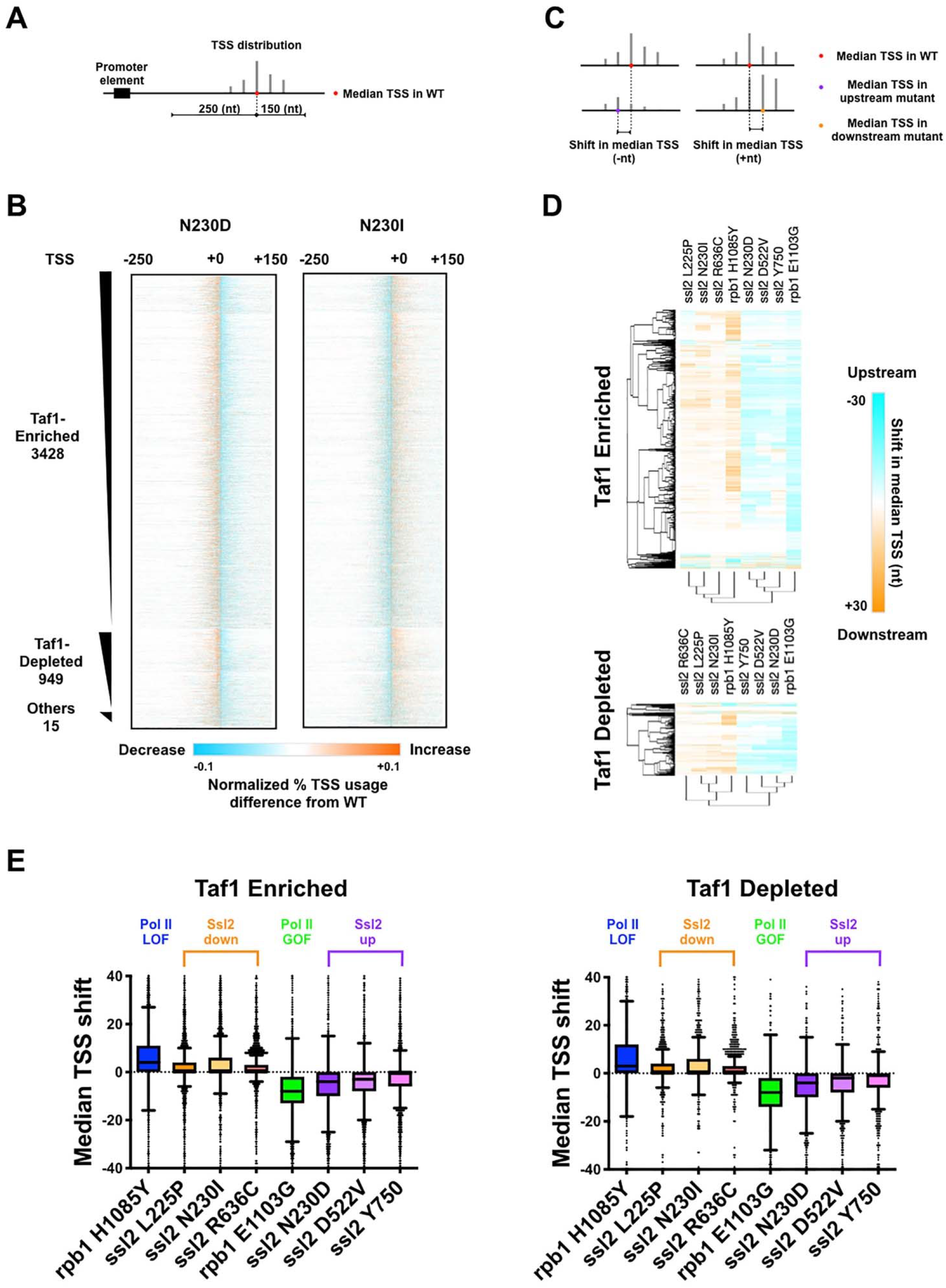
*ssl2* alleles shift transcription start site distribution genome-wide. **(A)** Schematic indicating TSS distribution within promoter window defined by median of the WT TSS distribution. **(B)** Heatmaps of TSS read differences between WT and *ssl2* mutants within defined promoter windows. Promoter classes defined by enrichment or depletion of Taf1 were rank-ordered according to total reads in WT. The promoter-normalized read differences between mutant and WT are shown by a color scheme ranging from cyan (negative)-white-orange (positive). **(C)** Schematic illustrating median TSS upstream or downstream shift in mutant relative to WT. **(D)** Heatmap and hierarchical clustering of median TSS shifts for *ssl2* or *rpb1* mutants for Taf1-enriched or -depleted promoter classes (promoters as in (B)). The shift of median TSS is indicated by cyan (upstream) or orange (downstream). **(E)** TSS shifts in *ssl2* mutants are less strong than in Pol II mutants. Median TSS shifts across promoters, regardless of promoter class are statistically distinguished from zero at p<0.0001 for all mutants (Wilcoxon Signed Rank Test).

### Distinct alterations to TSS distribution in *ssl2* mutants

To evaluate the effects of *ssl2* and Pol II alleles on scanning length, the width between positions of the 10th and 90th percentiles of the TSS signal at each promoter window was determined (TSS “spread”) **(Figure 5A)**. We observed obvious narrowing of TSS spreads in *ssl2* upstream shifting mutants and widening of TSS spreads in *ssl2* downstream shifting mutants **(Figure 5B-D)**. The profiles of the TSS spread difference between mutant and WT at each individual promoter **(Figure 5A)** also differentiates mutants into clear shift classes **(Figure 5C; Figure 5 – Figure supplement 1A, B)**. Changes in TSS spread for *ssl2* alleles are distinct from how Pol II mutants alter TSS spread **(Figure 5D)**. Both classes of *ssl2* mutant show large bias in direction of shift in spread, relative to WT across a number of types of assessment **(Figure 5 – Figure supplement 2D-F)**. Consistently, MPA^S^ *ssl2* alleles (those that shift TSSs upstream) showed narrowing in TSS spread at the majority of promoters while His+ *ssl2* alleles (those that shift TSSs downstream) showed widening in TSS spread at the majority of promoters. These results extend the idea that while classes of initiation allele may be easily identified for Pol II or Ssl2 using the same genetic phenotypes, their effects on TSSs at individual promoters are quantitatively and likely qualitatively distinct.

**Figure 5:**
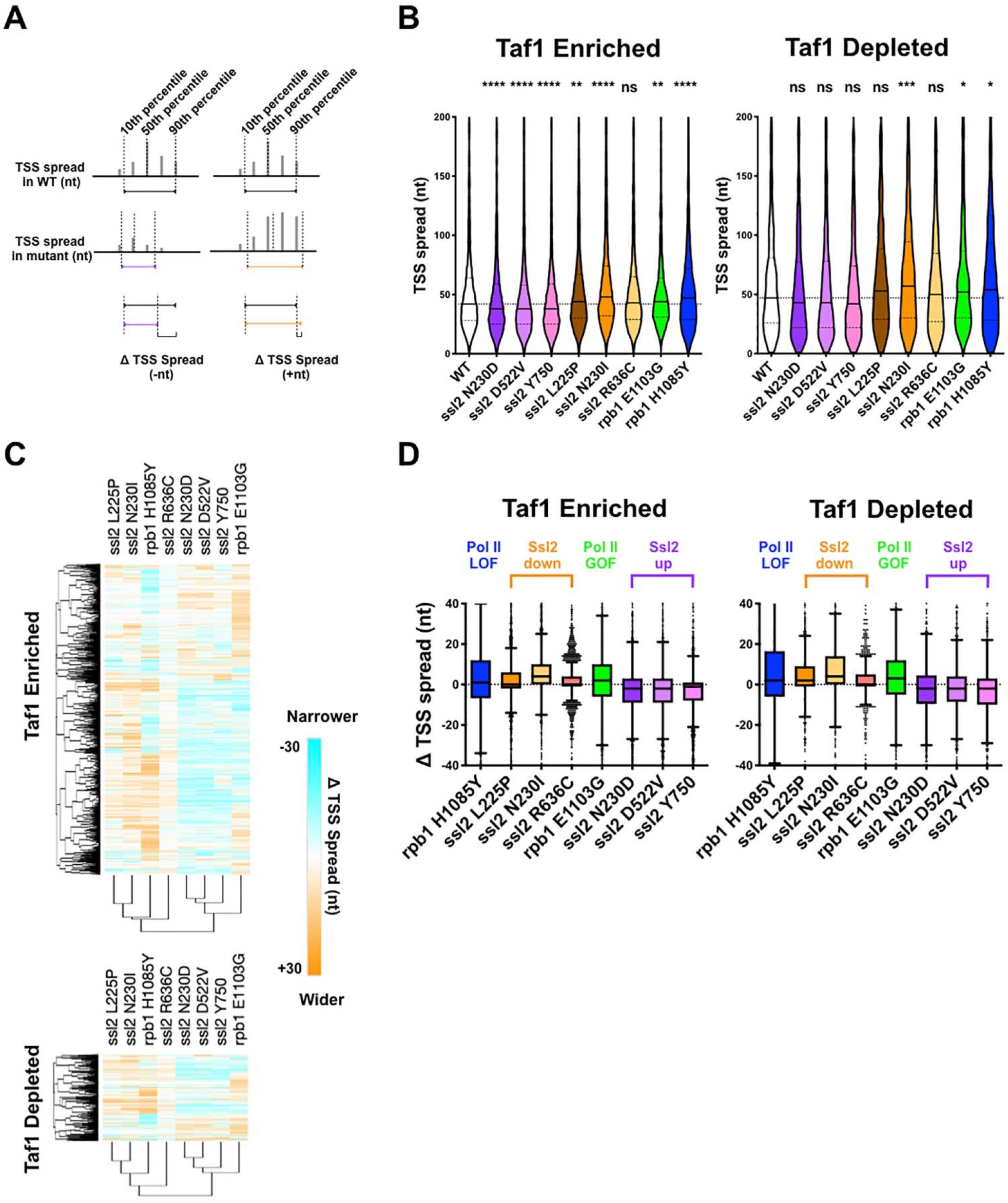
Distinct alterations to TSS distribution in *ssl2* mutants. **(A)** Schematic illustrating TSS “spread” reflecting the distance encompassing 80% of TSSs in a promoter window and the measurement of mutant changes in TSS spread. **(B)** TSS spreads in *ssl2* and Pol II mutants at Taf1-enriched or -depleted promoters. All mutants show a statistical difference in medians from WT at p<0.05 (Kruskal-Wallis test with Dunn’s multiple comparisons test). Asterisks indicate differences in means from WT (p<0.05 *, p<0.01 **, p<0.005 ***, p<0.001 ****). **(C)** Heatmap showing TSS spread changes for *ssl2* or *rpb1* mutants by promoter class (hierarchical clustering by mutant on the *x-axis* and promoter on the *y*-axis). **(D)** *ssl2* upstream and downstream shifting mutants narrow and widen TSS distributions at promoters as measured in (A). The median TSS spread changes of all the *ssl2* and Pol II mutants are statistically distinguished from zero at p<0.0001 (Wilcoxon Signed Rank Test).

### Genetic interactions between initiation factors and *ssl2* alleles suggest distinct roles for Ssl2 and other factors in TSS scanning

To explain the observed quantitative differences between Pol II initiation *efficiency* alleles and *ssl2* alleles, we hypothesize that *ssl2* alleles that narrow TSS spreads (*ssl2* N230D and similar alleles), resulting in upstream shifts in TSS distributions, are defective in scanning *processivity* due to decreased Ssl2/TFIIH translocase activity. In contrast, *ssl2* alleles that show increased TSS spreads and usage at downstream sites (*N230I* and similar alleles) behave as increased scanning processivity (GOF) alleles, due to an increase in Ssl2/TFIIH activity. Alternatively, *ssl2 N230I* might instead be a *scanning rate* GOF allele that decreases initiation efficiency across TSSs by decreasing the exposure time of each TSS within Pol II active site. As a consequence, there would be, hypothetically, increased TSS usage at downstream sites due to increased Pol II flux reaching those positions, similar to Pol II LOF efficiency alleles. To probe mechanisms of Ssl2 function we designed *ssl2* genetic interaction studies for which we have specific predictions based on their possible roles **(Figure 6 – Figure supplement 1)**. These studies emerge from our prior observations of genetic interactions from three angles between initiation factors such as Pol II activity mutants and alleles of GTFs^39^. First, we have examined general effects on growth between classes of mutant, which can manifest as synthetic sickness or lethality, suppression, or lack of effects on growth. Second, we have examined suppression or enhancement of transcription-related phenotypes in double mutants, allowing detection of additive or epistatic interactions based on putative transcription defects at the genetic reporter loci. Finally, we have quantitatively examined effects of double mutants on TSS distributions at *ADH1*, allowing additive or epistatic interactions to be observed at the individual promoter level. In our previous studies, we found that Pol II activity mutants and alleles of GTFs (*sua7*/TFIIB and *tfg2*/TFIIF) showed additivity/suppression for transcription-dependent phenotypes as well as additivity/suppression for alterations to TSS distributions at *ADH1*. These studies suggested these GTF alleles were functioning in same pathway as Pol II catalytic mutants, namely controlling the efficiency of initiation across individual TSSs. In order to understand how *ssl2* alleles interact with other initiation factors and how Ssl2 functions within the network of activities that are essential for TSS selection, we generated double mutants among a collection of *ssl2* alleles, Pol II activity-altering alleles, *sua7-1, tfg2*Δ*146-180*, and *sub1*Δ. To streamline display a large number of genetic interactions and growth phenotypes, we have converted general growth on plates and growth on phenotype-specific media to qualitative scores **(Figure 6D-F, Figure 6 – Figure supplement 2)** and these are shown as heat maps **(Figures 6, 7)** with primary data shown in **Figure 6 – Figure supplement 3, Figure 7 – Figure supplement 1 and Figure 7 – Figure supplement 2**.

**Figure 6:**
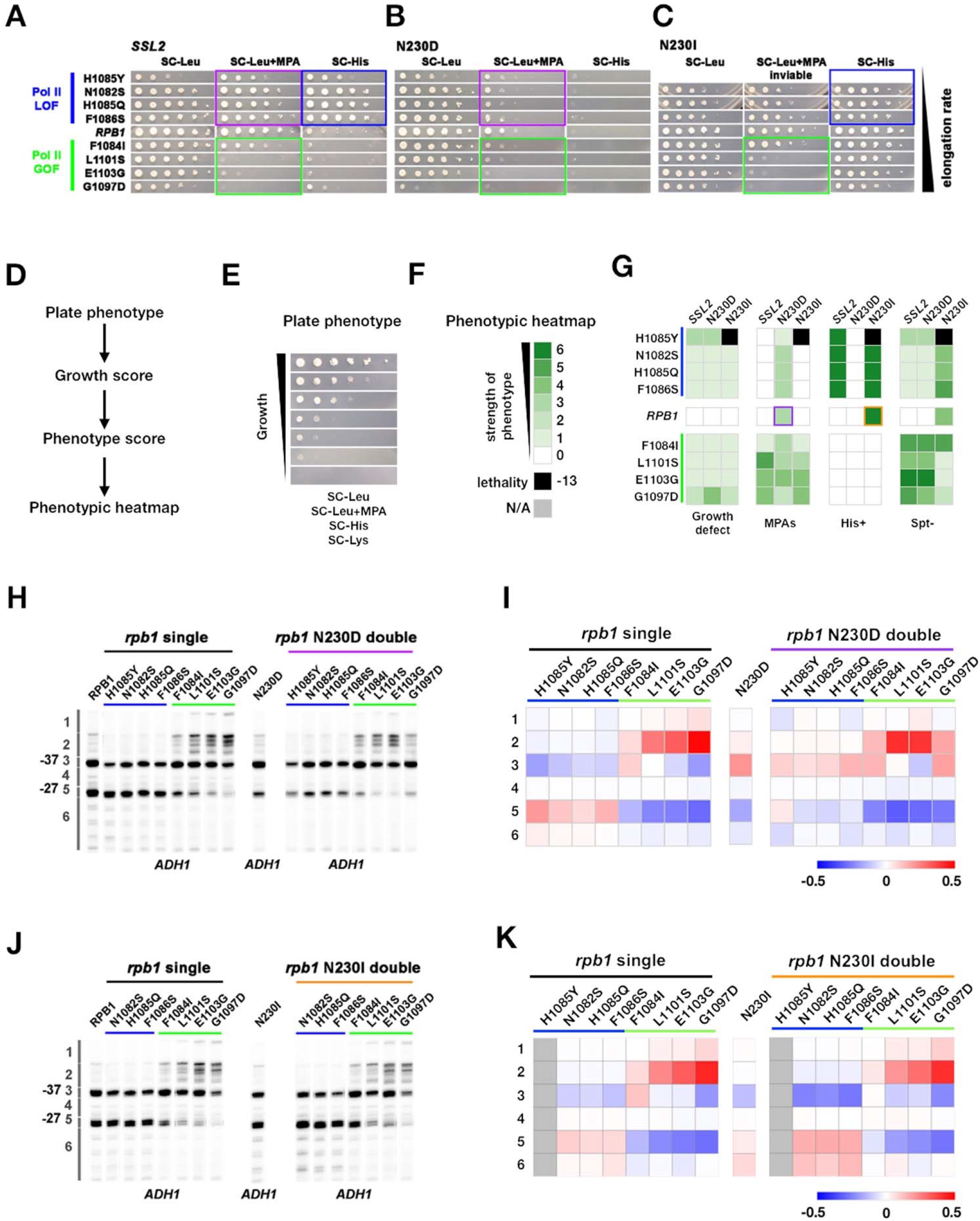
Genetic interactions between *ssl2* and Pol II initiation alleles suggest distinct functions of each in initiation by scanning. **(A)** Growth phenotypes of *rpb1, ssl2* N230D, *ssl2* N230I single or double mutants. *rpb1* mutants represent known catalytically hyperactive alleles or genetically similar (G1097D, E1103G, L1101S, F1084I) and four with reduced catalytic activity (F1086S, H1085Q, N1082S, H1085Y). Strains are arranged according to measured Pol II elongation rate *in vitro* (slowest at top). **(B)** *ssl2* MPA sensitive alleles are epistatic to Pol II LOF alleles’ His^+^ phenotypes (double mutants retain MPA^S^ of *ssl2* single mutant while His^+^ phenotypes of *rpb1* mutants are suppressed. Conversely, Pol II TSS upstream shifting alleles appear epistatic/non-additive with *ssl2* MPA^S^ alleles and do not show synthetic growth phenotypes. **(C)** Pol II upstream TSS shifting alleles appear epistatic to *ssl2* N230I phenotypes (MPA^S^ retained and His^+^ suppressed in double mutants). There are only minor synthetic defects between *ssl2* N230I and Pol II downstream TSS shifting mutants suggesting lack of synergistic defect and either mild additivity or epistasis. (Double mutant of N230I and H108Y is nearly dead and was not tested here or in E.) **(D,E)** Schematic (D) indicating how qualitative growth data of mutants encoded (E) for visualization in heat maps. **(F)** Phenotyping heat map legend. **(G)** Qualitative Heatmaps for *ssl2* and *rpb1* genetic interactions. Growth Phenotypes are detected using reporters described in Figure 1. **(H)** Primer extension of *ssl2* N230D and *rpb1* mutants at *ADH1. ssl2* N230D appears to truncate distribution of TSSs on downstream side and is epistatic to downstream shifting *rpb1* alleles (blue bar) while upstream shifting *rpb1* alleles (green bar) are non-additive or epistatic to *ssl2* N230D. Numbered regions indicate TSS positions that were binned for quantification in (I). Representative primer extension of ≥ 3 independent biological replicates is shown. **(I)** Quantification of (H) with heat map showing relative differences in TSS distribution binned by position (bins are numbered and shown in (H). Mean changes of ≥ 3 independent biological replicates are shown in the heat map. **(J)** Primer extension of *ssl2* N230I and *rpb1* mutants at *ADH1. ssl2* N230I appears to enhance usage of downstream TSSs and is additive with downstream shifting *rpb1* alleles (blue bar) while upstream shifting *rpb1* alleles (green bar) are epistatic to *ssl2* N230I. Numbered regions indicate TSS positions that are binned for quantification in (K). Representative primer extension of ≥ 3 independent biological replicates is shown. **(K)** Quantification of (J) with heat map showing relative differences in TSS distribution binned by position (bins are numbered and shown in (J). Mean changes of ≥ 3 independent biological replicates are shown in the heat map.

**Figure 7:**
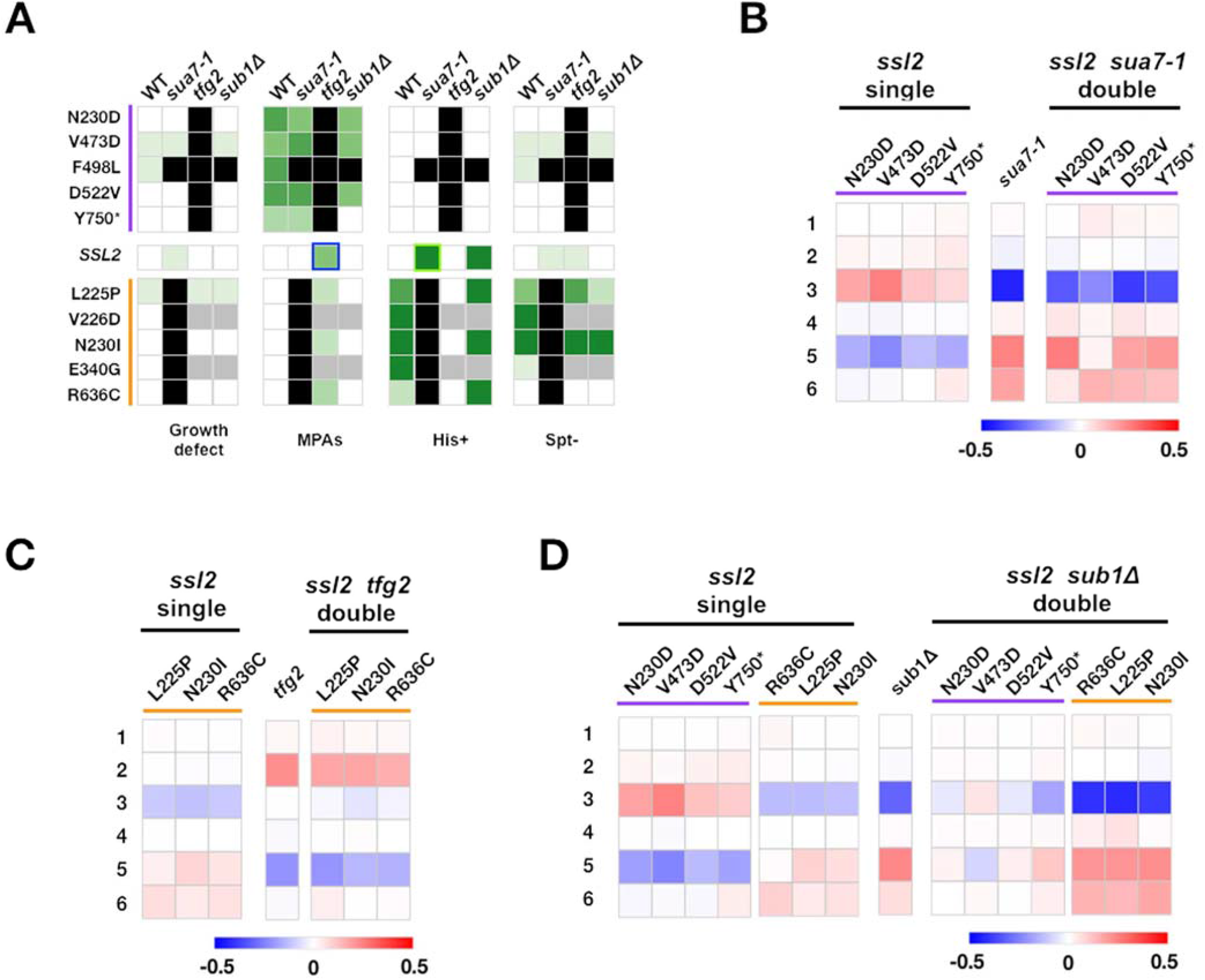
Complex genetic interactions between GTF initiation alleles suggest multiple distinct activities in initiation by scanning. **(A)** Genetic interactions between *ssl2*, TFIIB, TFIIF, and *sub1* mutants shown as a heat map indicating phenotypic strength of single and double mutants. Scaling as in 6F. **(B-D)** Heatmaps showing quantified *ADH1* primer extension data for *ssl2, sua7-1, tfg2, sub1*Δ single and double mutants. Primer extension as in Figure 1E *etc*. Mean changes of ≥ 3 independent biological replicates are shown in the heat maps.

Our detailed studies are summarized as follows (for detailed discussion, see version one of the pre-print of this work^69^). In contrast to the suppressive/additive interactions that were broadly observed between Pol II and TFIIB/TFIIF alleles^39^, we observe primarily epistatic effects between Pol II and *ssl2* alleles. First, the broad synthetic lethality or enhancement/additivity of transcriptional phenotypes through combining Pol II and TFIIB/TFIIF alleles that shift TSS distributions in the same direction were not observed between Pol II and *ssl2* alleles **(Figure 6)**. Examples of epistatic interactions, where double mutants between Pol II and *ssl2* alleles have phenotypes of either the Pol II single mutant or the *ssl2* single mutant, were found in a number of cases. Each case supports a model where *ssl2* alleles are functioning through scanning processivity and not initiation efficiency directly. This epistasis is best reflected by nearly complete absence of synthetic lethality between *ssl2* downstream shifting alleles and Pol II downstream shifting alleles **(Figure 6A-C)**, in contrast to interactions between all other classes of downstream shifting allele (*e*.*g*. Pol II, *sua7*/TFIIB, *sub1*Δ39). Epistasis for both transcription phenotypes and *ADH1* TSS shifts was also observed between Pol II upstream shifting alleles and both classes of *ssl2* allele, meaning that double mutants had phenotypes of Pol II single mutants **(Figure 6A-K, Figure 6 – Figure supplement 3)**. For each of these cases, results support a model where if initiation is efficient enough or early enough in a scanning window, *i*.*e*. due to increased Pol II initiation activity, then increase in scanning processivity (*e*.*g. ssl2* N230I) loses ability to alter TSS distributions, while a decrease in scanning processivity is buffered against due to high enough gain in transcription efficiency in tested Pol II alleles. Similarly, both classes of *ssl2* alleles appeared epistatic or non-additive with Pol II downstream shifting alleles **(Figure 6A-C, Figure 6 – Figure supplement 3)**, also consistent with determination of scanning window by *ssl2* activity to be upstream of ability of Pol II mutants to alter TSS distributions through altered initiation efficiency.

Interactions between *ssl2* alleles and other GTFs or *sub1*Δ reveal complexities that are of special note as they suggest non-obvious roles/interactions between these factors and Ssl2 function **(Figure 7, Feigure 7 – Figure supplements 1,2)**. We anticipated that *sua7-1* and *tfg2*Δ*146-180*, encoding mutant forms of TFIIB and TFIIF respectively, would behave strictly as TSS efficiency alleles due to their additive behavior with Pol II alleles^39^, and therefore would similarly show epistatic effects with *ssl2* alleles. Notably, lethal phenotypes were observed between individual *ssl2* alleles and *sua7-1* or *tfg2*Δ*146-180* alleles for combinations between single mutants that alter TSS distributions in the same direction, distinct from their interactions with Pol II alleles **(Figure 7A)**. We suggest two possibilities for this observation: first, *sua7-1* and *tfg2*Δ*146-180* could confer additional defects causing increased sensitivity to *ssl2* defects, for example, altered PIC integrity; second, *sua7-1* and *tfg2*Δ*146-180* might be sensitized to increased Ssl2 processivity (for *sua7-1*) or decreased Ssl2 processivity (for *tfg2*Δ*146-180*) in addition to their altered TSS efficiency effects (see Discussion). When we combined alleles of *sua7-1* or *tfg2*Δ*146-180* with *ssl2* alleles that shift TSS distributions in opposite directions, interactions were complex but significant epistasis was observed. Consistently, double mutants shifted *ADH1* TSS distributions to similar extent as the *tfg2*Δ*146-180* single mutant **(Figure 7C, Figure 7 – Figure supplement 1)** as predicted for an increase in initiation efficiency buffering against effects of increase in scanning processivity.

Sub1, a conserved factor (yeast homolog of mammalian PC4) was previously found to facilitate Pol II transcription in a variety of ways^70, 71^, to be recruited to the PIC^72^, and to alter accessibility of promoter single-stranded DNA, consistent with initiation functions^73^. *sub1*Δ has extensive genetic interactions with initiation factors and itself causes TSSs to shift downstream^29, 38, 50, 74^, though its actual role in initiation is unknown. We previously found *sub1*Δ to confer a His^+^ phenotype for the *imd2*Δ*::HIS3* initiation reporter^41^ and furthermore found that Pol II GOF alleles appeared epistatic to *sub1*Δ, leading to the proposal that *sub1*Δ effects in initiation were distinct from TFIIB or TFIIF alleles^39^. Because we have observed similar epistatic interactions between *ssl2* and Pol II alleles, we considered that Sub1 might also be behaving as a scanning processivity factor. Therefore, we predicted the possibility of additive effects between two types of processivity alleles, namely *ssl2* upstream and downstream shifting alleles and *sub1*Δ, if they are acting independently. First, no strong genetic interactions (lethality) were observed between *ssl2* and *sub1*Δ alleles **(Figure 7A, Figure 7 – Figure supplement 2A)**. Second, the majority of *sub1*Δ interactions with *ssl2* alleles appear to be additive when examining TSS distributions at *ADH1* as predicted for factors are acting on processivity independently. Third, and notably, we identified allele-specific interactions between *sub1*Δ and specific *ssl2* alleles within classes of *ssl2* allele that until these experiments have not been distinguishable. For example, most upstream shifting *ssl2* alleles were additive with *sub1*Δ for TSS distributions at *ADH1*, resulting in mutual suppression of TSS distribution shifts **(Figure 7D)**. In contrast, *sub1*Δ was epistatic to *ssl2* Y750*, suggesting that a putative block to processivity due to C-terminal truncation of Ssl2 can be relieved by *sub1*Δ, and potentially may be due to altered Sub1 function in *ssl2* Y750*. Finally, allele specificity of *ssl2* F498L was revealed by these genetic experiments. This TSS upstream shifting *ssl2* allele was unexpectedly synthetic lethal with both *sua7-1* and *sub1*Δ suggesting heretofore undetected phenotypic differences from other alleles of the same class.

### Two networks controlling initiation by promoter scanning

Results of genetic interaction studies are consistent with two distinct networks controlling TSS selection by scanning **(Figure 8)**. Additive/suppressive interactions were observed within networks while specific classes of mutants showed epistatic interactions between networks. One network impinges on Pol II catalysis and initiation efficiency, and genetic analyses suggest that the Pol II active site collaborates with activities of TFIIB and TFIIH in this process, consistent with experiments indicating effects of TFIIB and TFIIF on Pol II catalytic activity *e*.*g*.^21, 75, 76^. The other, we propose, impinges on scanning processivity through TFIIH with the participation of Sub1. Our genetic interactions also uncover functional connections between TFIIB and TFIIF and Ssl2 that are distinct from Pol II active site mutants. These results support predictions of altered PIC function for TFIIB and TFIIF mutants beyond phosphodiester bond formation and will be interesting to test in biophysical experiments. Extensive epistasis observed between networks **(Figure 8)** supports predictions for how efficiency and processivity should interact during initiation by promoter scanning (**Figure 9**, see Discussion).

**Figure 8:**
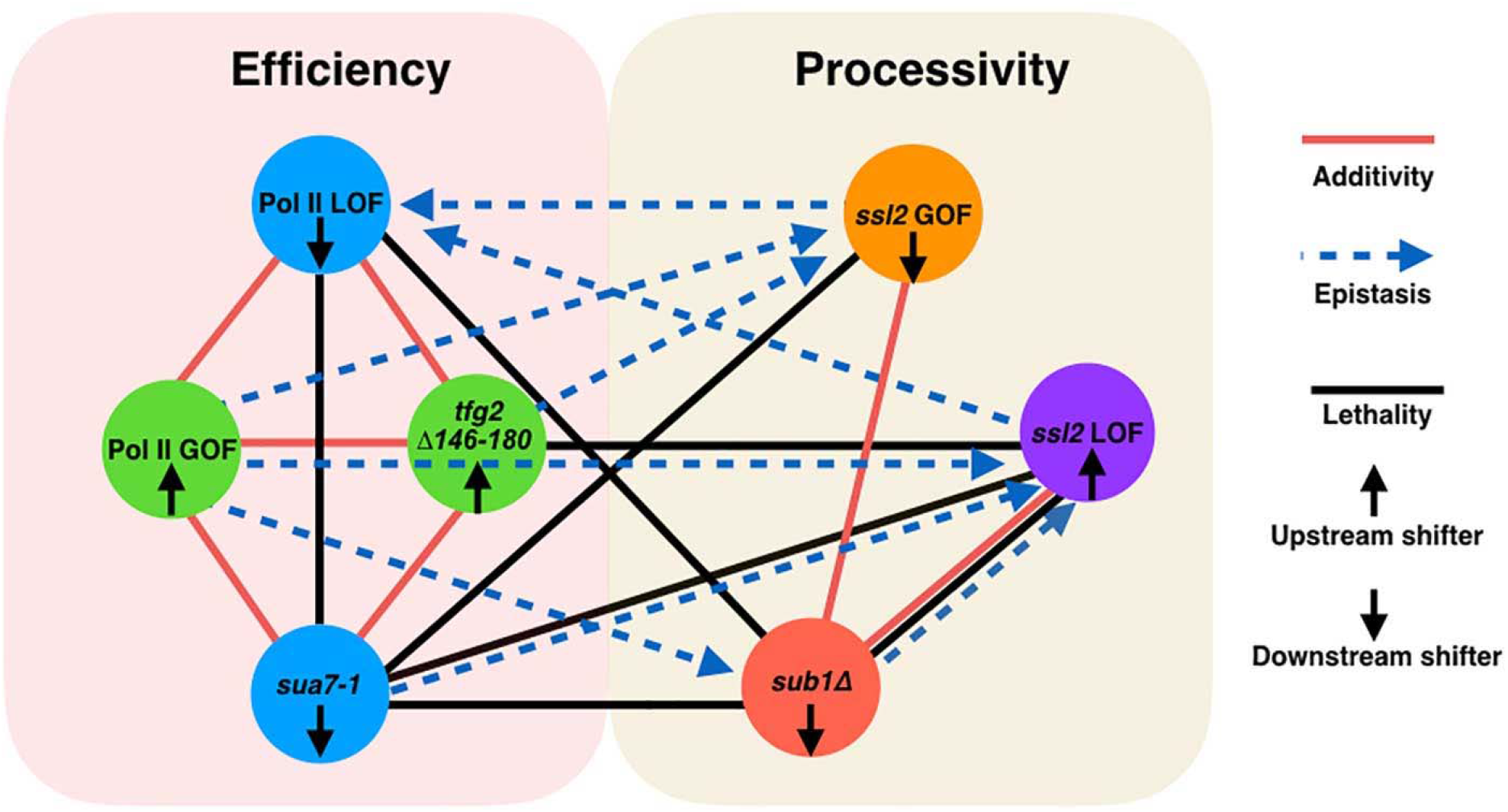
Two major functional networks controlling initiation by scanning. In our genetic experiments, additive/suppressive effects are mainly observed between alleles predicted to function by alteration to initiation efficiency (*rpb1* and tested alleles of TFIIB/TFIIF). Multiple lines between classes indicates allele specific interactions between a factor and individuals of an allele class *e*.*g. sub1*Δ and *ssl2* LOF alleles. In contrast to interactions within the “efficiency” network, widespread epistasis was observed between *ssl2* and other factors as predicted for interactions between processivity and efficiency alleles. *sub1*Δ generally shows additivity/suppression with *ssl2* alleles, consistent with it functioning as a scanning processivity factor. Unique lethal interactions between *ssl2* LOF upstream shifter F498L and *sua7-1* and *sub1*Δ indicate distinct behavior within the *ssl2* LOF class.

**Figure 9:**
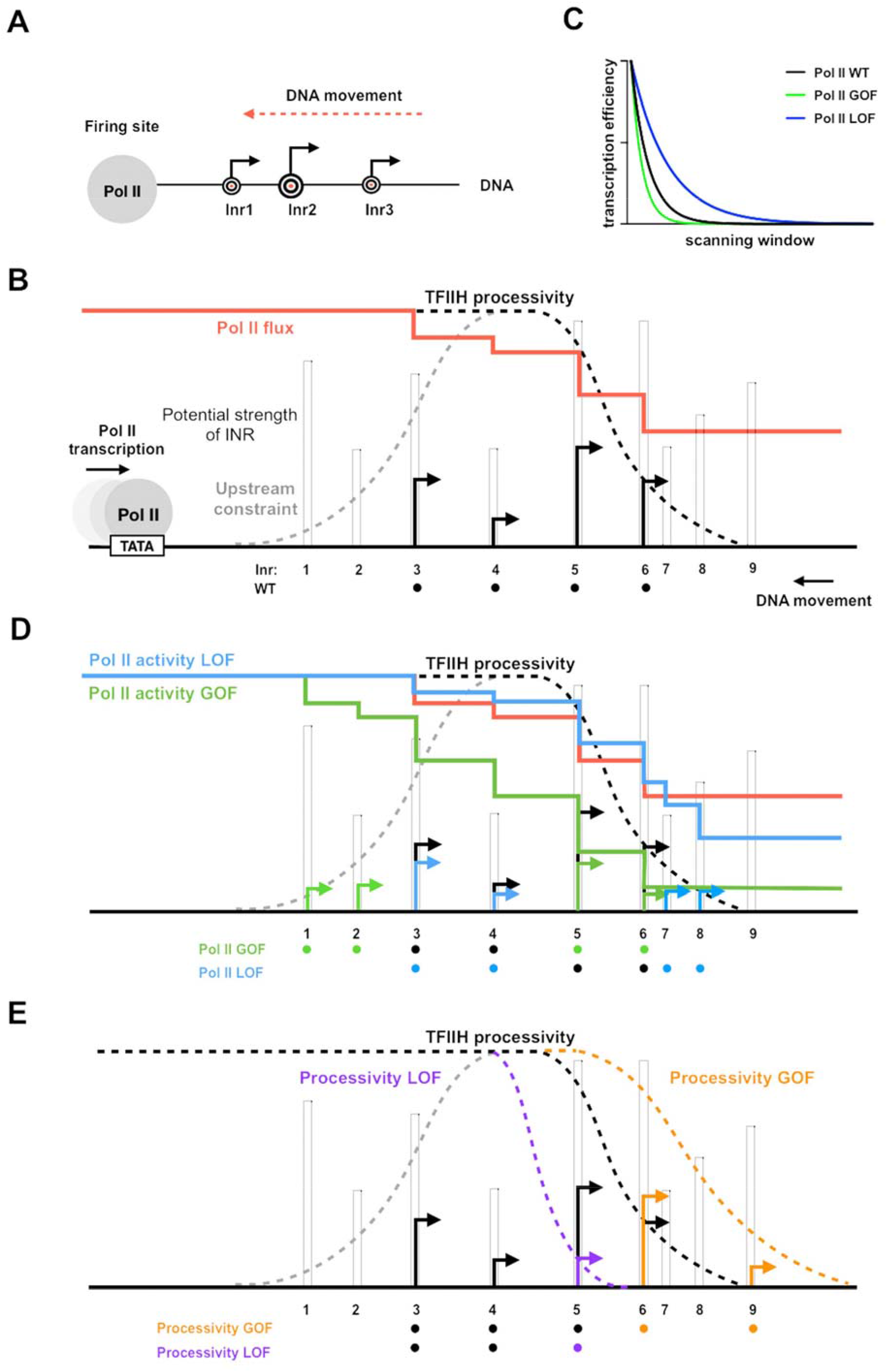
Model for interaction between initiation efficiency and scanning processivity. **(A)** The “Shooting Gallery” model. The Pol II active site controls initiation efficiency, *i*.*e*. “the rate of firing”. TFIIH controls the rate of the extent of scanning, *i*.*e*. “the speed of target passage and number of targets reached.” **(B)** Reduction in relative TSS usage as scanning Pol II initiates. As Pol II (WT) scans from upstream to downstream, successful initiation at upstream positions will reduce the amount of Pol II continuing to scan downstream. Increasing initiation efficiency at each position as is predicted for increased Pol II catalytic activity will result in a more rapid decrease in observed initiation from upstream to downstream. Conversely, reducing initiation efficiency at each position will flatten observed TSS distribution because more Pol II will reach downstream positions. **(C)** TSS distributions during promoter scanning in the “Shooting Gallery” model. The TSS distribution (black arrows) of a promoter window can be affected by Pol II catalytic activity, PIC scanning rate and processivity, TSS strength, Pol II flux, and additional observed (upstream limitation on initiation too close to PIC assembly) or potential (downstream limitation through chromatin structure) constraints. **(D)** Effects of Pol II catalytic activity on TSS distributions. Increased Pol II catalytic activity increases the efficiency of upstream TSSs that are encountered by Pol II and decreases the usage of downstream TSSs due to quickly reduced Pol II flux (changes indicated as green arrows). Decreased Pol II catalytic activity decreases TSS efficiency of upstream TSSs encountered by Pol II and increases apparent TSS usage at downstream sites due to failed upstream initiation, resulting in a downstream shifted TSS distribution within a window determined by PIC scanning potential (changes shown as blue arrows). **(E)** Effects of altered scanning processivity on TSS distributions. Increased processivity alleles are hypothesized to increase the probability of Pol II scanning further downstream if Pol II flux remains, thus expanding the scanning window and allowing Pol II usage of downstream TSSs if Pol II flux is not limiting (orange TSS). In contrast decreased processivity will limit Pol II scanning downstream, truncating the distribution of observed TSSs (purple TSS).

### *ssl2* alleles shift positioning of PIC-components genome-wide

We found previously that polar shifts of TSS distribution in Pol II catalytic activity mutants was accompanied with alteration in PIC localization as detected by ChIP-exo^13^. Shift in PIC components upon alteration to Pol II catalytic activity suggested that extent of scanning might be coupled to Pol II initiation, or that alteration to Pol II initiation kinetics affects observed distributions of GTFs. We performed these same ChIP-exo experiments on Sua7 and Ssl2 for two *ssl2* alleles, N230D and N230I. **(Figure 9 – Figure supplement 1A, B)**. Both shifted PIC localization genome-wide with the same polarity as they shift TSS distribution. We note that TAP-tagging Ssl2 confers slight phenotypes on its own and slight enhancement of *ssl2* N230D and slight suppression of *ssl2* N230I **(Figure 9 – Figure supplement 2)**. However, each tagged mutant was compared to the tagged WT and the results are robust and distinct for each mutant. The extent of ChIP-exo shifts were as strong or stronger than Pol II mutant shifts although Pol II mutants have stronger effects on TSS distributions^13^. These results could be consistent with scanning by TFIIH on DNA uncoupled from the Pol II initiation decision, i.e. *ssl2* mutants extend PIC scanning to downstream positions even though initiation has occurred. Such a result would be consistent with similar behavior of DNA compaction in optical tweezer analysis of initiation wherein dATP-supported reactions (presumptive TFIIH scanning-driven DNA translocation through Ssl2 use of dATP) and NTP-supported initiation reactions (TFIIH translocation and Pol II initiation allowed) have similar behavior^17^. The extent or mechanism of uncoupling between promoter scanning upon productive initiation are unknown and represent an open question in initiation mechanisms.

## DISCUSSION

Our studies now reveal the impact of altered Ssl2 function on initiation by promoter scanning in *S. cerevisiae*. We find distinct classes of Ssl2 allele that alter initiation genome-wide with distinct behaviors, with many alleles being in highly conserved residues. Our genomic and genetic data support a model wherein Ssl2 function as a DNA translocase can be genetically modulated and this modulation is consistent with TFIIH having either increased or decreased processivity during promoter scanning. The positioning and genetic behaviors of our allele classes are consistent with one class behaving biochemically as a loss of function and therefore truncating the scanning process prematurely and narrowing TSS distributions genome-wide. This is exactly the predicted outcome for a translocase with decreased processivity. Conversely, our other class increases downstream TSS usage and alters PIC localization at promoters by extending it downstream. These behaviors are consistent with increased translocase processivity. Genetic interactions between *ssl2* alleles, especially between *ssl2* and GTF mutants and *sub1*Δ suggest further distinctions between allele classes or within allele classes, generating testable predictions for biochemical studies. Putative increased *SSL2* activity alleles are also dominant or codominant genetically, consistent with being able to function on promoters in an increased capacity of some sort.

### Interpretation of Pol II and Ssl2 functions in the Shooting Gallery model for initiation by scanning

We have previously described how Pol II determines the *efficiency* of a TSS in a “Shooting Gallery” model, where the rate at which a TSS (conceived of as a target) passes the active site, the rate of firing (catalytic activity), and the size of the target (innate sequence strength) together contribute to the probability a target is hit (initiation happens) **(Figure 9A)**^13, 40^. Alteration of enzymatic activities supporting initiation, either the Pol II active site or TFIIH translocation will have predictable effects on individual TSS usage and the overall TSS distributions when initiation proceeds by scanning. In Pol II mutants with altered catalytic activity that is known to affect transcription efficiency, we observed polar changes to TSS distributions^13^. Distributions will also necessarily be shaped by an additional factor: Pol II flux. Pol II flux describes the relative number of polymerases encountering a given start site, which has a higher value at upstream TSSs and a lower value at downstream TSSs, resulting in reduced apparent usage at downstream position distinct from their inherent efficiencies **(Figure 9B)**. Additionally, the potential upstream and downstream constraints for defining the scanning window will have effects on TSS distributions. Studies suggest that very upstream TSSs close to the presumed location of PIC assembly show reduced transcription initiation^27^. The physical basis for defining the upstream boundary of the scanning window has not yet been determined. An obvious constraint is the minimum space required for PIC assembly. Moreover, we hypothesize that downstream constraints for defining the scanning window could be TFIIH’s processivity, the +1 nucleosome, or both. Previous single molecule studies suggested that TFIIH drives downstream scanning distances similar in length to the distribution of TSSs at yeast promoters^17^. We propose that TSS distribution of a promoter is established by the cooperation of Pol II’s catalytic activity and TFIIH’s processivity for reaching and activating TSSs at promoter sites.

When Pol II has increased catalytic activity, *e*.*g*. in Pol II catalytic activity GOF alleles, upstream TSSs will increase in efficiency **(Figure 9C, D, Pol II GOF)**. In this allele class, usage of downstream TSSs also will decrease due to reduction in Pol II flux reaching downstream sites due to prior initiation. Conversely, when Pol II has decreased catalytic activity, TSSs at upstream sites will be less efficiently used, more slowly reducing Pol II flux **(Figure 9C, D, Pol II LOF)**. Inability to initiate earlier in scanning will result in increased TSS usage at downstream sites and a flattening and spreading of the TSS distribution (as demonstrated by an efficiency curve with decreased slope). We hypothesize that alleles with increased processivity (*processivity* GOF allele) will expand the scanning window by allowing the PIC to scan further downstream while attempting initiation, increasing the probability that downstream TSSs are reached during any individual scanning event **(Figure 9E, processivity GOF)**. As a consequence, *processivity* GOF alleles increase the potential for scanning downstream but only if Pol II flux (Pol II molecules still scanning) persists to reach those sites. In contrast, a *processivity* LOF allele would limit the Pol II machinery’s access to downstream TSSs sites by reducing the scanning window **(Figure 9E, processivity LOF)**. Consequently, there would be an upstream shift in TSS distribution compared to WT, without the activation of additional upstream TSSs.

Prior biochemical and more recent structural analyses indicate that TFIIH is a fascinating complex with numerous contacts suggested or predicted to modulate or control TFIIH enzymatic subunits’ activities^77^. For example, Ssl2/XPB must be activated during transcription initiation to allow promoter opening. Recent structures suggest that interactions with both Mediator and TFIID may position parts of TFIIH for different functions in initiation^78-80^. Genetic and biochemical studies also suggest that TFIIH may itself impose a block to initiation that is then relieved by TFIIH activity through Ssl2/XPB^81, 82^. Both TFIIH ATPase subunits, Rad3/XPD and Ssl2/XPB, must also be regulated for TFIIH’s function in NER with Rad3/XPD held inactive during transcription and released for NER^64, 77^. XPB mutations in patients that result in Xeroderma Pigmentosum are straightforwardly interpreted as conferring NER defects, however transcriptional phenotypes may also be present depending on mutation^51-53^. Mutations in XPB that cause Trichothiodystrophy (TTD) localize to conserved residues in the XPB N-terminus where we have identified a number of mutations. TTD mutations in XPB have been interpreted as reducing the amount of TFIIH in the cell through potential destabilization, while one appears to impact folding and activity of XPB^64^. Our identification of putative loss and gain of function mutations in this domain in *S. cerevisiae* underscore the idea that observed conservation in this region may control key inputs to Ssl2/XPB activity. The substitutions we have identified are largely in residues conserved from yeast to humans (**Figure 2 – Figure supplement 3**), and we suggest these residues detect potential paths for allosteric regulation of Ssl2/XPB. Only a subset of our alleles confer UV sensitivity, suggestive of NER defects^83^. These are C-terminal and this suggests that our alleles uniquely alter Ssl2 modulation in transcription or that transcriptional functions of Ssl2 are sensitized to defects that do not appreciably lead to UV sensitivity. DNA translocases are the engines for chromatin remodeling and much regulation of chromatin remodelers relates to coupling of ATP hydrolysis to translocation potential^84, 85^. Mutations in remodelers that increase or decrease coupling have strong effects on remodeling. We posit that it is likely that a number of our alleles will act through altered coupling of ATPase activity and translocation, with the end result being increase or decrease in scanning processivity. Biochemical studies will reveal specific aspects of TFIIH activity that are altered by these substitutions.

Our putative gain-of-function alleles are concentrated in the TFB2C-like N-terminal domain and DRD of Ssl2. The TFB2C-domain has been implicated as a target of Tfb3/Mat1 in restricting Ssl2/XPB activity in holoTFIIH^62, 77, 86^. However, upon assembly into the PIC, Tfb3/Mat1 releases the N-terminus of Ssl2^78, 87^. This Ssl2 domain is also targeted by Tfb2/p52^61, 63, 64, 87^, which has long been implicated in modulating Ssl2/XPB activity in addition to assembling it into TFIIH. We have identified one His^+^ allele at the Ssl2-Tfb2 interface, though most are in internal interfaces at the putative nexus of Ssl2 NTD, DRD, and HD1/ATPase Lobe 1 (**Figure 2**).

Key open questions relate to how the PIC communicates to Ssl2/XPB to engage and open promoter DNA, what the mechanistic basis for any imposition of initiation block by Ssl2/XBP is, what the basis of its subsequent relief is, and how translocation is terminated upon or subsequent to productive initiation, *i*.*e*. are the processes coupled in anyway. Aibara, Schilbach *et al* have recently imaged the human PIC captured in two states, suggestive of pre- and post-translocation intermediates of TFIIH^88^. These structures show loss of contact between TFIIH through the MAT1 RING domain and the Pol II stalk/TFIIE in the proposed post-translocated state. This raises an attractive model for uncoupling TFIIH translocase from Pol II after a single translocation step, functionally limiting initiation to the small window of exposed TSSs within reach of the Pol II active site. These structures were from a minimal PIC lacking Mediator and TFIID and therefore it remains to be determined if this translocase is in fact uncoupled, as other potential TFIIH/PIC contacts remain. Other events during initiation may also propagate changes to the PIC, such as lengthening of the nascent RNA to potentially clash with TFIIB and potential subsequent reorganization of the PIC by this event, or due to Pol II CTD phosphorylation. In many organisms, nucleosome-depleted regions promote initiation bidirectionally and these regions are flanked by positioned nucleosomes^89-92^. Nucleosomes would potentially act as competitors for double stranded DNA being translocated by TFIIH. Transcription activity drives histone dynamics at promoters, consistent with TFIIH translocation proposed to function akin to a chromatin remodeler^93^. How the +1 nucleosome might feed back on scanning or regulate TFIIH translocation is an open question. In the absence of the remodeler RSC’s function in yeast, nucleosomes move upstream into normally nucleosome depleted regions and inhibit or narrow TSS usage at a number of promoters^94^. These results are consistent with nucleosomes competing with TFIIH for promoter DNA. However, these results are conditional on RSC depletion, and it is not clear if promoter nucleosome remodeling under normal conditions obviates the ability for +1 nucleosomes at active promoters to provide a block to scanning or initiation on the edges of their positions.

How Pol II specifies multiple TSSs at individual promoters across eukaryotes has long been an open question. The observation of downstream-located TSS sites relative to where DNA melting occurs in *S. cerevisiae* led to the original proposal of a scanning mechanism for TSS identification^9^. In contrast to how Pol II finds TSSs in *S. cerevisiae* for all promoters, in other eukaryotes including other fungi, a scanning process is not required for promoters with defined architecture specified by a TATA element^95, 96^. These promoters use highly focused TSSs immediately and precisely downstream of the DNA melting sites ∼30 basepairs downstream of the TATA element +1 position. For example, Lu *et al* have pinpointed the split between scanning from TATA-promoters and non-scanning species within the Saccharomycetes^95^. However, most eukaryotic promoters are TATA-less and use multiple TSSs^89, 97-107^. Therefore, whether or not scanning is also a mechanism in higher eukaryotic promoters, or minimally for a subset of eukaryotic promoters or within a specified window, is still an unanswered question as there have been no formal tests of this mechanism. It has been suggested, however, that each individual TSS is recognized as an individual promoter due to sequence signatures apparent in comparison of thousands of TSSs in humans^108^. Very recent results of cryo-EM studies on human PICs, especially in structures visualizing TFIID, indicate that promoter classes may assemble PIC components in distinct fashion within a single organism^79^, yet these assembly pathways similarly position an upstream TSS proximal to the Pol II active site, consistent with proposals that human promoters could contain information for assembling PICs individually^108^. Recent results suggest that there may be plasticity in TSS selection from individual PICs upon mutation of Inr sites in mouse, potentially supporting a type of scanning in mammals^109^. That diverse initiation mechanisms are supported by highly conserved factors suggests that we may yet to find additional unexpected plasticity in initiation across evolution in eukaryotes.

## METHODS

### Yeast strains

Yeast strains are derived from a *GAL*^*+*^ derivative of S288C (FY2)^110^. Yeast strains used in this study are listed in **Additional Table 1**.

### Plasmids and bacteria strains

Bacterial strains and plasmids used in this study are listed in **Additional Table 2**.

### Yeast media

Yeast media used in this study were made as previously described^39, 41, 47, 111^. Briefly, YP medium is made of yeast extract (1% w/v; BD) and peptone (2% w/v; BD). Solid YP medium contained bacto agar (2% w/v; BD), adenine (0.15 mM, Sigma-Aldrich) and tryptophan (0.4 mM, Sigma-Aldrich). YPD medium uses YP medium components supplemented with dextrose (2% w/v, VWR). YPRaf medium uses YP medium components supplemented with raffinose (2% w/v, Amresco) and antimycin A (1 mg/ml; Sigma-Aldrich). YPRafGal medium uses YP medium components and supplemented with raffinose (2% w/v), galactose (1% w/v; Amresco) and antimycin A (1 mg/ml; Sigma-Aldrich). Minimal media (SC-) was made with a slightly modified “Hopkins mix” (0.2% most amino acids w/v), and supplemented with Yeast Nitrogen Base containing ammonium sulfate (without amino acids, BD), bacto agar (2% w/v; BD) and dextrose (2% w/v, VWR). The original “Hopkins mix” and the slight modification were as previously described^47, 111^. All amino acids are from Sigma-Aldrich. SC-Leu+5FOA is minimal medium of SC-Leu supplemented with 5-fluoroorotic acid (5-FOA), (1 mg/ml, Gold Biotechnology). SC-Leu+MPA media is minimal media of SC-Leu supplemented with mycophenolic acid (20 ug/ml, Sigma-Aldrich, from a 10 mg/ml stock in ethanol). SC-His+3AT is minimal medium of SC-His supplemented with 3-aminotriazole (3-AT, 0.5mM, Sigma-Aldrich).

### Plate phenotyping and growth heatmaps

Yeast phenotyping assays were performed by spotting 10-fold serial dilutions of saturated YPD-liquid yeast cultures on various solid media, as previously described^47^. Yeast cells on various media were cultured at 30 °C except for temperature sensitivity phenotypes, which were at 16 °C (YPD 16) and at 37 °C (YPD 37). Yeast growth on specific media was recorded by taking pictures every 24 hours after an initial 16h of growth, from day 2 (40h) to day 7 for all media except for YPRaf/Gal (pictures to day 9). Growth phenotypes on specific media were scored on days when WT yeast reached mature colony sizes, as follows: YPD on day 2 (40h after spotting); SC-Leu, SC-His and SC-Trp on day 3 (64h); YPRaf on day 4 (88h); SC-Lys and SC-Leu+MPA on day 5 (112h); and YPRaf/Gal on day 7 (160h). To illustrate the strength and distribution of mutants on the two-dimensional structure of Ssl2 (**Figure 2**), growth phenotypes are converted to a numerical score, using the scale 1-5 to indicate the level of growth, where 1 indicates no growth and 5 indicates full growth. The level of growth is positively correlated with the strength of phenotypes for SC-His, SC-Lys and YPRaf/Gal medium, so the “growth score” is directly used as “phenotyping score” for making a heat map. For other media, the level of growth is negatively correlated with the strength of growth, thus growth score 1-5 is inversely converted to the phenotyping strength score 5-1, with 5 growth score converted into phenotyping score 1 to indicate no phenotypes, with 4 growth score converted into phenotyping score 2 to show a weak phenotype, and so on. The heatmap uses light to dark color showing weak (phenotyping score 2) to strong phenotypes (phenotyping score 5), no phenotype (phenotyping score 1) is not shown.

### Primer extension

To detect putative usage of TSSs in yeast, a primer extension (PE) assay was performed as previously described^47^, modifying original protocol in ^112^. Briefly, 30µg of total RNA isolated from yeast cells was used for each PE reaction. A primer complementary to *ADH1* mRNA was end-labeled with gamma-P32 ATP and T4 PNK and annealed to total RNA. Reverse-transcription was then performed by adding M-MLV reverse-transcriptase (Fermentas/ThermoFisher) and RNase Inhibitor (Fermentas/ThermoFisher). RNase A was added to remove RNA after reverse-transcription. Products were detected by running an 8% acrylamide gel (19:1 acrylamide:bisacrylamide) (Bio-Rad), 1X TBE, and 7M urea. PE gels were visualized by phosphorimaging (GE Healthcare or Bio-Rad) and quantified by ImageQuant 5.1 (GE) or Image Lab software (Bio-Rad).

### *ssl2* mutant screening

*ssl2* mutants were created by PCR-based random mutagenesis coupled with a gap repair. Briefly, mutation of *SSL2* (*ssl2\**) was accomplished by standard PCR reactions using Taq polymerase (New England Biolabs). *ssl2\** PCR products were then transformed into yeast along with a linearized pRS315 *SSL2 LEU2* plasmid with most of the wild-type *SSL2* sequence removed by restriction digest. Leu^+^ transformants were selected. Homologous sequences on each end of the *ssl2\** PCR products and the gapped *SSL2* vector allowed homologous recombination, resulting in a library of gap-repaired plasmids containing potential *ssl2\** alleles. Since *SSL2* is essential, these yeast cells are pre-transformed with a pRSII316 *SSL2 URA3* plasmid to support growth, while the genomic *SSL2* was deleted to allow plasmid *SSL2* alleles to exhibit phenotypes. After gap repair, cells retaining pRSII316 *SSL2 URA3* plasmids were killed by replica-plating transformants to medium containing 5FOA. Yeast cells were then plated on YPD media for growth and replica-plated to a variety of media to screen for mutants that have transcription-related or conditional phenotypes. Plasmids from yeast mutants were recovered and transformed into *E. coli* for amplification, followed by sequencing to identify mutations. Mutant yeast candidates were additionally mated with yeast cells that contain a WT *SSL2 URA3* plasmid to create diploid strains and perform phenotyping again to determine dominance/recessivity of *ssl2* mutations. Plasmid shuffling on diploid strains was performed by adding 5FOA on the medium so that presumably only presumptive *ssl2\** was kept. This was followed by an additional phenotyping to determine if the mutant phenotype is plasmid linked or not. All mutants described here were verified by retransformation into a clean genetic background.

### Transcription start site sequencing (TSS-seq)

Yeast cell cultures were grown in triplicates and cells were harvested at mid-log phase at a density of 1×10^7^ cells/mL, as determined by cell counting. For *S. cerevisiae* TSS-seq, cells collected from 50 mL of *S. cerevisiae* culture and 5 mL of *S. pombe* culture were mixed and total RNA was extracted as described^113^. We performed cDNA library construction for TSS-seq essentially as described by Vvedenskaya *et al*^67^, steps are described as follows. 100 μg of the isolated total RNA was treated with 30 U of DNase I (QIAGEN) and purified using RNeasy Mini Kit (QIAGEN). A Ribo-Zero Gold rRNA Removal Kit (Illumina) was used to deplete rRNAs from 5 μg of DNase-treated RNAs. The rRNA-depleted RNA was purified by ethanol precipitation and resuspended in 10 μL of nuclease-free water. To remove RNA transcripts carrying a 5’ monophosphate moiety (5’-P), 2 μg of rRNA-depleted RNA were treated with 1 U Terminator 5’-Phosphate-Dependent Exonuclease (Epicentre) in the 1x Buffer A in the presence of 40 U RNaseOUT in a 50 μL reaction at 30 °C for 1 h. Samples were extracted with acid phenol-chloroform pH 4.5 (ThermoFisher Scientific), and RNA was recovered by ethanol precipitation and resuspended in 30 μL of nuclease-free water. Next, to remove 5’-terminal phosphates, RNA was treated with 1.5 U CIP (NEB) in 1x NEBuffer 3 in the presence of 40 U RNaseOUT in a 50 μL reaction at 37 °C for 30 min. Samples were extracted with acid phenol-chloroform and RNA was recovered by ethanol precipitation and resuspended in 30 μL of nuclease-free water. To convert 5’-capped RNA transcripts to 5’-monophosphate RNAs ligatable to 5’ adaptor, CIP-treated RNAs were mixed with 12.5 U CapClip (Cellscript) and 40 U RNaseOUT in 1X CapClip reaction buffer in a 40 μLreaction and incubated at 37 °C for 1 h. RNAs were extracted with acid phenol-chloroform, recovered by ethanol precipitation and resuspended in 10 μL of nuclease-free water. To ligate the 5’ adapter, the CapClip-treated RNA products were combined with 1 μM 5’ adapter oligonucleotide s1086 (5’-GUUCAGAGUUCUACAGUCCGACGAUCNNNNNN-3’), 1X T4 RNA ligase buffer, 40 U RNaseOUT, 1 mM ATP, 10% PEG 8000 and 10 U T4 RNA ligase 1 in a 30 μL reaction. The mixtures were incubated at 16°C for 16 h and the reactions were stopped by adding 30 μL of 2X RNA loading dye. The mixtures were separated by electrophoresis on 10% 7 M urea slab gels in 1X TBE buffer and incubated with SYBR Gold nucleic acid gel stain. RNA products migrating above the 5’ adapter oligo were recovered from the gel as described^34^, purified by ethanol precipitation and resuspended in 10 μL of nuclease-free water. To generate first strand cDNA, 5’-adaptor-ligated products were mixed with 0.3 μL of 100 μM s1082 oligonucleotide (5’-GCCTTGGCACCCGAGAATTCCANNNNNNNNN3’, N=A/T/G/C) containing a randomized 9-nt sequence at the 3’ end, incubated at 65 °C for 5 min, and cooled to 4 °C. A solution containing 4 μL of 5X First-Strand buffer, 1 μL (40 U) RNaseOUT, 1 μL of 10 mM dNTP mix, 1 μL of 100 mM DTT, 1 μL (200 U) of SuperScript III Reverse Transcriptase and 1.7 μL of nuclease-free water was added to the mixture. Reactions were incubated at 25 °C for 5 min, 55 °C for 60 min, 70 °C for 15 min, and cooled to 25 °C. 10 U RNase H was added, the mixtures were incubated 20 min at 37 °C and 20 μL of 2X DNA loading solution (PippinPrep Reagent Kit, Sage Science) were added. Nucleic acids were separated by electrophoresis on 2% agarose gel (PippinPrep Reagent Kit, external Marker B) to collect species of ∼90 to ∼550 nt. cDNA was recovered by ethanol precipitation and resuspended in 20 μL of nuclease-free water. To amplify cDNA, 9 μL of gel-isolated cDNA was added to the mixture containing 1X Phusion HF reaction buffer, 0.2 mM dNTPs, 0.25 μM Illumina RP1 primer (5’-AATGATACGGCGACCACCGAGATCTACACGTTCAGAGTTCTACAGTCCGA-3’), 0.25 μM Illumina index primers RPI3-RPI16 (index primers have the same sequences on 5’ and 3’ ends, but different on 6-nt sequence that serves as a barcode (underlined); RPI3: 5’-CAAGCAGAAGACGGCATACGAGAT<u>GCCTAA</u>GTGACTGGAGTTCCTTGGCACCCGAGAATTCCA-3’), and 0.02 U/μL Phusion HF polymerase in 30 μL reaction. PCR was performed with an initial denaturation step of 10 s at 98 °C, amplification for 12 cycles (denaturation for 5 s at 98 °C, annealing for 15 s at 62 °C and extension for 15 s at 72°C), and a final extension for 5 min at 72 °C. Amplified cDNAs were isolated by electrophoresis on 2% agarose gel (PippinPrep Reagent Kit, external Marker B) and products of ∼180 to ∼550 nt were collected. cDNA was recovered by ethanol precipitation and resuspended in 13 μL of nuclease-free water. Barcoded libraries were pooled and sequenced on an Illumina NextSeq platform in high output mode using custom primer s1115 (5’-CTACACGTTCAGAGTTCTACAGTCCGACGATC-3’).

### TSS-seq data analysis

#### Mapping

##### TSS-seq data processing

Quality control on TSS sequencing library FASTQ files was performed to remove reads with low quality using fastq_quality_filter in the FASTX-Toolkit (http://hannonlab.cshl.edu/fastx_toolkit/) package with parameters ‘fastq_quality_filter -v -q 20 -p 75’. Cutadapt^114^ was then used to remove the 6 nucleotide 5’ linker with parameter of ‘cutadapt -u 6’. The resulting reads were trimmed from 3’ end to 35 nucleotides long with parameter of ‘cutadapt -l 35 --minimum-length=35’. Trimmed reads were mapped to the *S. cerevisiae* R64-1-1 (SacCer3) genome using Bowtie^115^ with allowance of no more than two mismatches with suppression of non-uniquely mapped reads ‘bowtie -p3 -v2 -m1 -q --sam --un’, reported in sam files. Uniquely mapped reads were then extracted from sam files using SAMtools^116^ and output in bam format ‘samtools view -F 4 -S -b’. Bam files were then sorted and converted into bed files by SAMtools ‘samtools sort -o’, and BEDTools^117^ ‘bedtools bamtobed -cigar’. Customized commands were then used on bed files to identify the genomic coordinate of the 5’ end of each uniquely mapped read ‘awk ‘BEGIN{FS=OFS=“\t” } $6==“+” {$3=$2+1} $6==“-” {$2=$3-1} {print}’’. BEDTools was then used to determine pileup (TSS coverage) across the genome with parameters of ‘bedtools genomecov -g R64.new.genome -i -bg -strand -’ and ‘bedtools genomecov -g R64.new.genome -i -bg -strand +’, resulting in stranded **bedGraph** files. FASTQ files of individual library were directly processed or contacted by strains/mutants to generate begraph files for correlation analysis. For each of 5979 selected yeast promoters, TSS usage was examined within 401-nt wide window, spanning 250 nt upstream and 150 nt downstream of the previously annotated median TSS. Using customized bash and R scripts, TSS coverage from the bedGraph files of a library or a mutant were assigned into the defined windows to generate a 401×5979 TSS **count table**, with each row representing one of the 5979 promoters in the same order as the promoter annotation file, each column represents a promoter position, and the number in each cell representing 5’ ends mapping to that position. These count tables were stored in csv files. Using customized R script and the count table of a library or a mutant, an **expression-spread-median file** containing promoter expression, median TSS position of the promoter, TSS spread of the promoter was generated. The median TSS position was defined as the actual TSS containing the 50th percentile of the promoter window. The spread of TSS, which measures the width of the middle 80% of TSS distribution, was calculated by subtracting positions of 10th percentile and 90th percentile of TSS counts in 401-nt promoter window and adding 1. The positions of 10th percentile and 90th percentile of TSS counts in each promoter window were also stored in this expression-spread-median file. The streamlined codes to generate bedGraph files, the prompter annotation file and the customized scripts to generate count tables and expression-spread-median files can be found on Github account TingtingSsl2, under repository of “Ssl2_scanning”.

##### TSS correlation

TSS coverage data in library-based bedgraph files were used to examine the correlation between TSS libraries by pairwise comparison. A custom R script was used to filter bedgraph files to examine genome positions with greater than two counts in each library. Log_2_ transformed TSS counts at the same genomic location in two examined libraries were plotted for all TSS sites to create a heat scatter plot using the LSD R package^117^ and the Pearson correlation coefficient was calculated. The correlation coefficients deriving from all pairwise comparisons were plotted by a web-based heatmap tool Morpheus (https://software.broadinstitute.org/morpheus/) and clustered by Euclidian distance. Replicates with correlation coefficient greater than 0.85 and the shortest Euclidian distance to each other in the clustering analysis among all the analyzed libraries were recognized as having good sequencing reproducibility and used for downstream analysis.

##### TSS count table and heatmaps

For each of 5979 selected promoters, TSS usage was examined within a 401-nt wide window, spanning 250-nt upstream and 150-nt downstream of the previously annotated median TSS^13^. Using BEDTools and customized R scripts, TSS coverage from the bedgraph files of a library or mutant were assigned into the defined windows to generate a 401×5979 TSS count table, with each row representing one of the 5979 promoters, each column represents a promoter position, and the number in each cell representing 5’ ends mapping to that position. The count table was filtered to keep data from n=4392 promoters with ≥100 sequence reads on average per WT library and used for downstream analyses. The filtered count table was row-normalized to get the relative TSS usage at each promoter position. TSS distribution differences were determined by subtracting normalized WT data (concatenated from both *RPB1* and *SSL2* WT libraries) from normalized mutant data and visualized using heatmaps (Morpheus).

##### TSS metrics

Data analyses for TSS distributions were based on customized R scripts and results were plotted in Graphpad Prism 8 (https://www.graphpad.com/scientific-software/prism/) unless otherwise indicated. The median TSS position was defined as the actual TSS containing the 50th percentile of the TSS distribution and was determined for each promoter. “TSS shift” represents the difference in nucleotide of the median TSS position for each promoter between two libraries or mutants. The distributions of TSS shifts for n=4392 promoters with ≥100 sequence reads on average per WT library or selected promoter classes in each library or mutant were illustrated by both heatmap (Morpheus) and boxplots.

##### TSS spread

The spread of TSS, which measures the width of the middle 80% of TSS distribution, was calculated by subtracting positions of 10th percentile and 90th percentile of TSS reads in 401-nt promoter window and adding 1. TSS spread of selected promoters are shown in boxplot and compared between libraries by performing one-way ANOVA. The differences of TSS spreads between WT and the mutant for selected promoter classes were presented in heatmap (Morpheus) and/or boxplot.

##### TSS-seq data processing

Quality control on TSS sequencing library FASTQ files was performed to remove reads with low quality using fastq_quality_filter in the FASTX-Toolkit (http://hannonlab.cshl.edu/fastx_toolkit/) package with parameters ‘fastq_quality_filter -v -q 20 -p 75’. Cutadapt^114^ was then used to remove the 6 nucleotide 5’ linker with parameter of ‘cutadapt -u 6’. The resulting reads were trimmed from 3’ end to 35 nucleotides long with parameter of ‘cutadapt -l 35 --minimum-length=35’. Trimmed reads were mapped to the *S. cerevisiae* R64-1-1 (SacCer3) genome using Bowtie with allowance of no more than two mismatches with suppression of non-uniquely mapped reads ‘bowtie -p3 -v2 -m1 -q --sam --un’, reported in sam files. Uniquely mapped reads were then extracted from sam files using SAMtools and output in bam format ‘samtools view -F 4 -S -b’. Bam files were then sorted and converted into bed files by SAMtools ‘samtools sort -o’, and BEDTools ‘bedtools bamtobed -cigar’. Customized commands were then used on bed files to identify the genomic coordinate of the 5’ end of each uniquely mapped read ‘awk ‘BEGIN{FS=OFS=“\t” } $6==“+” {$3=$2+1} $6==“-” {$2=$3-1} {print}’’. BEDTools was then used to determine pileup (TSS coverage) across the genome with parameters of ‘bedtools genomecov -g R64.new.genome -i -bg -strand -’ and ‘bedtools genomecov -g R64.new.genome -i -bg -strand +’, resulting in stranded **bedGraph** files. FASTQ files of individual library were directly processed or contacted by strains/mutants to generate begraph files for correlation analysis. For each of 5979 selected yeast promoters, TSS usage was examined within 401-nt wide window, spanning 250 nt upstream and 150 nt downstream of the previously annotated median TSS. Using customized bash and R scripts, TSS coverage from the bedGraph files of a library or a mutant were assigned into the defined windows to generate a 401×5979 TSS **count table**, with each row representing one of the 5979 promoters in the same order as the promoter annotation file, each column represents a promoter position, and the number in each cell representing 5’ ends mapping to that position. These count tables were stored in csv files. Using customized R script and the count table of a library or a mutant, an **expression-spread-median file** containing promoter expression, median TSS position of the promoter, TSS spread of the promoter was generated. The median TSS position was defined as the actual TSS containing the 50th percentile of the promoter window. The spread of TSS, which measures the width of the middle 80% of TSS distribution, was calculated by subtracting positions of 10th percentile and 90th percentile of TSS counts in 401-nt promoter window and adding 1. The positions of 10th percentile and 90th percentile of TSS counts in each promoter window were also stored in this expression-spread-median file. The streamlined codes to generate bedGraph files, the prompter annotation file and the customized scripts to generate count tables and expression-spread-median files can be found on Github repository https://github.com/Kaplan-Lab-Pitt/Ssl2_scanning.

##### ChIP-exo data processing

ChIP-exo data processing was performed as described by Rossi et al. in Nature Communications, 2018 and Qiu et al. in Genome Biology, 2020. Briefly, ChIP-exo libraries were sequenced on a NextSeq 500 in paired-end mode to generate 40 (read1) x 36 bp (read2) reads. Reads passing Q30 quality threshold were then aligned to the sacCer3 genome using the BWA-MEM alignment algorithm (v0.7.9a) with default parameters^118^. After alignment, PCR duplicates were removed using Picard and SAMtools assuming unique combinations of read1 and read2 were PCR duplicates. Using ScriptManager v0.12 (https://github.com/CEGRcode/scriptmanager), BAM files of a library were assigned into two **401×5979 matrices and saved in CDT files**, which stores counts of 5’ position of protein binding on top and bottom strands, respectively. The same as in TSS-seq data analysis, each row of 401×5979 matrix representing one of 5979 promoters and each column representing a position in the 401-nt promoter window. These **401×5979 matrices were also saved in csv format**. Matrices from the same mutant were combined into a single matrix by adding counts in library matrices at the same dimension and saved in csv files. Similar to TSS-seq analysis, the customized R script and the matrix of a library or a mutant were used to generate an **expression-spread-median file** containing promoter expression, median 5’ position of protein binding, the binding site of the spread of the promoter and saved in txt files.

### Availability of data and materials

Genomics datasets generated in the current study are available in the NCBI BioProject and SRA, under the accession numbers of PRJNA681384 and SRP295731, respectively. The processed genomic data files are available in GEO, under the accession number of GSE182792. The streamlined commands to generate TSS-seq bedGraph files, count tables, tables of expression, spread and median TSS can be found at https://github.com/Kaplan-Lab-Pitt/Ssl2_scanning. ChIP-exo and data analysis was performed as described by Rossi *et al*.^119^ and Qiu *et al*.^13^. Source data files are listed in **Additional Table 3**.

## Acknowledgements

The authors thank Chenxi Qiu for R scripts used in statistical analysis of Kruskal-Wallis test with Dunn’s correction and Mann–Whitney U test. We are deeply grateful to Scott Kuerten and Fred Hyde (Illumina) for advice and give of legacy RiboZero reagents for yeast rRNA removal.

## Additional information

BFP has a financial interest in Peconic, LLC, which utilizes the ChIP-exo technology implemented in this study and could potentially benefit from the outcomes of this research. All other authors declare that they have no competing interests. All other authors declare that they have no competing interests.

## Funding

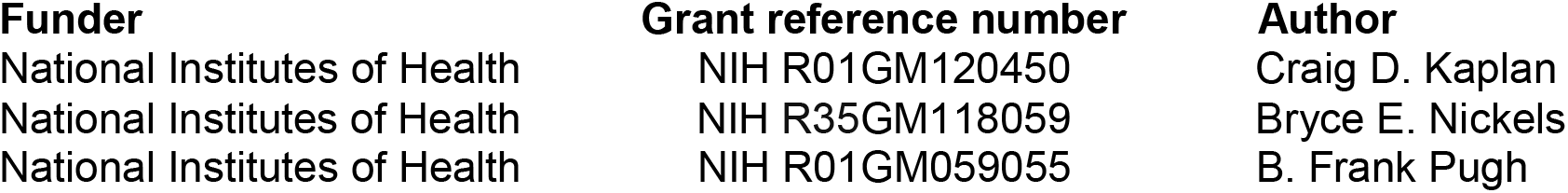

## Author contributions

TZ, Conception and design, Acquisition of data, Analysis and interpretation of data, Drafting and revising the article. IOV, TSS-seq library preparation and sequencing. WKML, ChIP-exo library preparation and processing. BFP, ChIP-Exo infrastructure support. SB, analysis of *SSL2-TAP* strain phenotypes. BEN, TSS-seq methodology and Funding acquisition. CDK, Conception and design, Funding acquisition, Analysis and interpretation of data, Drafting and revising the article.

## Figure Supplements

**Figure 1 – Figure supplement 1.**
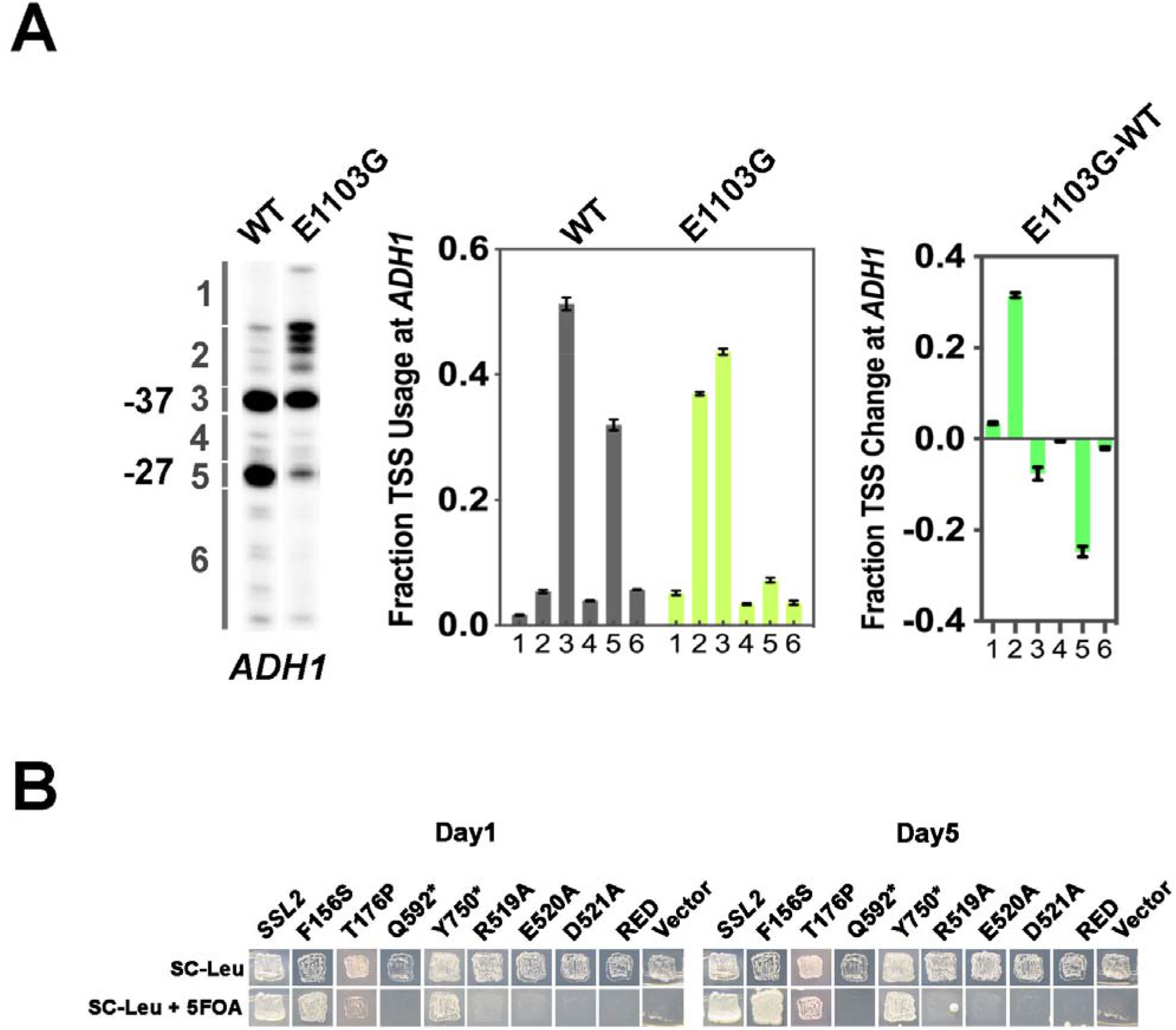
Growth phenotypes of human disease related and RED motif *ssl2* mutants. **(A)** Example of TSS usage quantification at *ADH1*. **Left panel**, primer extension detected TSS usage at *ADH1* in WT and *rpb1* E1103G, an upstream shifting TSS mutant. **Middle panel**, distribution of WT and mutant TSS signal within six *ADH1* bins. Each bin is quantified as the percentage of the total signal in the lane and displayed in a bar graph. Average of ≥ 3 biological replicates +/- standard deviation are shown. **Right panel**, quantitative showing the changes of TSS usage in *rpb1* E1103G compared to WT. **(B)** Viability of *ssl2* alleles with mutations homologous to human disease but inviability of RED motif substitutions (allele marked as “RED” is a triple substitution of R519A/E520A/D521A).

**Figure 2 – Figure supplement 1.**
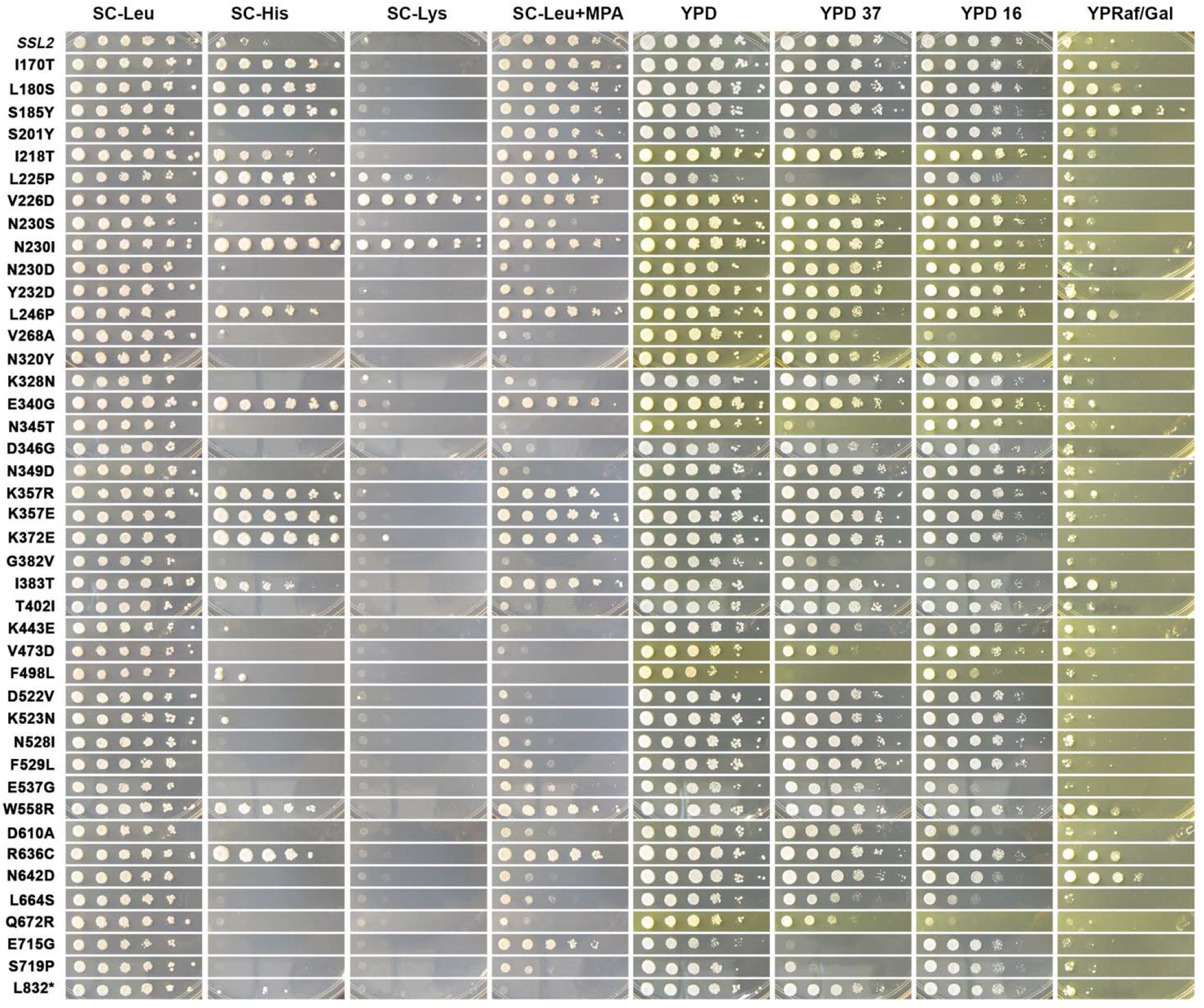
Growth phenotypes of *ssl2* mutants. Growth phenotypes of novel *ssl2* mutants illustrated by spotting of serial dilution (10-fold) of strains on different media. Phenotypes and media are as described in Figure 1. Serial dilution phenotypes shown are representative of at least two independent transformants (biological replicates).

**Figure 2 – Figure supplement 2.**
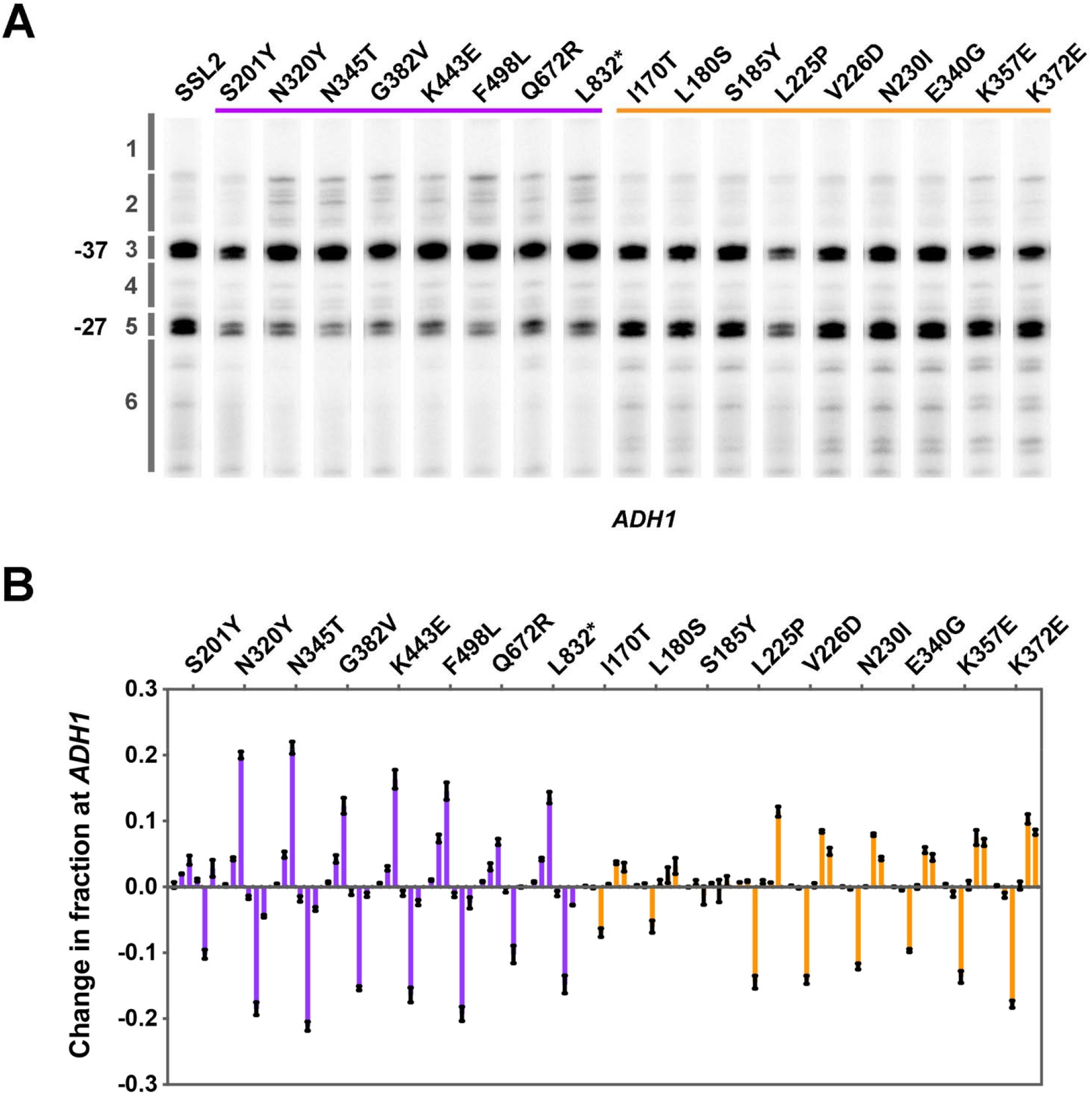
TSS usage of representative *ssl2* single mutants. **(A)** TSS usage for selected *ssl2* alleles at *ADH1* detected by primer extension. The purple bar indicates mutants showing MPA sensitivity (upstream shifting) and the orange bar indicates mutants showing the His^+^ phenotype (downstream shifting). Primer extensions shown are representative of ≥ 3 independent biological replicates. **(B)** Quantification of TSS usage for selected *ssl2* alleles at *ADH1* in (A). Error bars indicate average +/- SD of at least three independent biological replicates.

**Figure 2 – Figure supplement 3.**
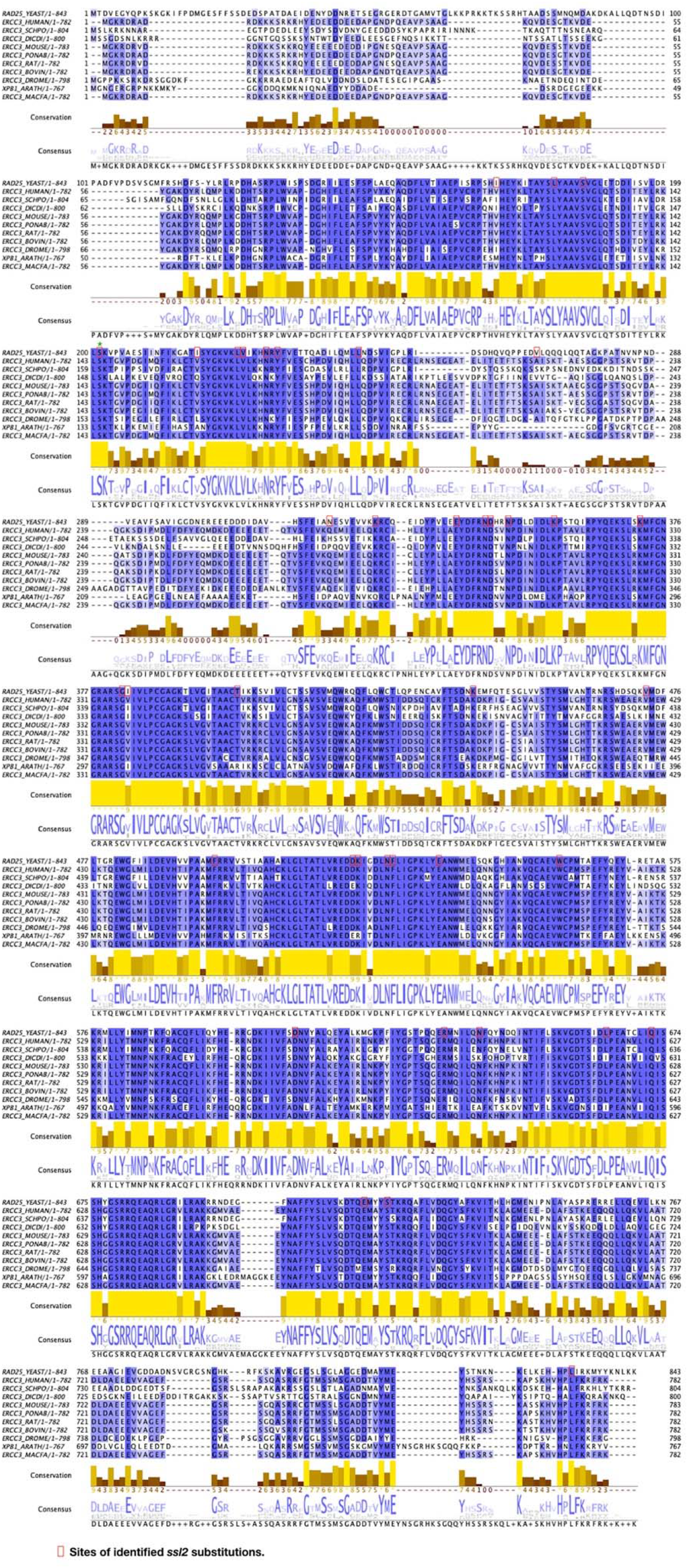
Alignment of Ssl2 homologs illustrating position of substitutions and the conservation of affected residues. Protein sequences of Ssl2 and homologs from a selection of eukaryotes were aligned and displayed in Jalview. Locations of *ssl2* mutations identified in this research were red-boxed on the yeast sequence (“*RAD25_YEAST*”). Locations of *ssl2* mutations were marked with red or green stars if they were annotated in the yeast PIC structure.

**Figure 3 – Figure supplement 1.**
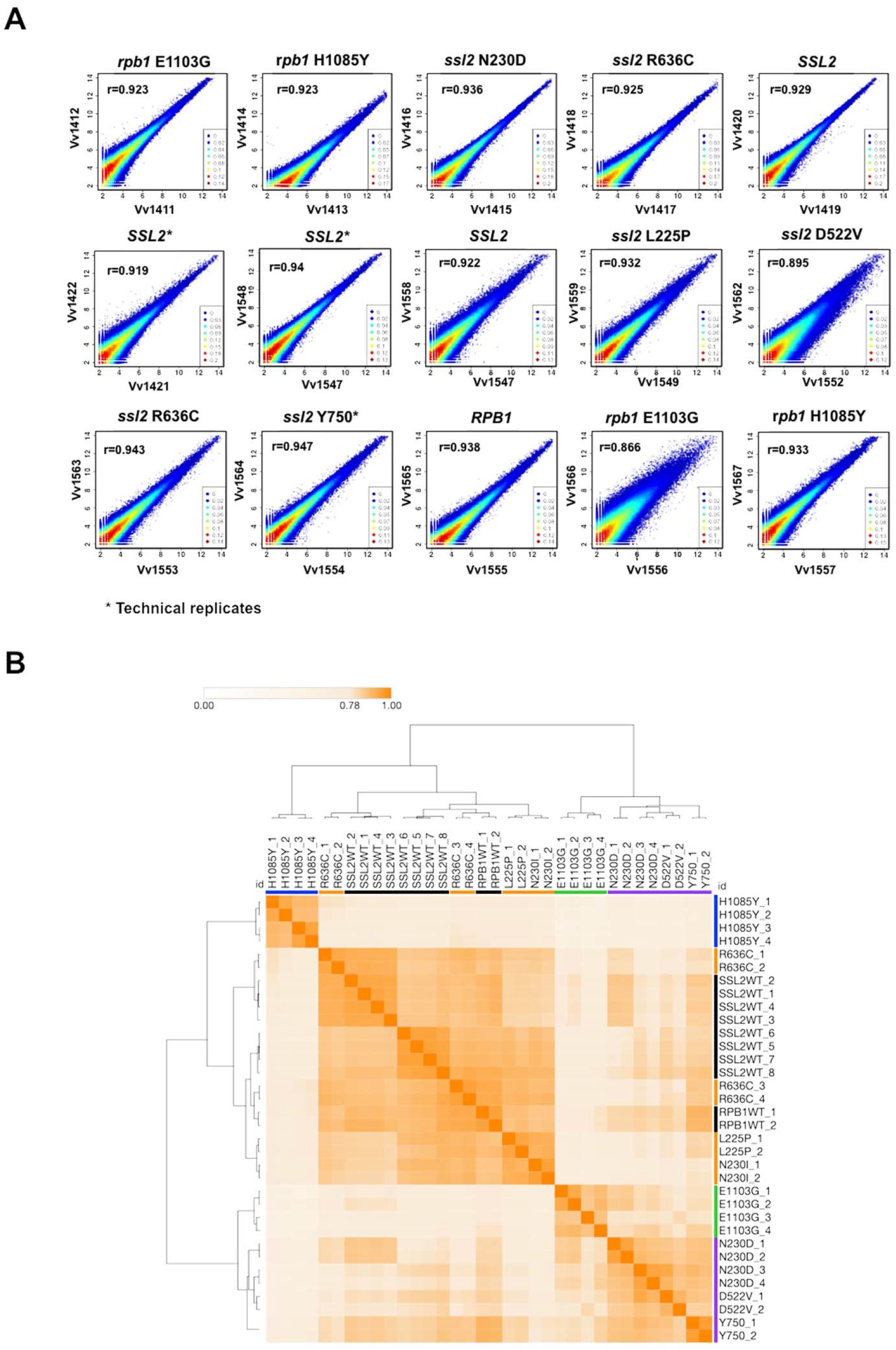
Correlation of read counts between TSS-seq replicates at TSSs across the genome. **(A)** Scatter plot showing the correlation of sequencing reads across all genome positions for all positions ≥3 reads between replicates in *ssl2* and Pol II mutants. **(B)** Matrix heatmap and the hierarchical clustering of Pearson correlation coefficients between TSS-seq libraries from independent biological replicates labeled by substitution and replicate number (H1085Y and E1103G are *rpb1* alleles, all others are *ssl2*). SSL2WT_2,3,4 are technical replicates as are SSL2WT_5,6,7. All other libraries represent biological replicates. Color bars indicate allele class as described in Figure 1.

**Figure 4 – Figure supplement 1.**
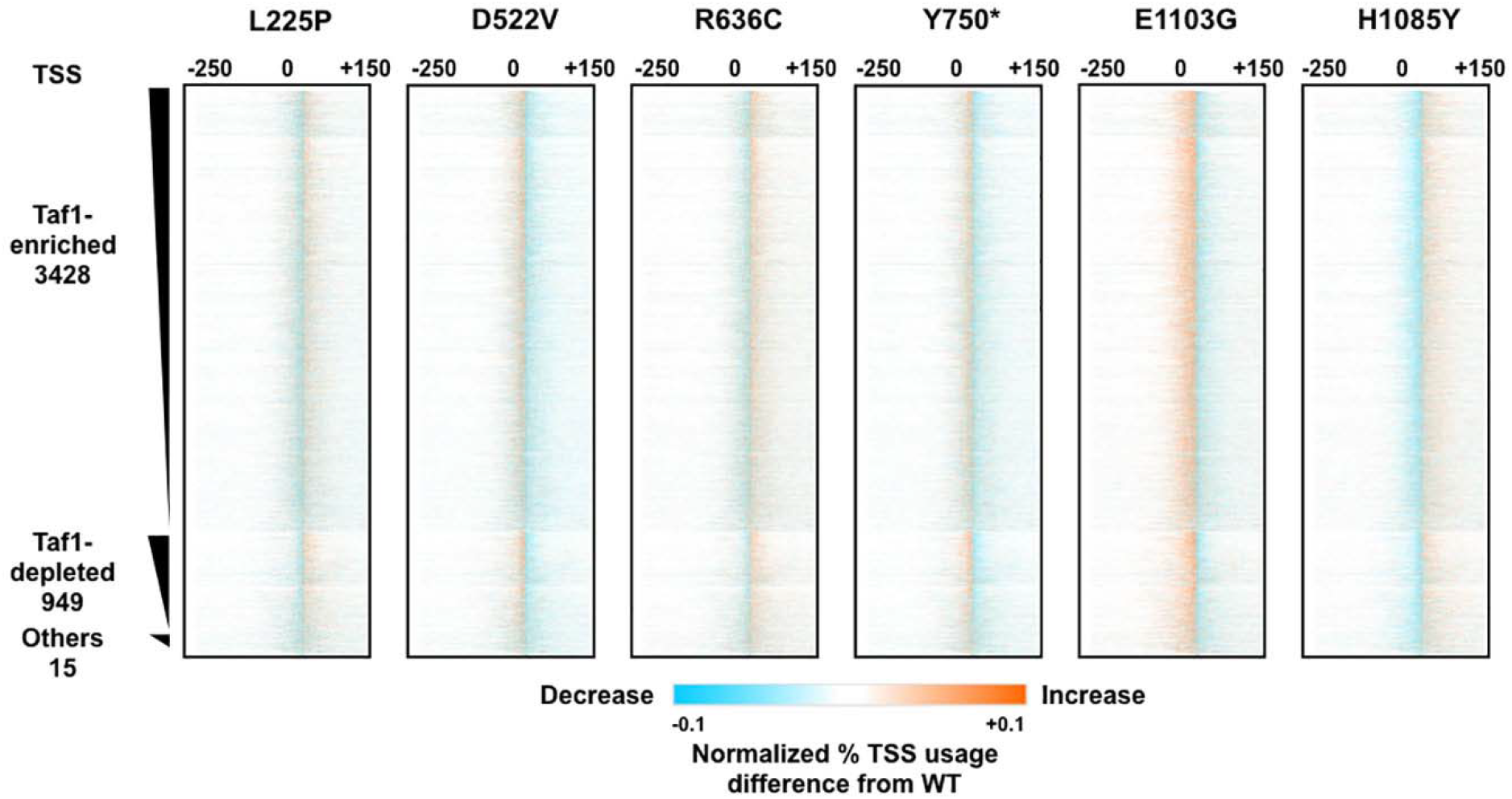
Analysis of TSS shift in *ssl2* and Pol II mutants. Difference heatmaps of TSS distributions between WT and other *ssl2* mutants as described in Figure 4B. The number of points in the figure is lower than the resolution so some averaging will be present. Note that the color scale has been set with sensitive thresholds to have effects be more visually obvious.

**Figure 4 – Figure supplement 2.**
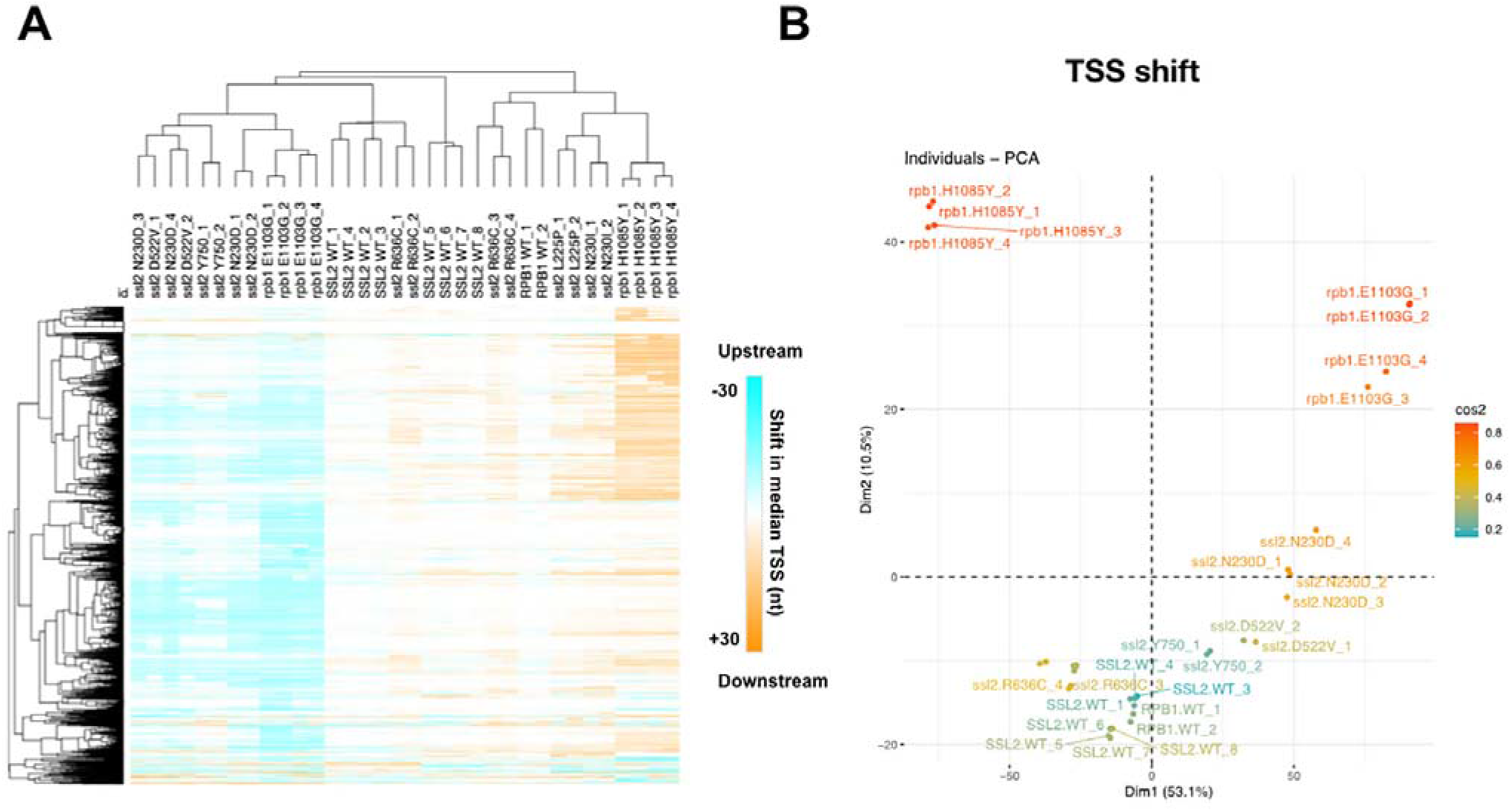
Analysis of TSS shift in *ssl2* and Pol II mutants. **(A)** Heatmap and hierarchical clustering of median TSS shift for *ssl2* or *rpb1* mutant in individual replicates. As described in Figure 4D except performed for individual replicate TSS-seq libraries. SSL2WT_2,3,4 are technical replicates as are SSL2WT_5,6,7. All other libraries represent biological replicates. **(B)** PCA analysis for median TSS shift of *ssl2* and Pol II mutants in individual replicate TSS-seq libraries. TSS-seq libraries are separated into five major classes (*ssl2* upstream and downstream mutants, WT, Pol II upstream and downstream mutants) by the first dimension consistent with direction of shift, and the second dimension consistent with the different genes.

**Figure 5 – Figure supplement 1.**
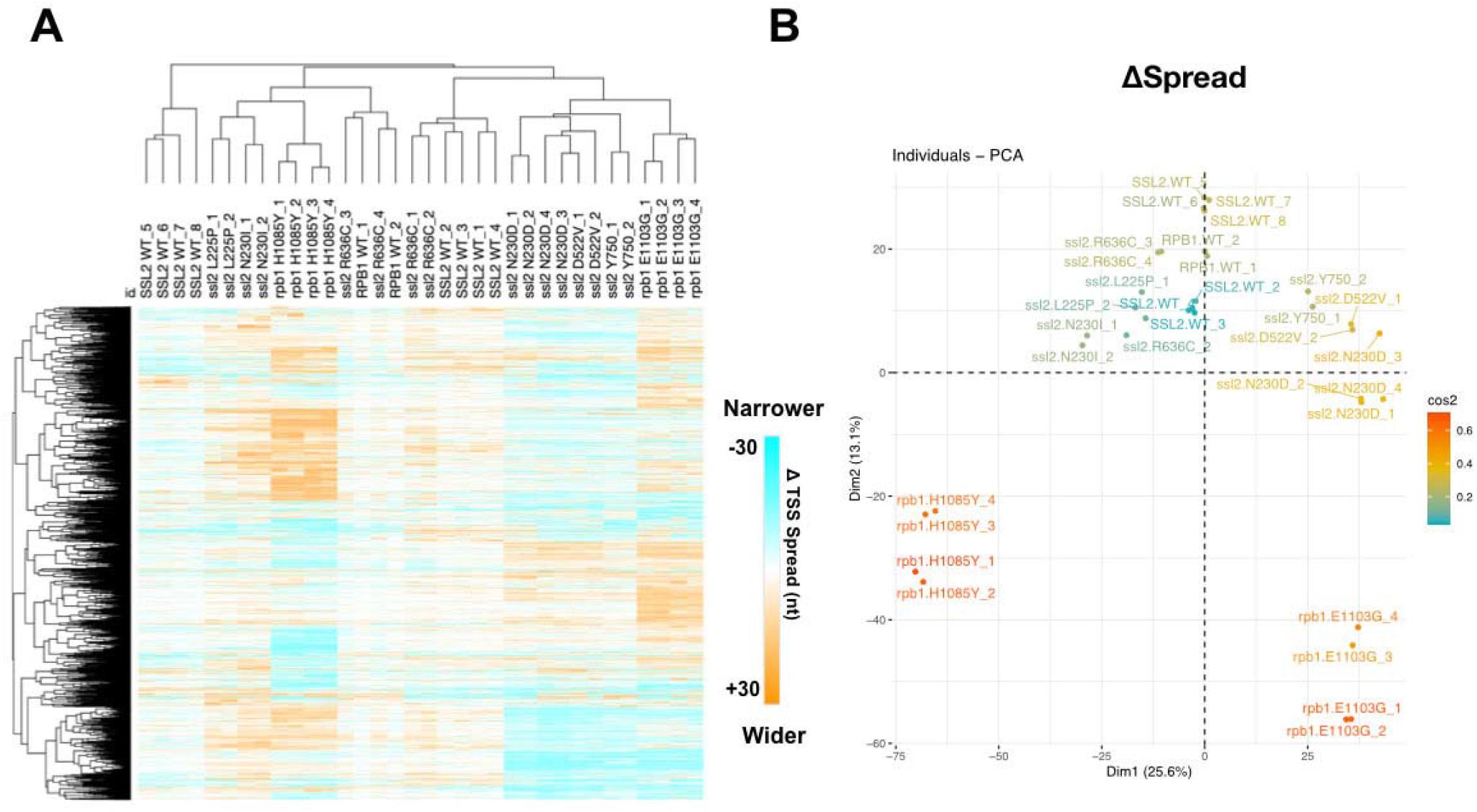
Analysis of TSS spread in *ssl2* and Pol II mutants. **(A)** Heatmap and hierarchical clustering of TSS spread change for *ssl2* or *rpb1* mutant in individual replicate. As described in Figure 5C. SSL2WT_2,3,4 are technical replicates as are SSL2WT_5,6,7. All other libraries represent biological replicates. **(B)** PCA analysis for change of TSS spread in *ssl2* and Pol II individual TSS-seq libraries. TSS-seq libraries are separated into five major classes (*ssl2* upstream and downstream mutants, WT, Pol II upstream and downstream mutants).

**Figure 5 – Figure supplement 2.**
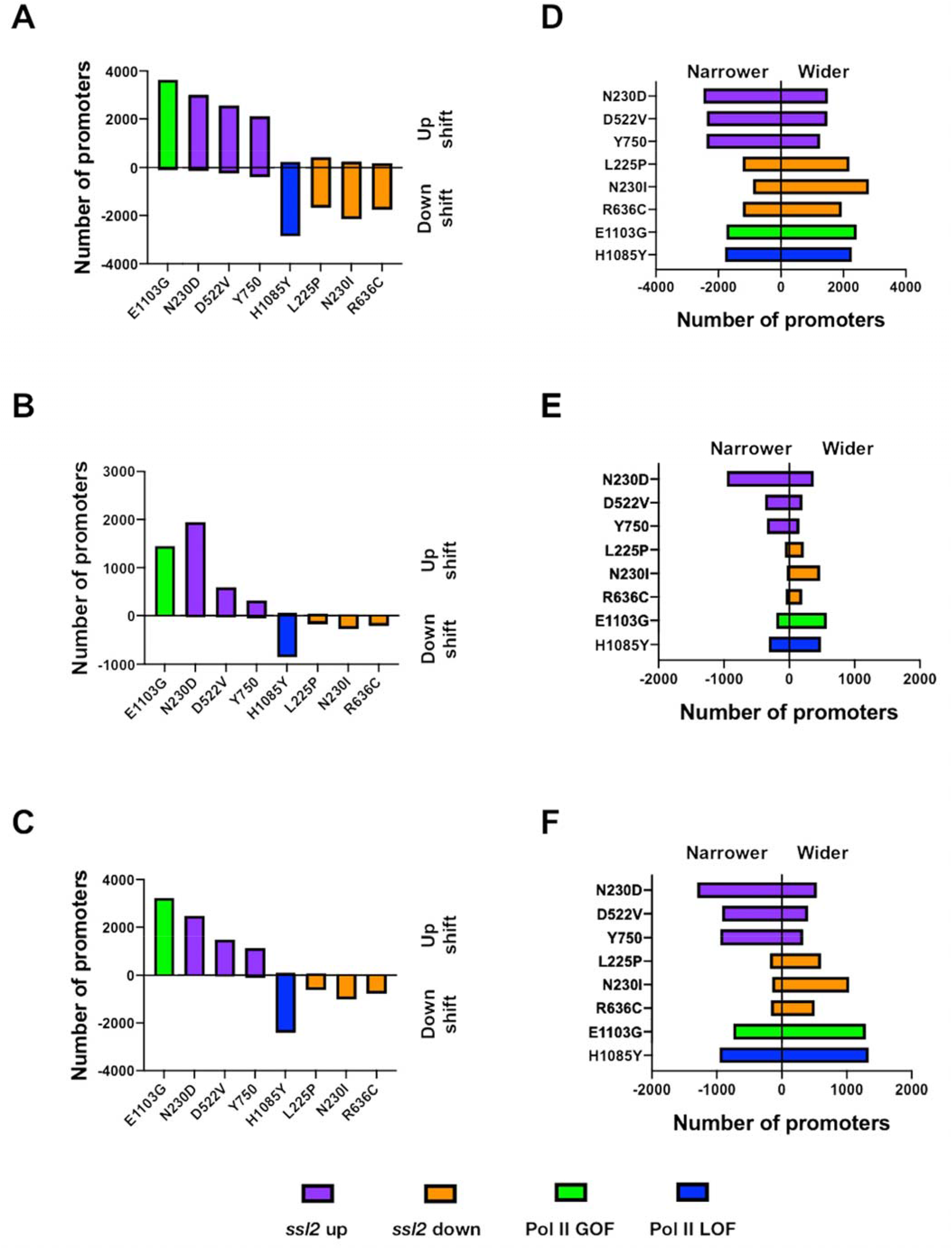
Number of promoters affected by *ssl2* or Pol II mutants. **(A)** Bar plot showing number of promoters that shift median TSS position either upstream or downstream in each mutant for any degree of shift. **(B)** Bar plot indicating number of promoters that shift median TSS position significantly upstream or downstream in each mutant, as determined by Kruskal-Wallis test with Dunn’s correction for multiple comparisons (p≤0.05). The test was performed individually on each promoter and all mutants were taken into account. **(C)** Bar plot showing number of promoters that shift median TSS position significantly upstream or downstream in each mutant as in (B) but determined by Mann-Whitney U test (p≤0.05). **(D-F)** As tested in A-C but examined for change of TSS spread instead of median TSS shift. **(D)** Bar plot showing number of promoters that have narrower or wider TSS spread in each mutant. TSS spread was determined as described in Figure 5A. **(E)** Bar plot indicating number of promoters that have significantly increased or decreased TSS spread in each mutant as determined by Kruskal-Wallis test with Dunn’s correction for multiple comparisons (p≤0.05). The test was also performed individually on each promoter and all the mutants were taken into account. **(F)** Bar plot showing number of promoters that have significantly increased or decreased TSS spread in each mutant as in E but determined by Mann-Whitney U test (p≤0.05).

**Figure 6 – Figure supplement 1.**
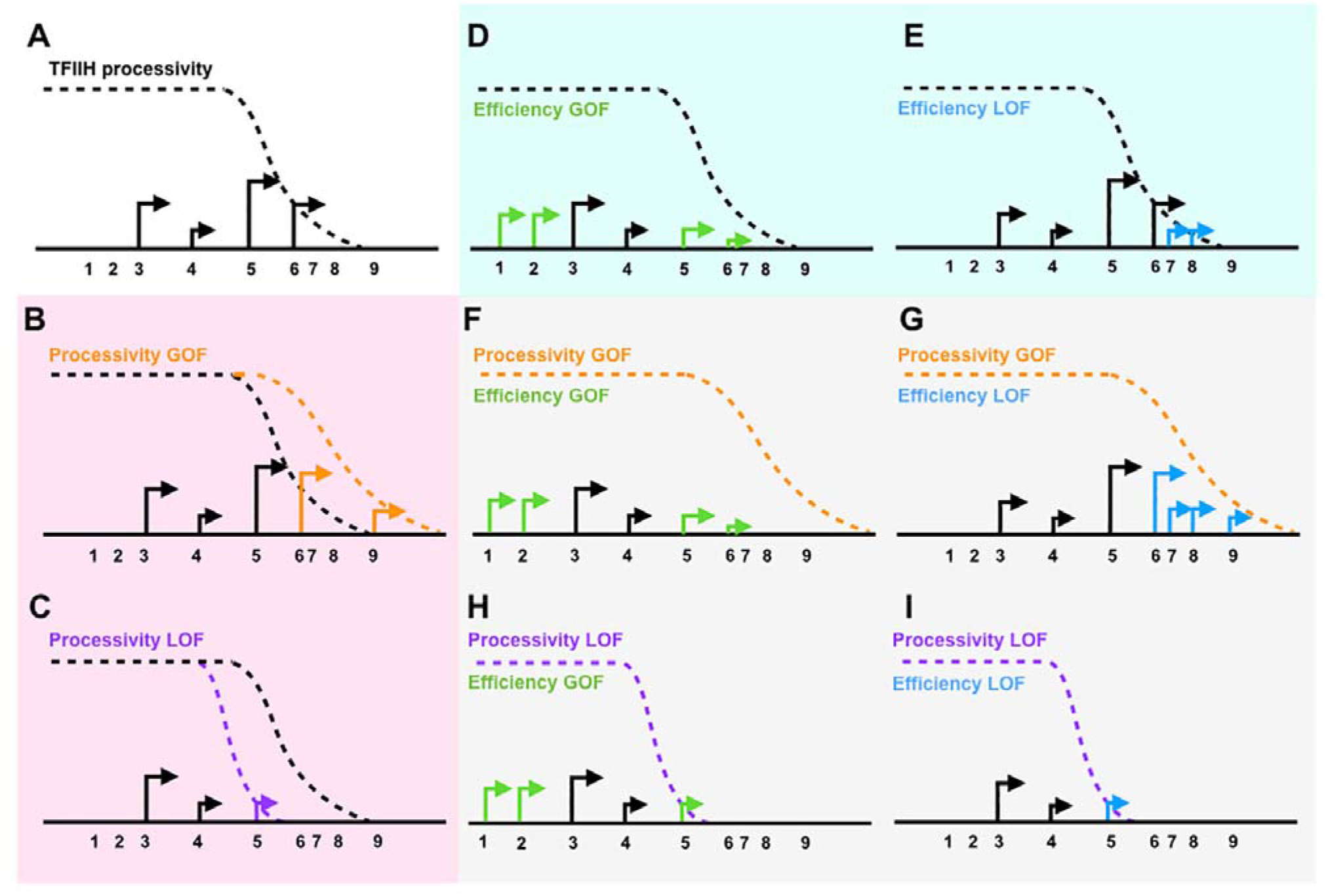
Design of *ssl2* genetic interaction tests with efficiency alleles. **(A)** Schematic showing of a promoter window with TSS. **(B)** Increased processivity is hypothesized to increase TSS usage downstream. **(C)** Decreased processivity is hypothesized to reduce TSS usage downstream. **(D)** Increased efficiency is expected to activate TSS usage upstream. **(E)** Decreased efficiency is expected to increase apparent TSS usage downstream. **(F)** A processivity GOF allele and efficiency GOF double mutant is expected to show efficiency GOF single mutant allele’s effects on activating upstream TSSs (epistasis). **(G)** A processivity GOF and efficiency LOF double mutant is expected to allow or enhance efficiency LOF single allele’s effect on increasing apparent TSS usage downstream (potential additivity). **(H)** A processivity LOF and efficiency GOF double mutant is expected to show efficiency GOF single allele’s effect on activation TSS usage upstream (epistasis). **(I)** A processivity LOF and efficiency LOF double mutant is expected to show processivity LOF single allele’s effect on reducing downstream TSS usage (epistasis).

**Figure 6 – Figure supplement 2.**
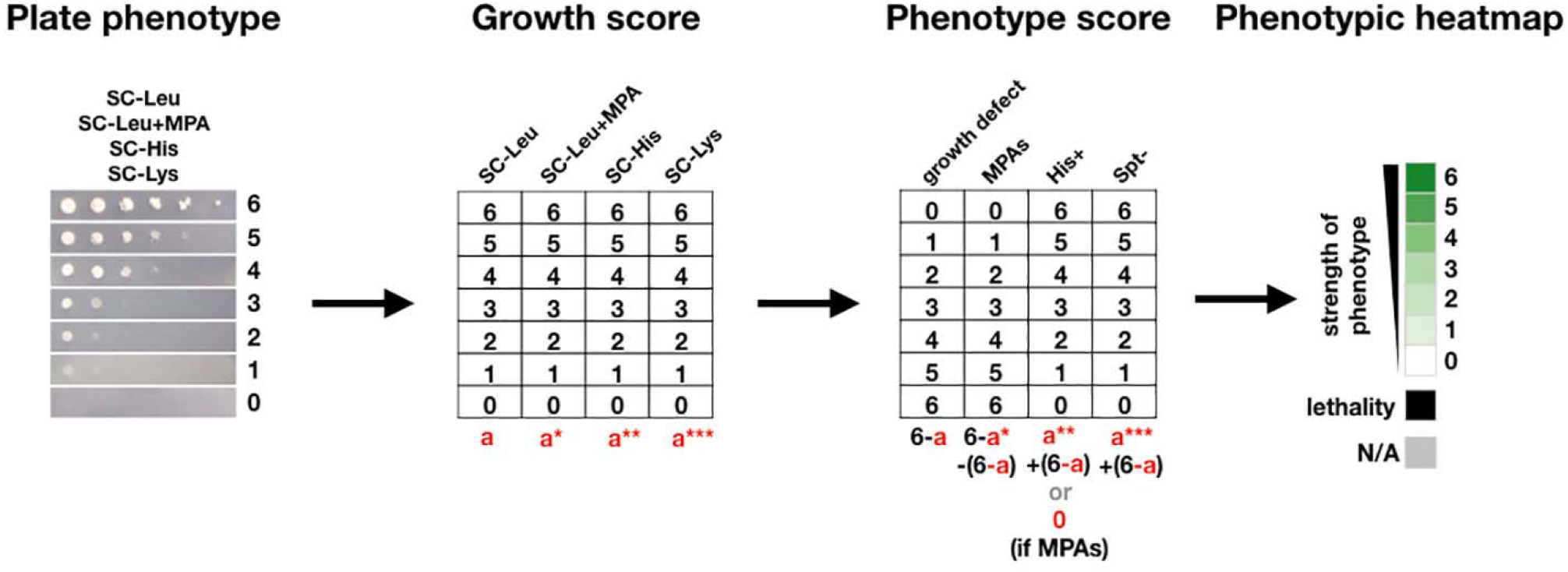
The scoring method used to quantify yeast growth phenotypes and make phenotypic heatmaps.

**Figure 6 – Figure supplement 3:**
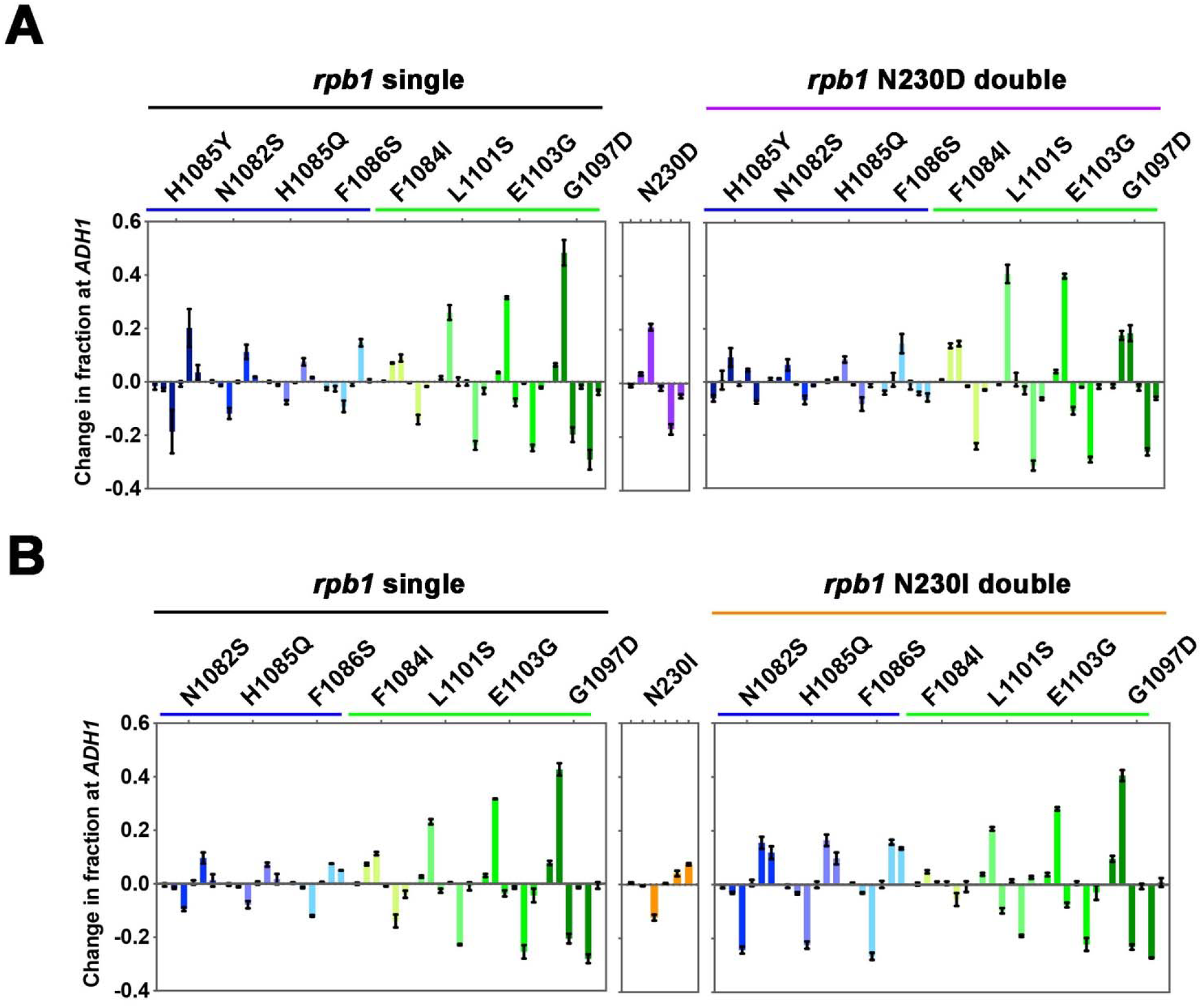
Pol II efficiency alleles are able to increase TSS efficiency within the processivity defined scanning window. **(A)** Quantification of TSS usage at *ADH1* in *rpb1* single mutant, *ssl2* N230D single mutant and the double mutants. TSS usage in single or double mutants is compared to WT TSS usage. Bars are average +/- standard deviation of ≥ 3 independent biological replicates. **(B)** Quantification of TSS usage at *ADH1* in *rpb1* single mutant, *ssl2* N230I single mutant and the double mutants. TSS usage in double mutants is compared to WT TSS usage. Bars are average +/- standard deviation of ≥ 3 independent biological replicates.

**Figure 7 – Figure supplement 1:**
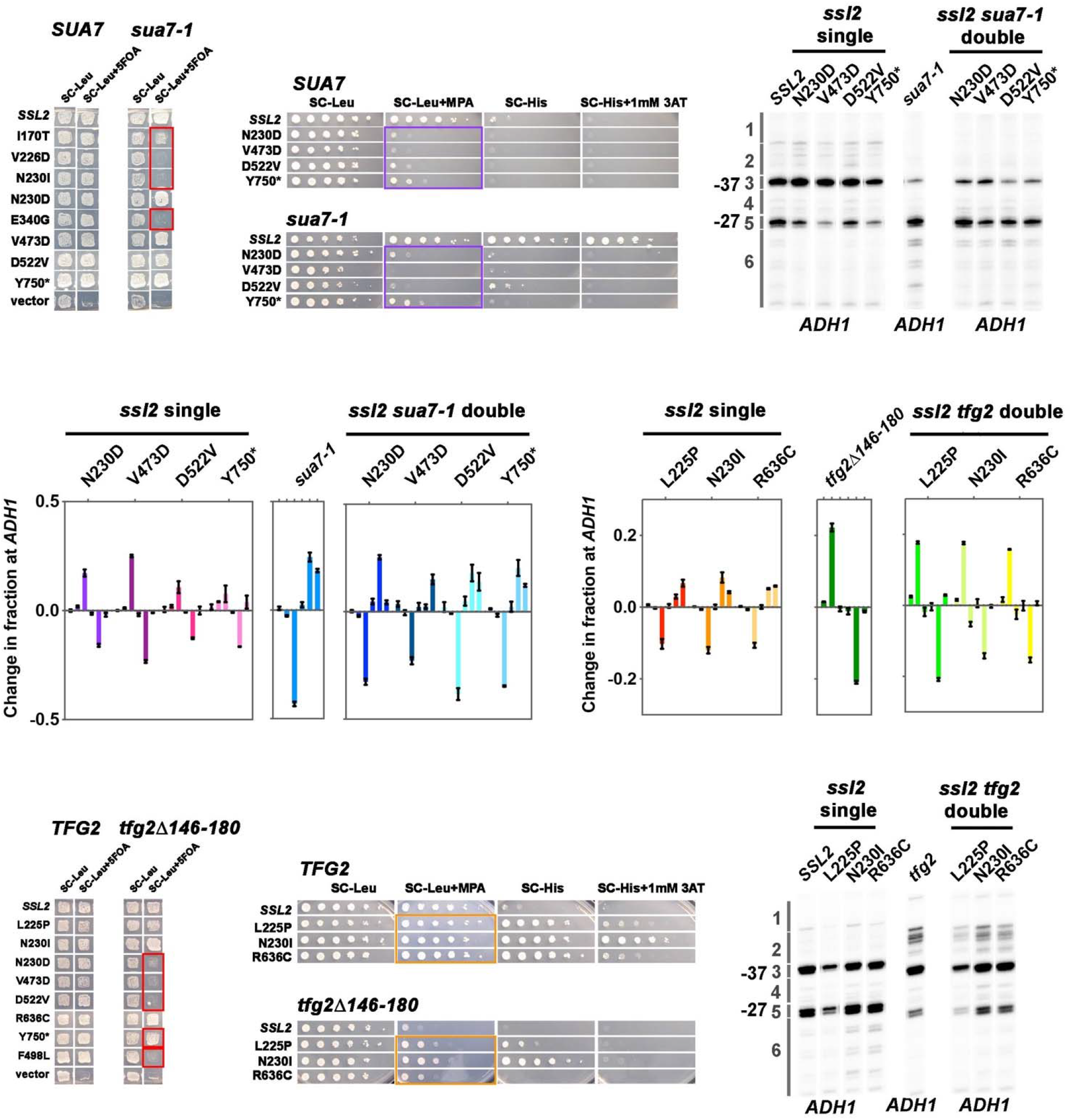
TFIIB and TFIIF alleles show strong and distinct genetic interaction behavior with *ssl2* alleles. **(A)** Lethality is broadly observed between *ssl2* downstream shifting alleles and the *sua7-1* downstream shifting allele. Patch assay showing that six downstream shifting alleles I170T, V226D, N230I, E340G, L225P and R636C, show synthetic lethality phenotypes when combined with *sua7-1* allele (*ssl2* I170T and *sua7-1* double mutant is very sick on 5FOA plate but cannot be propagated to single colonies). Four *ssl2* upstream shifting alleles N230D, V473D, D522V, Y750* show normal growth phenotypes when *sua7-1* is incorporated. One *ssl2* upstream shifting allele F498L showed unique (lethality) genetic interactions compared to other *ssl2* upstream shifting alleles (normal growth). This allele additionally shows lethality with both TFIIF and *sub1*Δ alleles as discussed in the corresponding sections. **(B)** *ssl2* upstream shifting alleles show epistasis with the *sua7-1* downstream shifting allele for MPA^S^ phenotypes. Spot assay shows that four upstream shifting *ssl2* alleles’ strong MPA^S^ phenotypes (N230D, V473D, D522V and Y750*) are almost completely unaffected when *sua7-1* allele, which shows a His^+^ phenotype by itself, is incorporated. Primer extensions shown are representative of ≥ 3 independent biological replicates. **(C)** Double mutants of *ssl2* upstream shifting alleles and *sua7-1* downstream shifting allele shift TSS distribution downstream. Primer extension results show that *ssl2* alleles’ effect on shifting TSS distribution upstream is almost completely reversed to downstream shifting after incorporation of *sua7-1* allele, with only slight additive effect seen in double mutants’ TSS usage. Primer extensions shown are representative of ≥ 3 independent biological replicates. **(D)** Quantification of primer extension detected TSS usage at *ADH1* in *ssl2, sua7-1* single or double mutants. Bars are average +/- standard deviation of ≥ 3 independent biological replicates. **(E)** Lethality is observed between *ssl2* upstream shifting alleles and the *tfg2*Δ*146-180* upstream shifting allele. Patch assay illustrates that upstream shifting alleles N230D, V473D, D522V and F498L show synthetic lethality phenotypes when combined with *tfg2*Δ*146-180* allele (*ssl2* Y750* and *tfg2*Δ*146-180* double mutant shows moderate level of growth on 5FOA plate, however, shows strong sickness when propagated to single colonies). Three *ssl2* downstream shifting alleles L225P, N230I and R636C show normal growth phenotypes when *tfg2*Δ*146-180* is incorporated. (R636C in double mutant shows slight sicker growth phenotype than as single mutant). **(F)** *ssl2* downstream shifting alleles show both epistasis and additive effects with *tfg2*Δ*146-180* downstream shifting allele. Spot assay shows that the double mutants of *ssl2* downstream alleles and *tfg2*Δ*146-180* allele show MPA^S^ phenotypes; however, double mutants’ MPA^S^ phenotypes are not as strong as observed in single *tfg2*Δ*146-180* allele. **(G)** Double mutants of *ssl2* downstream shifting allele and *tfg2*Δ*146-180* upstream shifting allele shift TSS distribution upstream. Primer extension results show that *ssl2* alleles’ effect on shifting TSS distribution upstream is almost completely reversed to upstream shifting after incorporation of *tfg2*Δ*146-180* allele. Primer extensions shown are representative of ≥ 3 independent biological replicates. **(H)** Quantification of primer extension detected TSS usage at *ADH1* in *ssl2, tfg2*Δ*146-180* single or double mutants. Bars are average +/- standard deviation of ≥ 3 independent biological replicates.

**Figure 7 – Figure supplement 2:**
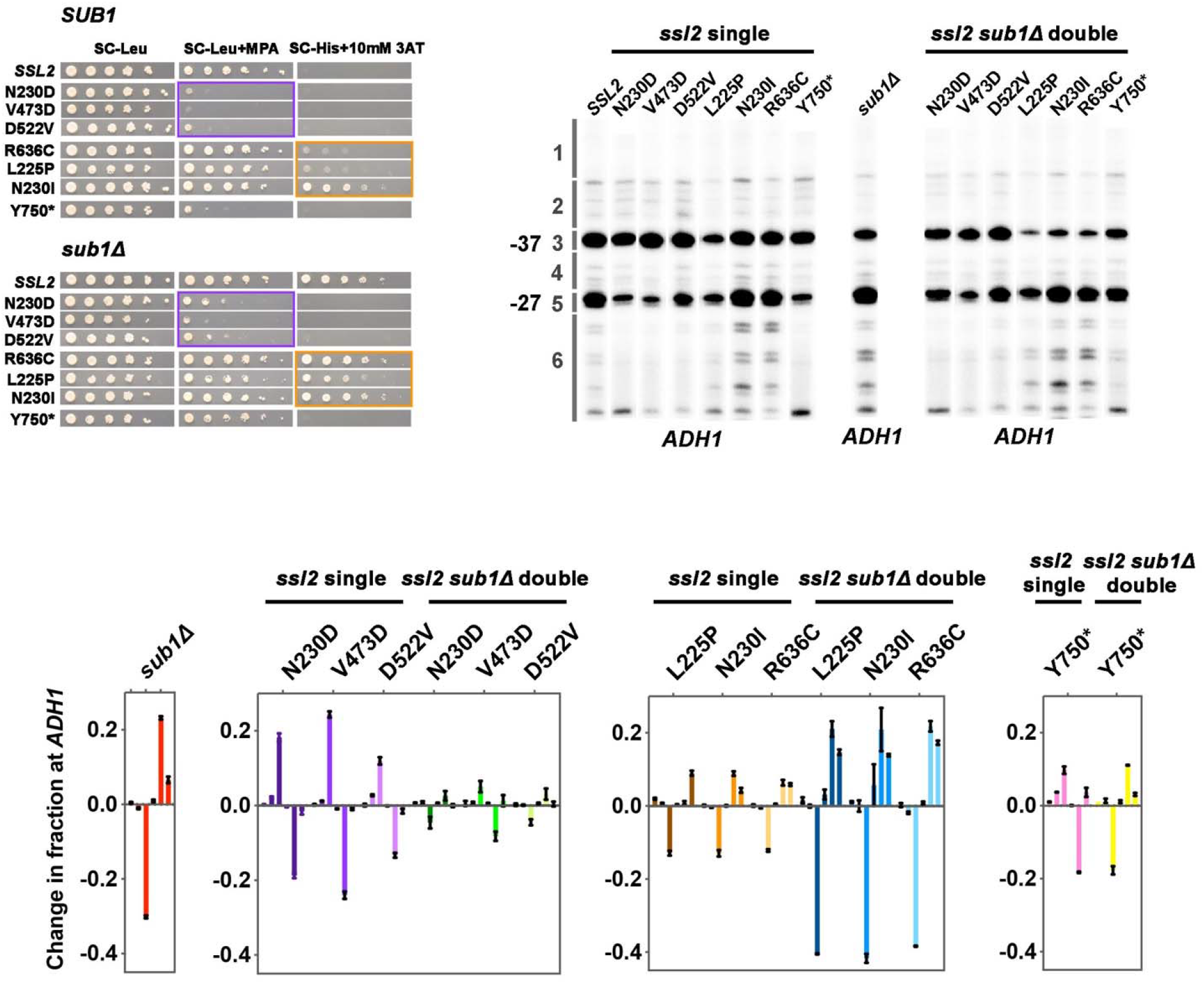
Multiple genetic interactions between *ssl2* and *sub1*Δ alleles. **(A)** Three types of genetic interactions are observed between *ssl2* alleles and *sub1*Δ for the *IMD2* promoter. First, three *ssl2* upstream shifting alleles (N230D, V473D or D522V) are epistatic to *sub1*Δ allele for genetic phenotypes. Spot assay shows that the double mutants of *sub1*Δ (His^+^) and *ssl2* upstream shifting alleles (MPA^S^) show MPA^S^ phenotypes like *ssl2* upstream single alleles. Second, *sub1*Δ is epistatic to *ssl2* allele Y750*. Double mutant of *sub1*Δ (His^+^) and Y750* (MPA^S^) shows a His^+^ phenotype. Third, *sub1*Δ is epistatic to *ssl2* downstream shifting alleles (L225P, N230I, R636C). Double mutants of *sub1*Δ (His^+^) and *ssl2* downstream shifting alleles (His^+^) show His^+^ phenotypes that are not stronger than in *sub1*Δ alone. **(B)** Genetic interactions between *ssl2* and *sub1*Δ alleles at *ADH1* promoter. First, additive effects are observed between *ssl2* upstream shifting alleles and *sub1*Δ. Double mutants of *ssl2* upstream shifting alleles and *sub1*Δ show a compromised TSS distribution. Second, additive effects are also observed between *ssl2* downstream shifting alleles and *sub1*Δ. Double mutants show stronger downstream shift than either single alone. Third, epistasis is observed between *ssl2* Y750* and *sub1*Δ. In the double mutant, *ssl2* Y750*’s effect on shifting TSS distribution upstream is completely reversed to downstream shifting by *sub1*Δ. Primer extensions shown are representative of ≥ 3 independent biological replicates. **(C)** Quantification of primer extension detected TSS usage at *ADH1* in *ssl2, sub1*Δ single or double mutants. Bars are average +/- standard deviation of ≥ 3 independent biological replicates.

**Figure 9 – Figure supplement 1:**
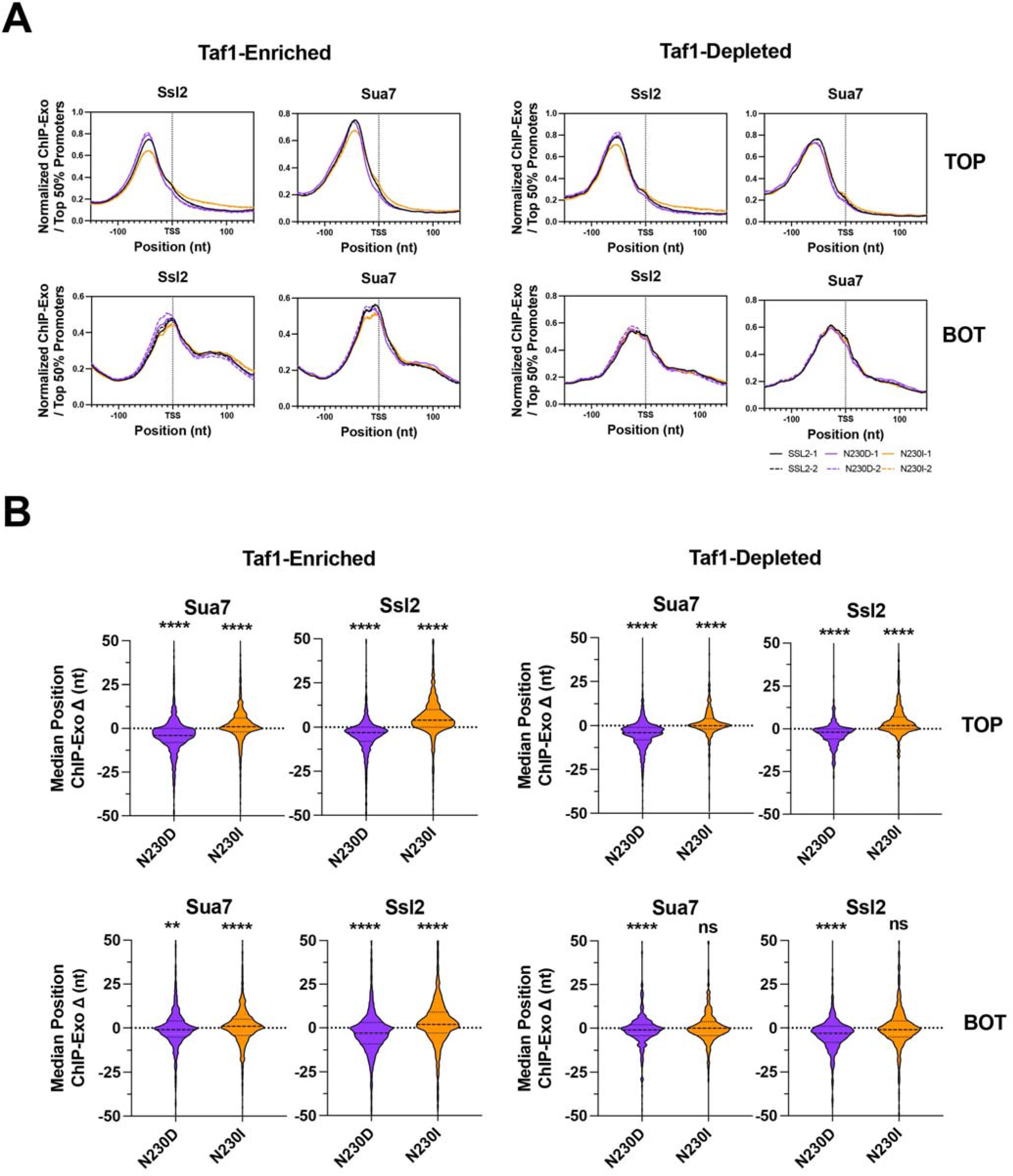
*ssl2* alleles shift PIC-component positioning genome-wide as predicted for mutants altering scanning processivity. **(A)** Positions of PIC components Sua7 and Ssl2 determined by ChIP-exo in WT and *ssl2* mutant strains. Aggregated genome-wide ChIP-exo signals on top (TOP) or bottom (BOT) strand at ± 150 nt TSS centered promoter windows. Enrichment of ChIP-exo signals from the top or bottom strand of Sua7 or Ssl2 sites are shown for two biological replicates per genotype. Curves are LOWESS smoothing of aggregated ChIP-exo reads from the top 50% promoters in WT. **(B)** *ssl2* alleles shift positioning of PIC components Sua7 and Ssl2 at most promoters. Showing is the shift of median Sua7 or Ssl2 position determined by ChIP-exo as illustrated in (A). Statistical analysis shows that the shift of the average median positions of Sua7 and Ssl2 in two replicates are significantly different from zero (Wilcoxon Signed Rank test) at p<0.0001 (****).

**Figure 9 – Figure supplement 2.**
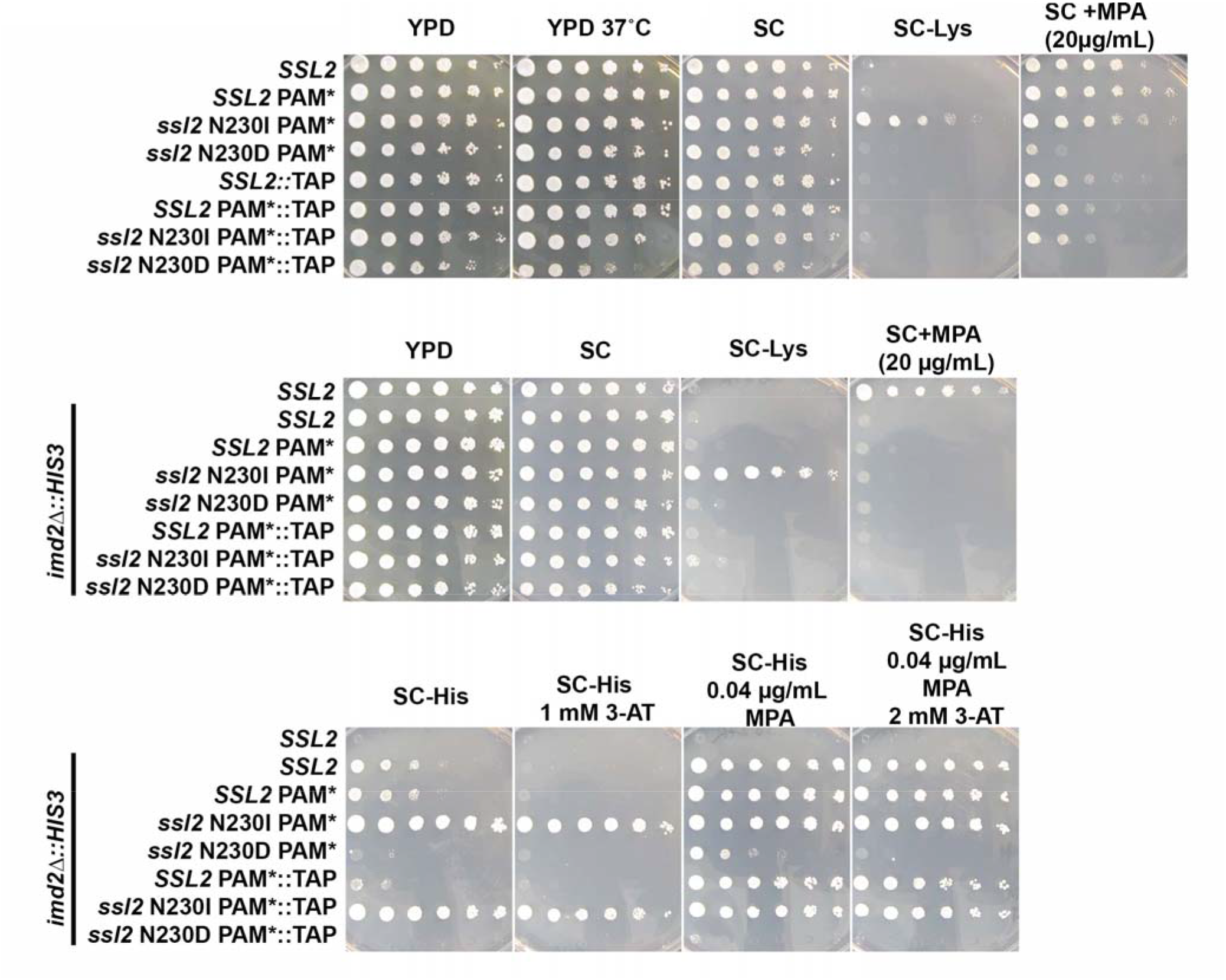
Effects of TAP-tagging *SSL2*. A TAP-tag was integrated at the 3’ end of *SSL2* and CRISPR-Cas9 was used to introduced mutations at codon 230 along with an adjacent silent PAM site mutation to prevent Cas9 recutting. It was discovered that the TAP-tag modulated *SSL2* phenotypes, behaving like a slight loss of function, causing mild MPA sensitivity, enhancing MPA sensitivity of *ssl2* N230D (top row, SC+MPA plate), suppressing the Spt^-^ phenotype of N230I (SC-Lys plates), but having no effect on the His^+^ phenotype of N230I in the background of the *imd2*Δ*::HIS3* initiation reporter (bottom row, SC-His, SC-His 1 mM 3-AT). 3-aminotriazole (3-AT) is a competitive inhibitor of the *HIS3* gene product. All serial dilution phenotyping results are representative of at least two independently constructed strains (biological replicates).

## References

1. Roeder RG: 50+ years of eukaryotic transcription: an expanding universe of factors and mechanisms. Nat Struct Mol Biol 2019, 26(9):783-791. PMCID or (doi): PMC6867066. https://www.ncbi.nlm.nih.gov/pubmed/31439941

2. Arribere JA, Gilbert WV: Roles for transcript leaders in translation and mRNA decay revealed by transcript leader sequencing. Genome Res 2013, 23(6):977-987. PMCID or (doi): 3668365. http://www.ncbi.nlm.nih.gov/pubmed/23580730

3. Rojas-Duran MF, Gilbert WV: Alternative transcription start site selection leads to large differences in translation activity in yeast. Rna 2012, 18(12):2299-2305. PMCID or (doi): 3504680. http://www.ncbi.nlm.nih.gov/pubmed/23105001

4. Malabat C, Feuerbach F, Ma L, Saveanu C, Jacquier A: Quality control of transcription start site selection by nonsense-mediated-mRNA decay. Elife 2015, 4. PMCID or (doi): 4434318. http://www.ncbi.nlm.nih.gov/pubmed/25905671

5. Sample PJ, Wang B, Reid DW, Presnyak V, McFadyen IJ, Morris DR, Seelig G: Human 5’ UTR design and variant effect prediction from a massively parallel translation assay. Nat Biotechnol 2019, 37(7):803-809. PMCID or (doi): PMC7100133. https://www.ncbi.nlm.nih.gov/pubmed/31267113

6. Cuperus JT, Groves B, Kuchina A, Rosenberg AB, Jojic N, Fields S, Seelig G: Deep learning of the regulatory grammar of yeast 5’ untranslated regions from 500,000 random sequences. Genome Res 2017, 27(12):2015-2024. PMCID or (doi): PMC5741052. https://www.ncbi.nlm.nih.gov/pubmed/29097404

7. Akirtava C, McManus CJ: Control of translation by eukaryotic mRNA transcript leaders-Insights from high-throughput assays and computational modeling. Wiley interdisciplinary reviews RNA 2020:e1623. PMCID or (doi): PMC7914273. https://www.ncbi.nlm.nih.gov/pubmed/32869519

8. Lin Y, May GE, Kready H, Nazzaro L, Mao M, Spealman P, Creeger Y, McManus CJ: Impacts of uORF codon identity and position on translation regulation. Nucleic Acids Res 2019, 47(17):9358-9367. PMCID or (doi): PMC6755093. https://www.ncbi.nlm.nih.gov/pubmed/31392980

9. Giardina C, Lis JT: DNA melting on yeast RNA polymerase II promoters. Science 1993, 261(5122):759-762. PMCID or (doi): http://www.ncbi.nlm.nih.gov/pubmed/8342041

10. Kuehner JN, Brow DA: Quantitative analysis of in vivo initiator selection by yeast RNA polymerase II supports a scanning model. J Biol Chem 2006, 281(20):14119-14128. PMCID or (doi): (10.1074/jbc.M601937200) http://www.ncbi.nlm.nih.gov/pubmed/16571719

11. Hampsey M: The Pol II initiation complex: finding a place to start. Nat Struct Mol Biol 2006, 13(7):564-566. PMCID or (doi): (10.1038/nsmb0706-564) http://www.ncbi.nlm.nih.gov/pubmed/16826228

12. Fishburn J, Galburt E, Hahn S: Transcription Start Site Scanning and the Requirement for ATP during Transcription Initiation by RNA Polymerase II. J Biol Chem 2016, 291(25):13040-13047. PMCID or (doi): PMC4933221. https://www.ncbi.nlm.nih.gov/pubmed/27129284

13. Qiu C, Jin H, Vvedenskaya I, Llenas JA, Zhao T, Malik I, Visbisky AM, Schwartz SL, Cui P, Cabart P, Han KH, Lai WKM, Metz RP, Johnson CD, Sze SH, Pugh BF, Nickels BE, Kaplan CD: Universal promoter scanning by Pol II during transcription initiation in Saccharomyces cerevisiae. Genome Biol 2020, 21(1):132. PMCID or (doi): PMC7265651. https://www.ncbi.nlm.nih.gov/pubmed/32487207

14. Tomko EJ, Luyties O, Rimel JK, Tsai C-L, Fuss JO, Fishburn J, Hahn S, Tsutakawa SE, Taatjes DJ, Galburt EA: The role of XPB/Ssl2 dsDNA translocation processivity in transcription-start-site scanning. Journal of Molecular Biology 2021:166813. PMCID or (doi): (https://doi.org/10.1016/j.jmb.2021.166813)http://www.sciencedirect.com/science/article/pii/S0022283621000073

15. Tomko EJ, Fishburn J, Hahn S, Galburt EA: TFIIH generates a six-base-pair open complex during RNAP II transcription initiation and start-site scanning. Nat Struct Mol Biol 2017, 24(12):1139-1145. PMCID or (doi): PMC5741190.https://www.ncbi.nlm.nih.gov/pubmed/29106413

16. Fishburn J, Tomko E, Galburt E, Hahn S: Double-stranded DNA translocase activity of transcription factor TFIIH and the mechanism of RNA polymerase II open complex formation. Proc Natl Acad Sci U S A 2015, 112(13):3961-3966. PMCID or (doi): 4386358. http://www.ncbi.nlm.nih.gov/pubmed/25775526

17. Fazal FM, Meng CA, Murakami K, Kornberg RD, Block SM: Real-time observation of the initiation of RNA polymerase II transcription. Nature 2015, 525(7568):274-277. PMCID or (doi): (10.1038/nature14882) http://www.ncbi.nlm.nih.gov/pubmed/26331540

18. Struhl K: Promoters, activator proteins, and the mechanism of transcriptional initiation in yeast. Cell 1987, 49(3):295-297. PMCID or (doi): http://www.ncbi.nlm.nih.gov/pubmed/2882858

19. Goel S, Krishnamurthy S, Hampsey M: Mechanism of start site selection by RNA polymerase II: interplay between TFIIB and Ssl2/XPB helicase subunit of TFIIH. J Biol Chem 2012, 287(1):557-567. PMCID or (doi): 3249109. http://www.ncbi.nlm.nih.gov/pubmed/22081613

20. Yang C, Ponticelli AS: Evidence that RNA polymerase II and not TFIIB is responsible for the difference in transcription initiation patterns between Saccharomyces cerevisiae and Schizosaccharomyces pombe. Nucleic Acids Res 2012, 40(14):6495-6507. PMCID or (doi): 3413132. http://www.ncbi.nlm.nih.gov/pubmed/22510268

21. Khaperskyy DA, Ammerman ML, Majovski RC, Ponticelli AS: Functions of Saccharomyces cerevisiae TFIIF during transcription start site utilization. Mol Cell Biol 2008, 28(11):3757-3766. PMCID or (doi): 2423299. http://www.ncbi.nlm.nih.gov/pubmed/18362165

22. Pal M, Ponticelli AS, Luse DS: The role of the transcription bubble and TFIIB in promoter clearance by RNA polymerase II. Mol Cell 2005, 19(1):101-110. PMCID or (doi): (10.1016/j.molcel.2005.05.024) http://www.ncbi.nlm.nih.gov/pubmed/15989968

23. Majovski RC, Khaperskyy DA, Ghazy MA, Ponticelli AS: A functional role for the switch 2 region of yeast RNA polymerase II in transcription start site utilization and abortive initiation. J Biol Chem 2005, 280(41):34917-34923. PMCID or (doi): http://www.ncbi.nlm.nih.gov/entrez/query.fcgi?cmd=Retrieve&db=PubMed&dopt=Citation&list_uids=16081422

24. Freire-Picos MA, Krishnamurthy S, Sun ZW, Hampsey M: Evidence that the Tfg1/Tfg2 dimer interface of TFIIF lies near the active center of the RNA polymerase II initiation complex. Nucleic acids research 2005, 33(16):5045-5052. PMCID or (doi): 1201334. http://www.ncbi.nlm.nih.gov/pubmed/16147988

25. Ghazy MA, Brodie SA, Ammerman ML, Ziegler LM, Ponticelli AS: Amino acid substitutions in yeast TFIIF confer upstream shifts in transcription initiation and altered interaction with RNA polymerase II. Mol Cell Biol 2004, 24(24):10975-10985. PMCID or (doi): 533996. http://www.ncbi.nlm.nih.gov/entrez/query.fcgi?cmd=Retrieve&db=PubMed&dopt=Citation&list_uids=15572698

26. Chen BS, Hampsey M: Functional interaction between TFIIB and the Rpb2 subunit of RNA polymerase II: implications for the mechanism of transcription initiation. Mol Cell Biol 2004, 24(9):3983-3991. PMCID or (doi): 387735. http://www.ncbi.nlm.nih.gov/pubmed/15082791

27. Faitar SL, Brodie SA, Ponticelli AS: Promoter-specific shifts in transcription initiation conferred by yeast TFIIB mutations are determined by the sequence in the immediate vicinity of the start sites. Mol Cell Biol 2001, 21(14):4427-4440. PMCID or (doi): 87103. http://www.ncbi.nlm.nih.gov/pubmed/11416123

28. Pappas DL, Jr., Hampsey M: Functional interaction between Ssu72 and the Rpb2 subunit of RNA polymerase II in Saccharomyces cerevisiae. Mol Cell Biol 2000, 20(22):8343-8351. PMCID or (doi): 102141. http://www.ncbi.nlm.nih.gov/pubmed/11046131

29. Wu WH, Pinto I, Chen BS, Hampsey M: Mutational analysis of yeast TFIIB. A functional relationship between Ssu72 and Sub1/Tsp1 defined by allele-specific interactions with TFIIB. Genetics 1999, 153(2):643-652. PMCID or (doi): 1460761. http://www.ncbi.nlm.nih.gov/entrez/query.fcgi?cmd=Retrieve&db=PubMed&dopt=Citation&list_uids=10511545

30. Bangur CS, Faitar SL, Folster JP, Ponticelli AS: An interaction between the N-terminal region and the core domain of yeast TFIIB promotes the formation of TATA-binding protein-TFIIB-DNA complexes. The Journal of biological chemistry 1999, 274(33):23203-23209. PMCID or (doi): http://www.ncbi.nlm.nih.gov/pubmed/10438492

31. Pardee TS, Bangur CS, Ponticelli AS: The N-terminal region of yeast TFIIB contains two adjacent functional domains involved in stable RNA polymerase II binding and transcription start site selection. The Journal of biological chemistry 1998, 273(28):17859-17864. PMCID or (doi): http://www.ncbi.nlm.nih.gov/pubmed/9651390

32. Sun ZW, Tessmer A, Hampsey M: Functional interaction between TFIIB and the Rpb9 (Ssu73) subunit of RNA polymerase II in Saccharomyces cerevisiae. Nucleic acids research 1996, 24(13):2560-2566. PMCID or (doi): 145985. http://www.ncbi.nlm.nih.gov/pubmed/8692696

33. Sun ZW, Hampsey M: Identification of the gene (SSU71/TFG1) encoding the largest subunit of transcription factor TFIIF as a suppressor of a TFIIB mutation in Saccharomyces cerevisiae. Proceedings of the National Academy of Sciences of the United States of America 1995, 92(8):3127-3131. PMCID or (doi): 42118. http://www.ncbi.nlm.nih.gov/pubmed/7724527

34. Pinto I, Wu WH, Na JG, Hampsey M: Characterization of sua7 mutations defines a domain of TFIIB involved in transcription start site selection in yeast. J Biol Chem 1994, 269(48):30569-30573. PMCID or (doi): http://www.ncbi.nlm.nih.gov/entrez/query.fcgi?cmd=Retrieve&db=PubMed&dopt=Citation&list_uids=7982976

35. Berroteran RW, Ware DE, Hampsey M: The sua8 suppressors of Saccharomyces cerevisiae encode replacements of conserved residues within the largest subunit of RNA polymerase II and affect transcription start site selection similarly to sua7 (TFIIB) mutations. Mol Cell Biol 1994, 14(1):226-237. PMCID or (doi): 358373. http://www.ncbi.nlm.nih.gov/pubmed/8264591

36. Pinto I, Ware DE, Hampsey M: The yeast SUA7 gene encodes a homolog of human transcription factor TFIIB and is required for normal start site selection in vivo. Cell 1992, 68(5):977-988. PMCID or (doi): (0092-8674(92)90040-J [pii]) http://www.ncbi.nlm.nih.gov/pubmed/1547497

37. Hampsey M, Na JG, Pinto I, Ware DE, Berroteran RW: Extragenic suppressors of a translation initiation defect in the cyc1 gene of Saccharomyces cerevisiae. Biochimie 1991, 73(12):1445-1455. PMCID or (doi): http://www.ncbi.nlm.nih.gov/pubmed/1666843

38. Knaus R, Pollock R, Guarente L: Yeast SUB1 is a suppressor of TFIIB mutations and has homology to the human co-activator PC4. Embo J 1996, 15(8):1933-1940. PMCID or (doi): 450112. http://www.ncbi.nlm.nih.gov/entrez/query.fcgi?cmd=Retrieve&db=PubMed&dopt=Citation&list_uids=8617240

39. Jin H, Kaplan CD: Relationships of RNA polymerase II genetic interactors to transcription start site usage defects and growth in Saccharomyces cerevisiae. G3 2014, 5(1):21-33. PMCID or (doi): 4291466. http://www.ncbi.nlm.nih.gov/pubmed/25380729

40. Kaplan CD: Basic mechanisms of RNA polymerase II activity and alteration of gene expression in Saccharomyces cerevisiae. Biochim Biophys Acta 2013, 1829(1):39-54. PMCID or (doi): 4026157. http://www.ncbi.nlm.nih.gov/pubmed/23022618

41. Malik I, Qiu C, Snavely T, Kaplan CD: Wide-ranging and unexpected consequences of altered Pol II catalytic activity in vivo. Nucleic Acids Res 2017, 45(8):4431-4451. PMCID or (doi): PMC5416818. https://www.ncbi.nlm.nih.gov/pubmed/28119420

42. Gulyas KD, Donahue TF: SSL2, a suppressor of a stem-loop mutation in the HIS4 leader encodes the yeast homolog of human ERCC-3. Cell 1992, 69(6):1031-1042. PMCID or (doi): (10.1016/0092-8674(92)90621-i) https://www.ncbi.nlm.nih.gov/pubmed/1318786

43. Qiu H, Park E, Prakash L, Prakash S: The Saccharomyces cerevisiae DNA repair gene RAD25 is required for transcription by RNA polymerase II. Genes Dev 1993, 7(11):2161-2171. PMCID or (doi): (10.1101/gad.7.11.2161) https://www.ncbi.nlm.nih.gov/pubmed/7693549

44. Lee BS, Lichtenstein CP, Faiola B, Rinckel LA, Wysock W, Curcio MJ, Garfinkel DJ: Posttranslational inhibition of Ty1 retrotransposition by nucleotide excision repair/transcription factor TFIIH subunits Ssl2p and Rad3p. Genetics 1998, 148(4):1743-1761. PMCID or (doi): PMC1460110. https://www.ncbi.nlm.nih.gov/pubmed/9560391

45. Kuehner JN, Brow DA: Regulation of a eukaryotic gene by GTP-dependent start site selection and transcription attenuation. Mol Cell 2008, 31(2):201-211. PMCID or (doi): (10.1016/j.molcel.2008.05.018) http://www.ncbi.nlm.nih.gov/pubmed/18657503

46. Jenks MH, O’Rourke TW, Reines D: Properties of an intergenic terminator and start site switch that regulate IMD2 transcription in yeast. Mol Cell Biol 2008, 28(12):3883-3893. PMCID or (doi): 2423123. http://www.ncbi.nlm.nih.gov/pubmed/18426909

47. Kaplan CD, Jin H, Zhang IL, Belyanin A: Dissection of Pol II trigger loop function and Pol II activity-dependent control of start site selection in vivo. PLoS Genet 2012, 8(4):e1002627. PMCID or (doi): 3325174. http://www.ncbi.nlm.nih.gov/pubmed/22511879

48. Eichner J, Chen HT, Warfield L, Hahn S: Position of the general transcription factor TFIIF within the RNA polymerase II transcription preinitiation complex. Embo J 2010, 29(4):706-716. PMCID or (doi): 2829161. http://www.ncbi.nlm.nih.gov/pubmed/20033062

49. Kaplan CD, Larsson KM, Kornberg RD: The RNA polymerase II trigger loop functions in substrate selection and is directly targeted by alpha-amanitin. Mol Cell 2008, 30(5):547-556. PMCID or (doi): 2475549. http://www.ncbi.nlm.nih.gov/pubmed/18538653

50. Braberg H, Jin H, Moehle EA, Chan YA, Wang S, Shales M, Benschop JJ, Morris JH, Qiu C, Hu F, Tang LK, Fraser JS, Holstege FC, Hieter P, Guthrie C, Kaplan CD, Krogan NJ: From structure to systems: high-resolution, quantitative genetic analysis of RNA polymerase II. Cell 2013, 154(4):775-788. PMCID or (doi): 3932829. http://www.ncbi.nlm.nih.gov/pubmed/23932120

51. Oh KS, Khan SG, Jaspers NG, Raams A, Ueda T, Lehmann A, Friedmann PS, Emmert S, Gratchev A, Lachlan K, Lucassan A, Baker CC, Kraemer KH: Phenotypic heterogeneity in the XPB DNA helicase gene (ERCC3): xeroderma pigmentosum without and with Cockayne syndrome. Hum Mutat 2006, 27(11):1092-1103. PMCID or (doi): (10.1002/humu.20392) https://www.ncbi.nlm.nih.gov/pubmed/16947863

52. Cleaver JE, Thompson LH, Richardson AS, States JC: A summary of mutations in the UV-sensitive disorders: xeroderma pigmentosum, Cockayne syndrome, and trichothiodystrophy. Hum Mutat 1999, 14(1):9-22. PMCID or (doi): (10.1002/(SICI)1098-1004(1999)14:1<9::AID-HUMU2>3.0.CO;2-6) https://www.ncbi.nlm.nih.gov/pubmed/10447254

53. Weeda G, Eveno E, Donker I, Vermeulen W, Chevallier-Lagente O, Taieb A, Stary A, Hoeijmakers JH, Mezzina M, Sarasin A: A mutation in the XPB/ERCC3 DNA repair transcription gene, associated with trichothiodystrophy. Am J Hum Genet 1997, 60(2):320-329. PMCID or (doi): PMC1712398. https://www.ncbi.nlm.nih.gov/pubmed/9012405

54. Weeda G, Ma LB, van Ham RC, van der Eb AJ, Hoeijmakers JH: Structure and expression of the human XPBC/ERCC-3 gene involved in DNA repair disorders xeroderma pigmentosum and Cockayne’s syndrome. Nucleic Acids Res 1991, 19(22):6301-6308. PMCID or (doi): PMC329143. https://www.ncbi.nlm.nih.gov/pubmed/1956789

55. Vermeulen W, Scott RJ, Rodgers S, Muller HJ, Cole J, Arlett CF, Kleijer WJ, Bootsma D, Hoeijmakers JH, Weeda G: Clinical heterogeneity within xeroderma pigmentosum associated with mutations in the DNA repair and transcription gene ERCC3. Am J Hum Genet 1994, 54(2):191-200. PMCID or (doi): PMC1918172. https://www.ncbi.nlm.nih.gov/pubmed/8304337

56. Sweder KS, Hanawalt PC: The COOH terminus of suppressor of stem loop (SSL2/RAD25) in yeast is essential for overall genomic excision repair and transcription-coupled repair. J Biol Chem 1994, 269(3):1852-1857. PMCID or (doi): https://www.ncbi.nlm.nih.gov/pubmed/8294433

57. Weeda G, van Ham RC, Vermeulen W, Bootsma D, van der Eb AJ, Hoeijmakers JH: A presumed DNA helicase encoded by ERCC-3 is involved in the human repair disorders xeroderma pigmentosum and Cockayne’s syndrome. Cell 1990, 62(4):777-791. PMCID or (doi): (10.1016/0092-8674(90)90122-u) https://www.ncbi.nlm.nih.gov/pubmed/2167179

58. Simchen G, Winston F, Styles CA, Fink GR: Ty-mediated gene expression of the LYS2 and HIS4 genes of Saccharomyces cerevisiae is controlled by the same SPT genes. Proc Natl Acad Sci U S A 1984, 81(8):2431-2434. PMCID or (doi): 345074. http://www.ncbi.nlm.nih.gov/entrez/query.fcgi?cmd=Retrieve&db=PubMed&dopt=Citation&list_uids=6326126

59. Kaplan CD, Holland MJ, Winston F: Interaction between transcription elongation factors and mRNA 3’-end formation at the Saccharomyces cerevisiae GAL10-GAL7 locus. J Biol Chem 2005, 280(2):913-922. PMCID or (doi): (10.1074/jbc.M411108200) http://www.ncbi.nlm.nih.gov/pubmed/15531585

60. Greger IH, Proudfoot NJ: Poly(A) signals control both transcriptional termination and initiation between the tandem GAL10 and GAL7 genes of Saccharomyces cerevisiae. Embo J 1998, 17(16):4771-4779. PMCID or (doi): http://www.ncbi.nlm.nih.gov/entrez/query.fcgi?cmd=Retrieve&db=PubMed&dopt=Citation&list_uids=9707436

61. Schilbach S, Aibara S, Dienemann C, Grabbe F, Cramer P: Structure of RNA polymerase II pre-initiation complex at 2.9 A defines initial DNA opening. Cell 2021, 184(15):4064-4072 e4028. PMCID or (doi): (10.1016/j.cell.2021.05.012) https://www.ncbi.nlm.nih.gov/pubmed/34133942

62. Luo J, Cimermancic P, Viswanath S, Ebmeier CC, Kim B, Dehecq M, Raman V, Greenberg CH, Pellarin R, Sali A, Taatjes DJ, Hahn S, Ranish J: Architecture of the Human and Yeast General Transcription and DNA Repair Factor TFIIH. Mol Cell 2015, 59(5):794-806. PMCID or (doi): 4560838. http://www.ncbi.nlm.nih.gov/pubmed/26340423

63. He Y, Yan C, Fang J, Inouye C, Tjian R, Ivanov I, Nogales E: Near-atomic resolution visualization of human transcription promoter opening. Nature 2016, 533(7603):359-365. PMCID or (doi): PMC4940141. https://www.ncbi.nlm.nih.gov/pubmed/27193682

64. Greber BJ, Toso DB, Fang J, Nogales E: The complete structure of the human TFIIH core complex. Elife 2019, 8. PMCID or (doi): PMC6422496. https://www.ncbi.nlm.nih.gov/pubmed/30860024

65. Jawhari A, Laine JP, Dubaele S, Lamour V, Poterszman A, Coin F, Moras D, Egly JM: p52 Mediates XPB function within the transcription/repair factor TFIIH. J Biol Chem 2002, 277(35):31761-31767. PMCID or (doi): (10.1074/jbc.M203792200) https://www.ncbi.nlm.nih.gov/pubmed/12080057

66. Coin F, Oksenych V, Egly JM: Distinct roles for the XPB/p52 and XPD/p44 subcomplexes of TFIIH in damaged DNA opening during nucleotide excision repair. Mol Cell 2007, 26(2):245-256. PMCID or (doi): (10.1016/j.molcel.2007.03.009) https://www.ncbi.nlm.nih.gov/pubmed/17466626

67. Vvedenskaya IO, Goldman SR, Nickels BE: Preparation of cDNA libraries for high-throughput RNA sequencing analysis of RNA 5’ ends. Methods Mol Biol 2015, 1276:211-228. PMCID or (doi): 4349364. http://www.ncbi.nlm.nih.gov/pubmed/25665566

68. Rhee HS, Pugh BF: Genome-wide structure and organization of eukaryotic pre-initiation complexes. Nature 2012, 483(7389):295-301. PMCID or (doi): 3306527. http://www.ncbi.nlm.nih.gov/pubmed/22258509

69. Zhao T, Vvedenskaya IO, Lai WKM, Basu S, Pugh BF, Nickels BE, Kaplan CD: Ssl2/TFIIH function in Transcription Start Site Scanning by RNA Polymerase II in Saccharomyces cerevisiae. bioRxiv 2021:2021.2005.2005.442816. PMCID or (doi): (10.1101/2021.05.05.442816) http://biorxiv.org/content/early/2021/05/05/2021.05.05.442816.abstract

70. Garavis M, Calvo O: Sub1/PC4, a multifaceted factor: from transcription to genome stability. Curr Genet 2017, 63(6):1023-1035. PMCID or (doi): (10.1007/s00294-017-0715-6) https://www.ncbi.nlm.nih.gov/pubmed/28567479

71. Calvo O: Sub1 and RNAPII, until termination does them part. Transcription 2018, 9(1):52-60. PMCID or (doi): PMC5791812. https://www.ncbi.nlm.nih.gov/pubmed/28853990

72. Sikorski TW, Ficarro SB, Holik J, Kim T, Rando OJ, Marto JA, Buratowski S: Sub1 and RPA associate with RNA polymerase II at different stages of transcription. Mol Cell 2011, 44(3):397-409. PMCID or (doi): PMC3227220. https://www.ncbi.nlm.nih.gov/pubmed/22055186

73. Lada AG, Kliver SF, Dhar A, Polev DE, Masharsky AE, Rogozin IB, Pavlov YI: Disruption of Transcriptional Coactivator Sub1 Leads to Genome-Wide Re-distribution of Clustered Mutations Induced by APOBEC in Active Yeast Genes. PLoS Genet 2015, 11(5):e1005217. PMCID or (doi): PMC4420506. https://www.ncbi.nlm.nih.gov/pubmed/25941824

74. Koyama H, Sumiya E, Nagata M, Ito T, Sekimizu K: Transcriptional repression of the IMD2 gene mediated by the transcriptional co-activator Sub1. Genes Cells 2008, 13(11):1113-1126. PMCID or (doi): (GTC1229 [pii] 10.1111/j.1365-2443.2008.01229.x) http://www.ncbi.nlm.nih.gov/entrez/query.fcgi?cmd=Retrieve&db=PubMed&dopt=Citation&list_uids=18823333

75. Cabart P, Jin H, Li L, Kaplan CD: Activation and reactivation of the RNA polymerase II trigger loop for intrinsic RNA cleavage and catalysis. Transcription 2014, 5(3):e28869. PMCID or (doi): PMC4574878. https://www.ncbi.nlm.nih.gov/pubmed/25764335

76. Sainsbury S, Niesser J, Cramer P: Structure and function of the initially transcribing RNA polymerase II-TFIIB complex. Nature 2013, 493(7432):437-440. PMCID or (doi): (10.1038/nature11715) http://www.ncbi.nlm.nih.gov/pubmed/23151482

77. Nogales E, Greber BJ: High-resolution cryo-EM structures of TFIIH and their functional implications. Curr Opin Struct Biol 2019, 59:188-194. PMCID or (doi): PMC6951423. https://www.ncbi.nlm.nih.gov/pubmed/31600675

78. Abdella R, Talyzina A, Chen S, Inouye CJ, Tjian R, He Y: Structure of the human Mediator-bound transcription preinitiation complex. Science 2021. PMCID or (doi): (10.1126/science.abg3074) https://www.ncbi.nlm.nih.gov/pubmed/33707221

79. Chen X, Qi Y, Wu Z, Wang X, Li J, Zhao D, Hou H, Li Y, Yu Z, Liu W, Wang M, Ren Y, Li Z, Yang H, Xu Y: Structural insights into preinitiation complex assembly on core promoters. Science 2021:eaba8490. PMCID or (doi): (10.1126/science.aba8490) http://science.sciencemag.org/content/early/2021/03/31/science.aba8490.abstract

80. Rengachari S, Schilbach S, Aibara S, Dienemann C, Cramer P: Structure of human Mediator-RNA polymerase II pre-initiation complex. Nature 2021. PMCID or (doi): (10.1038/s41586-021-03555-7) https://www.ncbi.nlm.nih.gov/pubmed/33902108

81. Alekseev S, Nagy Z, Sandoz J, Weiss A, Egly JM, Le May N, Coin F: Transcription without XPB Establishes a Unified Helicase-Independent Mechanism of Promoter Opening in Eukaryotic Gene Expression. Mol Cell 2017, 65(3):504-514 e504. PMCID or (doi): (10.1016/j.molcel.2017.01.012) https://www.ncbi.nlm.nih.gov/pubmed/28157507

82. Lin YC, Choi WS, Gralla JD: TFIIH XPB mutants suggest a unified bacterial-like mechanism for promoter opening but not escape. Nat Struct Mol Biol 2005, 12(7):603-607. PMCID or (doi): (10.1038/nsmb949) https://www.ncbi.nlm.nih.gov/pubmed/15937491

83. van Eeuwen T, Shim Y, Kim HJ, Zhao T, Basu S, Garcia BA, Kaplan C, Min J-H, Murakami K: Structure of TFIIH/Rad4-Rad23-Rad33 in damaged DNA opening in Nucleotide Excision Repair. bioRxiv 2021:2021.2001.2014.426755. PMCID or (doi): (10.1101/2021.01.14.426755) http://biorxiv.org/content/early/2021/01/15/2021.01.14.426755.abstract

84. Clapier CR, Iwasa J, Cairns BR, Peterson CL: Mechanisms of action and regulation of ATP-dependent chromatin-remodelling complexes. Nat Rev Mol Cell Biol 2017, 18(7):407-422. PMCID or (doi): (10.1038/nrm.2017.26) https://www.ncbi.nlm.nih.gov/pubmed/28512350

85. Clapier CR, Kasten MM, Parnell TJ, Viswanathan R, Szerlong H, Sirinakis G, Zhang Y, Cairns BR: Regulation of DNA Translocation Efficiency within the Chromatin Remodeler RSC/Sth1 Potentiates Nucleosome Sliding and Ejection. Mol Cell 2016, 62(3):453-461. PMCID or (doi): PMC5291166. https://www.ncbi.nlm.nih.gov/pubmed/27153540

86. Greber BJ, Nguyen THD, Fang J, Afonine PV, Adams PD, Nogales E: The cryo-electron microscopy structure of human transcription factor IIH. Nature 2017, 549(7672):414-417. PMCID or (doi): PMC5844561. https://www.ncbi.nlm.nih.gov/pubmed/28902838

87. Schilbach S, Hantsche M, Tegunov D, Dienemann C, Wigge C, Urlaub H, Cramer P: Structures of transcription pre-initiation complex with TFIIH and Mediator. Nature 2017, 551(7679):204-209. PMCID or (doi): PMC6078178. https://www.ncbi.nlm.nih.gov/pubmed/29088706

88. Aibara S, Schilbach S, Cramer P: Structures of mammalian RNA polymerase II pre-initiation complexes. Nature 2021. PMCID or (doi): (10.1038/s41586-021-03554-8) https://www.ncbi.nlm.nih.gov/pubmed/33902107

89. Vo Ngoc L, Wang YL, Kassavetis GA, Kadonaga JT: The punctilious RNA polymerase II core promoter. Genes Dev 2017, 31(13):1289-1301. PMCID or (doi): PMC5580651. https://www.ncbi.nlm.nih.gov/pubmed/28808065

90. Duttke SH, Lacadie SA, Ibrahim MM, Glass CK, Corcoran DL, Benner C, Heinz S, Kadonaga JT, Ohler U: Perspectives on Unidirectional versus Divergent Transcription. Mol Cell 2015, 60(3):348-349. PMCID or (doi): 4643650. http://www.ncbi.nlm.nih.gov/pubmed/26545075

91. Chen Y, Athma AP, Herudek J, Lubas M, Meola N, Järvelin AI, Andersson R, Pelechano V, Steinmetz LM, Jensen TH, Sendelin A: Principles for RNA metabolism and alternative transcription initiation within closely spaced promoters. Nature Genetics 2016. PMCID or (doi):

92. Kaplan CD: Pairs of promoter pairs in a web of transcription. Nat Genet 2016, 48(9):975-976. PMCID or (doi): (10.1038/ng.3649) http://www.ncbi.nlm.nih.gov/pubmed/27573684

93. Tramantano M, Sun L, Au C, Labuz D, Liu Z, Chou M, Shen C, Luk E: Constitutive turnover of histone H2A.Z at yeast promoters requires the preinitiation complex. Elife 2016, 5. PMCID or (doi): PMC4995100. https://www.ncbi.nlm.nih.gov/pubmed/27438412

94. Klein-Brill A, Joseph-Strauss D, Appleboim A, Friedman N: Dynamics of Chromatin and Transcription during Transient Depletion of the RSC Chromatin Remodeling Complex. Cell Rep 2019, 26(1):279-292 e275. PMCID or (doi): PMC6315372. https://www.ncbi.nlm.nih.gov/pubmed/30605682

95. Lu Z, Lin Z: The origin and evolution of a distinct mechanism of transcription initiation in yeasts. Genome Res 2021, 31(1):51-63. PMCID or (doi): (10.1101/gr.264325.120) https://www.ncbi.nlm.nih.gov/pubmed/33219055

96. Breathnach R, Chambon P: Organization and expression of eucaryotic split genes coding for proteins. Annu Rev Biochem 1981, 50:349-383. PMCID or (doi): (10.1146/annurev.bi.50.070181.002025) https://www.ncbi.nlm.nih.gov/pubmed/6791577

97. Li H, Hou J, Bai L, Hu C, Tong P, Kang Y, Zhao X, Shao Z: Genome-wide analysis of core promoter structures in Schizosaccharomyces pombe with DeepCAGE. RNA Biol 2015, 12(5):525-537. PMCID or (doi): (10.1080/15476286.2015.1022704) http://www.ncbi.nlm.nih.gov/pubmed/25747261

98. Consortium F, the RP, Clst Forrest AR, Kawaji H, Rehli M, Baillie JK, de Hoon MJ, Haberle V, Lassmann T, Kulakovskiy IV, Lizio M, Itoh M, Andersson R, Mungall CJ, Meehan TF, Schmeier S, Bertin N, Jorgensen M, Dimont E, Arner E, Schmidl C, Schaefer U, Medvedeva YA, Plessy C, Vitezic M, Severin J, Semple C, Ishizu Y, Young RS, Francescatto M, Alam I, Albanese D, Altschuler GM, Arakawa T, Archer JA, Arner P, Babina M, Rennie S, Balwierz PJ, Beckhouse AG, Pradhan-Bhatt S, Blake JA, Blumenthal A, Bodega B, Bonetti A, Briggs J, Brombacher F, Burroughs AM, Califano A, Cannistraci CV, Carbajo D, Chen Y, Chierici M, Ciani Y, Clevers HC, Dalla E, Davis CA, Detmar M, Diehl AD, Dohi T, Drablos F, Edge AS, Edinger M, Ekwall K, Endoh M, Enomoto H, Fagiolini M, Fairbairn L, Fang H, Farach-Carson MC, Faulkner GJ, Favorov AV, Fisher ME, Frith MC, Fujita R, Fukuda S, Furlanello C, Furino M, Furusawa J, Geijtenbeek TB, Gibson AP, Gingeras T, Goldowitz D, Gough J, Guhl S, Guler R, Gustincich S, Ha TJ, Hamaguchi M, Hara M, Harbers M, Harshbarger J, Hasegawa A, Hasegawa Y, Hashimoto T, Herlyn M, Hitchens KJ, Ho Sui SJ, Hofmann OM, Hoof I, Hori F, Huminiecki L, Iida K, Ikawa T, Jankovic BR, Jia H, Joshi A, Jurman G, Kaczkowski B, Kai C, Kaida K, Kaiho A, Kajiyama K, Kanamori-Katayama M, Kasianov AS, Kasukawa T, Katayama S, Kato S, Kawaguchi S, Kawamoto H, Kawamura YI, Kawashima T, Kempfle JS, Kenna TJ, Kere J, Khachigian LM, Kitamura T, Klinken SP, Knox AJ, Kojima M, Kojima S, Kondo N, Koseki H, Koyasu S, Krampitz S, Kubosaki A, Kwon AT, Laros JF, Lee W, Lennartsson A, Li K, Lilje B, Lipovich L, Mackay-Sim A, Manabe R, Mar JC, Marchand B, Mathelier A, Mejhert N, Meynert A, Mizuno Y, de Lima Morais DA, Morikawa H, Morimoto M, Moro K, Motakis E, Motohashi H, Mummery CL, Murata M, Nagao-Sato S, Nakachi Y, Nakahara F, Nakamura T, Nakamura Y, Nakazato K, van Nimwegen E, Ninomiya N, Nishiyori H, Noma S, Noma S, Noazaki T, Ogishima S, Ohkura N, Ohimiya H, Ohno H, Ohshima M, Okada-Hatakeyama M, Okazaki Y, Orlando V, Ovchinnikov DA, Pain A, Passier R, Patrikakis M, Persson H, Piazza S, Prendergast JG, Rackham OJ, Ramilowski JA, Rashid M, Ravasi T, Rizzu P, Roncador M, Roy S, Rye MB, Saijyo E, Sajantila A, Saka A, Sakaguchi S, Sakai M, Sato H, Savvi S, Saxena A, Schneider C, Schultes EA, Schulze-Tanzil GG, Schwegmann A, Sengstag T, Sheng G, Shimoji H, Shimoni Y, Shin JW, Simon C, Sugiyama D, Sugiyama T, Suzuki M, Suzuki N, Swoboda RK t Hoen PA, Tagami M, Takahashi N, Takai J, Tanaka H, Tatsukawa H, Tatum Z, Thompson M, Toyodo H, Toyoda T, Valen E, van de Wetering M, van den Berg LM, Verado R, Vijayan D, Vorontsov IE, Wasserman WW, Watanabe S, Wells CA, Winteringham LN, Wolvetang E, Wood EJ, Yamaguchi Y, Yamamoto M, Yoneda M, Yonekura Y, Yoshida S, Zabierowski SE, Zhang PG, Zhao X, Zucchelli S, Summers KM, Suzuki H, Daub CO, Kawai J, Heutink P, Hide W, Freeman TC, Lenhard B, Bajic VB, Taylor MS, Makeev VJ, Sandelin A, Hume DA, Carninci P, Hayashizaki Y: A promoter-level mammalian expression atlas. Nature 2014, 507(7493):462-470. PMCID or (doi): 4529748. http://www.ncbi.nlm.nih.gov/pubmed/24670764

99. Saito TL, Hashimoto S, Gu SG, Morton JJ, Stadler M, Blumenthal T, Fire A, Morishita S: The transcription start site landscape of C. elegans. Genome Res 2013, 23(8):1348-1361. PMCID or (doi): 3730108. http://www.ncbi.nlm.nih.gov/pubmed/23636945

100. Nepal C, Hadzhiev Y, Previti C, Haberle V, Li N, Takahashi H, Suzuki AM, Sheng Y, Abdelhamid RF, Anand S, Gehrig J, Akalin A, Kockx CE, van der Sloot AA, van Ijcken WF, Armant O, Rastegar S, Watson C, Strahle U, Stupka E, Carninci P, Lenhard B, Muller F: Dynamic regulation of the transcription initiation landscape at single nucleotide resolution during vertebrate embryogenesis. Genome Res 2013, 23(11):1938-1950. PMCID or (doi): PMC3814893. https://www.ncbi.nlm.nih.gov/pubmed/24002785

101. Chen RA, Down TA, Stempor P, Chen QB, Egelhofer TA, Hillier LW, Jeffers TE, Ahringer J: The landscape of RNA polymerase II transcription initiation in C. elegans reveals promoter and enhancer architectures. Genome Res 2013, 23(8):1339-1347. PMCID or (doi): 3730107. http://www.ncbi.nlm.nih.gov/pubmed/23550086

102. Yamashita R, Sathira NP, Kanai A, Tanimoto K, Arauchi T, Tanaka Y, Hashimoto S, Sugano S, Nakai K, Suzuki Y: Genome-wide characterization of transcriptional start sites in humans by integrative transcriptome analysis. Genome Res 2011, 21(5):775-789. PMCID or (doi): 3083095. http://www.ncbi.nlm.nih.gov/pubmed/21372179

103. Hoskins RA, Landolin JM, Brown JB, Sandler JE, Takahashi H, Lassmann T, Yu C, Booth BW, Zhang D, Wan KH, Yang L, Boley N, Andrews J, Kaufman TC, Graveley BR, Bickel PJ, Carninci P, Carlson JW, Celniker SE: Genome-wide analysis of promoter architecture in Drosophila melanogaster. Genome Res 2011, 21(2):182-192. PMCID or (doi): 3032922. http://www.ncbi.nlm.nih.gov/pubmed/21177961

104. Kawaji H, Frith MC, Katayama S, Sandelin A, Kai C, Kawai J, Carninci P, Hayashizaki Y: Dynamic usage of transcription start sites within core promoters. Genome Biol 2006, 7(12):R118. PMCID or (doi): PMC1794431. https://www.ncbi.nlm.nih.gov/pubmed/17156492

105. Carninci P, Kasukawa T, Katayama S, Gough J, Frith MC, Maeda N, Oyama R, Ravasi T, Lenhard B, Wells C, Kodzius R, Shimokawa K, Bajic VB, Brenner SE, Batalov S, Forrest AR, Zavolan M, Davis MJ, Wilming LG, Aidinis V, Allen JE, Ambesi-Impiombato A, Apweiler R, Aturaliya RN, Bailey TL, Bansal M, Baxter L, Beisel KW, Bersano T, Bono H, Chalk AM, Chiu KP, Choudhary V, Christoffels A, Clutterbuck DR, Crowe ML, Dalla E, Dalrymple BP, de Bono B, Della Gatta G, di Bernardo D, Down T, Engstrom P, Fagiolini M, Faulkner G, Fletcher CF, Fukushima T, Furuno M, Futaki S, Gariboldi M, Georgii-Hemming P, Gingeras TR, Gojobori T, Green RE, Gustincich S, Harbers M, Hayashi Y, Hensch TK, Hirokawa N, Hill D, Huminiecki L, Iacono M, Ikeo K, Iwama A, Ishikawa T, Jakt M, Kanapin A, Katoh M, Kawasawa Y, Kelso J, Kitamura H, Kitano H, Kollias G, Krishnan SP, Kruger A, Kummerfeld SK, Kurochkin IV, Lareau LF, Lazarevic D, Lipovich L, Liu J, Liuni S, McWilliam S, Madan Babu M, Madera M, Marchionni L, Matsuda H, Matsuzawa S, Miki H, Mignone F, Miyake S, Morris K, Mottagui-Tabar S, Mulder N, Nakano N, Nakauchi H, Ng P, Nilsson R, Nishiguchi S, Nishikawa S, Nori F, Ohara O, Okazaki Y, Orlando V, Pang KC, Pavan WJ, Pavesi G, Pesole G, Petrovsky N, Piazza S, Reed J, Reid JF, Ring BZ, Ringwald M, Rost B, Ruan Y, Salzberg SL, Sandelin A, Schneider C, Schonbach C, Sekiguchi K, Semple CA, Seno S, Sessa L, Sheng Y, Shibata Y, Shimada H, Shimada K, Silva D, Sinclair B, Sperling S, Stupka E, Sugiura K, Sultana R, Takenaka Y, Taki K, Tammoja K, Tan SL, Tang S, Taylor MS, Tegner J, Teichmann SA, Ueda HR, van Nimwegen E, Verardo R, Wei CL, Yagi K, Yamanishi H, Zabarovsky E, Zhu S, Zimmer A, Hide W, Bult C, Grimmond SM, Teasdale RD, Liu ET, Brusic V, Quackenbush J, Wahlestedt C, Mattick JS, Hume DA, Kai C, Sasaki D, Tomaru Y, Fukuda S, Kanamori-Katayama M, Suzuki M, Aoki J, Arakawa T, Iida J, Imamura K, Itoh M, Kato T, Kawaji H, Kawagashira N, Kawashima T, Kojima M, Kondo S, Konno H, Nakano K, Ninomiya N, Nishio T, Okada M, Plessy C, Shibata K, Shiraki T, Suzuki S, Tagami M, Waki K, Watahiki A, Okamura-Oho Y, Suzuki H, Kawai J, Hayashizaki Y, Consortium F, Group RGER, Genome Science G: The transcriptional landscape of the mammalian genome. Science 2005, 309(5740):1559-1563. PMCID or (doi): (10.1126/science.1112014) http://www.ncbi.nlm.nih.gov/pubmed/16141072

106. Kadonaga JT: Perspectives on the RNA polymerase II core promoter. Wiley Interdiscip Rev Dev Biol 2012, 1(1):40-51. PMCID or (doi): 3695423. http://www.ncbi.nlm.nih.gov/pubmed/23801666

107. Haberle V, Stark A: Eukaryotic core promoters and the functional basis of transcription initiation. Nat Rev Mol Cell Biol 2018, 19(10):621-637. PMCID or (doi): PMC6205604. https://www.ncbi.nlm.nih.gov/pubmed/29946135

108. Luse DS, Parida M, Spector BM, Nilson KA, Price DH: A unified view of the sequence and functional organization of the human RNA polymerase II promoter. Nucleic Acids Res 2020, 48(14):7767-7785. PMCID or (doi): PMC7641323. https://www.ncbi.nlm.nih.gov/pubmed/32597978

109. Chou S-P, Alexander AK, Rice EJ, Choate LA, Cohen PE, Danko CG: Genetic Dissection of the RNA Polymerase II Transcription Cycle. bioRxiv 2021:2021.2005.2023.445279. PMCID or (doi): (10.1101/2021.05.23.445279) http://biorxiv.org/content/early/2021/05/30/2021.05.23.445279.abstract

110. Winston F, Dollard C, Ricupero-Hovasse SL: Construction of a set of convenient Saccharomyces cerevisiae strains that are isogenic to S288C. Yeast 1995, 11(1):53-55. PMCID or (doi): (10.1002/yea.320110107) http://www.ncbi.nlm.nih.gov/pubmed/7762301

111. Amberg DC, Burke DJ, Strathern JN: Methods in Yeast Genetics: A Cold Spring Harbor Laboratory Course Manual, 2005 Edition, 2005 Edition edn. Cold Spring Harbor, NY: Cold Spring Harbor Press; 2005

112. Ranish JA, Hahn S: The yeast general transcription factor TFIIA is composed of two polypeptide subunits. The Journal of biological chemistry 1991, 266(29):19320-19327. PMCID or (doi): http://www.ncbi.nlm.nih.gov/pubmed/1918049

113. Schmitt ME, Brown TA, Trumpower BL: A rapid and simple method for preparation of RNA from Saccharomyces cerevisiae. Nucleic acids research 1990, 18(10):3091-3092. PMCID or (doi): 330876. http://www.ncbi.nlm.nih.gov/pubmed/2190191

114. Martin M: Cutadapt removes adapter sequences from high-throughput sequencing reads. EMBnetjournal; Vol 17, No 1: Next Generation Sequencing Data AnalysisDO - 1014806/ej171200 2011. PMCID or (doi): https://journal.embnet.org/index.php/embnetjournal/article/view/200/479

115. Langmead B, Trapnell C, Pop M, Salzberg SL: Ultrafast and memory-efficient alignment of short DNA sequences to the human genome. Genome Biol 2009, 10(3):R25. PMCID or (doi): PMC2690996. https://www.ncbi.nlm.nih.gov/pubmed/19261174

116. Li H, Handsaker B, Wysoker A, Fennell T, Ruan J, Homer N, Marth G, Abecasis G, Durbin R, Genome Project Data Processing S: The Sequence Alignment/Map format and SAMtools. Bioinformatics 2009, 25(16):2078-2079. PMCID or (doi): PMC2723002. https://www.ncbi.nlm.nih.gov/pubmed/19505943

117. Schwalb B, Tresch A, Torkler P, Duemcke S, Demel C, Ripley B, Venables B: LSD: Lots of Superior Depictions. In.; 2011.

118. Li H: Aligning sequence reads, clone sequences and assembly contigs with BWA-MEM. arXiv 2013, 1303.3997. PMCID or (doi):

119. Rossi MJ, Lai WKM, Pugh BF: Simplified ChIP-exo assays. Nat Commun 2018, 9(1):2842. PMCID or (doi): PMC6054642. https://www.ncbi.nlm.nih.gov/pubmed/30030442

